# Live imaging of breast tumors shows macrophage-dependent induction and TMEM-mediated enrichment of cancer stem cells during metastatic dissemination

**DOI:** 10.1101/2020.09.18.303388

**Authors:** Ved P Sharma, Binwu Tang, Yarong Wang, George S Karagiannis, Emily A Xue, David Entenberg, Lucia Borriello, Anouchka Coste, Camille L Duran, Robert J Eddy, Gina Kim, Xianjun Ye, Joan G Jones, Eli Grunblatt, Nathan Agi, Sweta Roy, Gargi Bandyopadhyaya, Esther Adler, Chinmay R Surve, Dominic Esposito, Sumanta Goswami, Wenjun Guo, John S Condeelis, Lalage M. Wakefield, Maja H Oktay

**Affiliations:** Department of Anatomy and Structural Biology, Albert Einstein College of Medicine, Bronx, NY, USA; Gruss-Lipper Biophotonics Center, Albert Einstein College of Medicine, Bronx, NY, USA; Integrated Imaging Program, Albert Einstein College of Medicine, Bronx, NY, USA; Laboratory of Cancer Biology and Genetics, National Cancer Institute, Bethesda, MD USA; Protein Expression Laboratory, Frederick National Laboratory for Cancer Research, Frederick MD, USA; Department of Pathology, Albert Einstein College of Medicine, Bronx, NY, USA; Department of Surgery, Albert Einstein College of Medicine, Bronx, NY, USA; Department of Epidemiology and Population Health, Albert Einstein College of Medicine, Bronx, NY, USA; Department of Biology, Yeshiva University, New York, NY, USA; Department of Pathology, NYU Langone Medical Center, NY, USA; Department of Cell Biology, Albert Einstein College of Medicine, Bronx, NY, USA; Ruth L. and David S. Gottesman Institute for Stem Cell and Regenerative Medicine Research, Albert Einstein College of Medicine, Bronx, NY, USA

**Author notes:** Co-senior and corresponding authors.

## Abstract

Cancer stem cells (CSCs) play an important role during metastasis, but the dynamic behavior and induction mechanisms of CSCs are not well understood. We employed high-resolution intravital microscopy using a CSC biosensor to directly observe CSCs in live mice with mammary tumors. CSCs display the slow-migratory, invadopod-rich phenotype that is the hallmark of disseminating tumor cells. CSCs are enriched near macrophages, particularly near macrophage-containing intravasation sites called Tumor Microenvironment of Metastasis (TMEM) doorways. A dramatic enrichment of CSCs occurs on association with TMEM doorways, contributing to the finding that CSCs represent ∼>60% of circulating tumor cells. Mechanistically, stemness is induced in non-stem cancer cells upon their direct contact with macrophages via Notch signaling. In breast cancers from patients, the density of TMEM doorways correlates strongly with the proportion of cancer cells expressing stem cell markers, indicating that in human breast cancer TMEM doorways are not only cancer cell intravasation portals but also CSC programming sites.

**One Sentence Summary:** Intravital imaging reveals macrophage-mediated induction of cancer stem cells in vivo and their dramatic enrichment on dissemination through TMEM doorways.

## INTRODUCTION

A high proportion of cancer stem cells (CSCs) in primary tumors is associated with poor prognosis and increased metastatic relapse (1). Metastasis, which accounts for >90% of cancer-related deaths, is a multi-step process involving cancer cell intravasation into the bloodstream, dissemination to distant sites, and seeding in new organs (2). A conceptual advance in understanding of this complex process came from realization that breast tumors are hierarchically organized with respect to individual cell proliferative potential and differentiation status (3). At the apex of the hierarchy are the CSCs. CSCs have enhanced ability to self-renew which makes them uniquely capable of initiating and sustaining primary and metastatic tumor growth (4). Importantly CSCs are intrinsically more therapy-resistant than their more differentiated progeny (5). Thus, a better understanding of the *in vivo* properties of CSCs is essential for the development of more effective anti-cancer therapies.

Cancer progression is profoundly influenced by complex dynamic and reciprocal interactions between the tumor cells and the other cellular and acellular components of the tumor microenvironment (6, 7). Various cell types, as well as extracellular matrix, surrounding CSCs contribute to the microenvironmental niche that nurtures and sustains the CSC phenotype and influences its subsequent fate (8). Indeed, the emerging concepts in CSC biology argue against a stable CSC phenotype which could be reliably tested *in vitro* because it is now understood that the retention of cancer cells in the phenotypic “stem” state in solid tumors can depend on signals from tumor microenvironment (9). Moreover, certain microenvironmental signals can induce phenotypic plasticity and allow non-stem cancer cells to re-acquire a stem-like phenotype (10). This complexity is challenging to model adequately *in vitro*, and existing approaches to understanding CSC biology *in vivo*, while informative, have suffered from a number of limitations. CSCs are commonly identified by tumor excision and digestion to single cells, followed by flow cytometry for previously validated combinations of cell-surface markers (4), a process that results in loss of both spatial and dynamic information. Lineage tracing approaches in genetically-engineered animals retain spatial context and have yielded useful insights, such as the multipotent vs. unipotent nature of the mammary stem cell during development (5) and the origin and fate of cancer stem cells in the intestine (11, 12). However, cell fate trajectories are generally inferred from sequential static images, and cause and effect relationships as well as reversible phenomena cannot be readily observed.

A complementary approach that preserves both positional and dynamic information is to use intravital imaging coupled with fluorescent reporters that are transiently switched on in specific phenotypic states. To this end, we used an extensively validated lentiviral-based fluorescent reporter that is activated by binding of the stem cell master transcription factors Sox2 and Oct4, or their paralogs, to an artificial enhancer element (SORE6) (13). This reporter specifically identifies tumor cells with the expected properties of cancer stem cell (13). Since the fluorescent protein is destabilized, the acquisition and loss of the stem-like phenotype can be monitored with good kinetic resolution and high selectivity, which is critical to capture the plasticity of cancer stem cells (9). Coupled with single-cell resolution multiphoton microscopy in live animals, this sensor allowed us to address dynamic properties of the cancer stem cell in real-time in the living tumors.

Here we use intravital imaging at single cell resolution and longitudinally during tumor progression to show for the first time the phenotype of individual CSCs *in vivo* during dissemination and seeding. We identify the mechanism that links the induction of stemness to form CSCs, intravasation and dissemination, and investigate the extraordinary enrichment of CSCs during progression of metastasis. Furthermore, we explore whether similar relationships between tumor cell dissemination and density of CSCs occur in breast cancer patients.

## RESULTS

### 1. Breast CSCs are a minority population in vivo in primary tumors

To characterize the CSC population *in vivo* in the living animal and *in situ* in tissue, we used our previously-validated “SORE6” CSC biosensor (13), with minor modifications to enable detection in FFPE tissue (see Fig. S1A and Methods). In this sensor (“SORE6>GFP”), six repeats of a composite SOX2/OCT4 response element coupled to a minimal CMV promoter (minCMV) drives expression of a destabilized fluorescent protein following activation by the CSC master transcription factors, SOX2 and OCT4, or their paralogs. A parallel construct (“minCMV>GFP”) lacking the SOX2/OCT4 response elements was used as a control for FACS gating and setting microscopy detection thresholds. Using an *in vivo* limiting dilution assay, we confirmed that SORE6+ cells are significantly enriched (7.8-fold) for tumor-initiating activity in the MDA-MB-231 subline used here (Fig S1B). This level of enrichment is similar to the enrichment (9.5-fold) that we previously found using the original sensor in MDA-MB-231 cells, and confirmed by 5 passages of serial transplantation (13). It is also comparable to the enrichment found by others (8.5-fold) using a different (integrin-based) sorting strategy to identify CSCs in the MDA-MB231 model (14). Moreover, we demonstrated greatly enhanced chemoresistance in SORE6 + cells (Figs S1C, D), a property associated with the stem phenotype (5), as well as increased expression of canonical stem cell transcription factors (Fig. S1E). Importantly, we also found that SORE6+ (compared to SORE6-) cells express significantly higher levels of the transcription factor Snail1 which is involved in epithelial to mesenchymal transition (EMT). This indicates that SORE6+ cells have also activated an embryonic program critical for invasion and metastatic dissemination (15–17).

To address CSC representation in primary tumors, orthotopic xenograft breast tumors were generated from MDA-MB-231 cells expressing the SORE6>GFP CSC sensor and a constitutive tdTomato volume marker of all tumor cells. Using fixed-frozen tumor imaging, we found that the SORE6+ cells (i.e. GFP+ cells) comprised a minority population in primary tumors compared to all tumor cells (Fig 1A). FACS analysis of primary tumors also showed that the CSC population was a minority population of cancer cells (Fig S1F). We also investigated the representation of SORE6+ CSCs in formalin-fixed, paraffin-embedded (FFPE) xenograft tissues. Since FFPE destroys the fluorescent signal (dsCopGFP and tdTomato), we immunostained for the FLAG tag on the SORE6 biosensor (see Fig S2A, S2B for FLAG antibody characterization *in vitro* and *in vivo*, respectively). In SCID mouse hosts, tumor tissue expressing the SORE6 construct showed a small population of FLAG+ single CSCs distributed throughout the tumor tissue (Fig 1B) which were not seen in tumor tissue expressing the minCMV>GFP control construct. Similar results were seen in an independent experiment in nude mouse hosts (Fig S2C).

**Figure 1:**
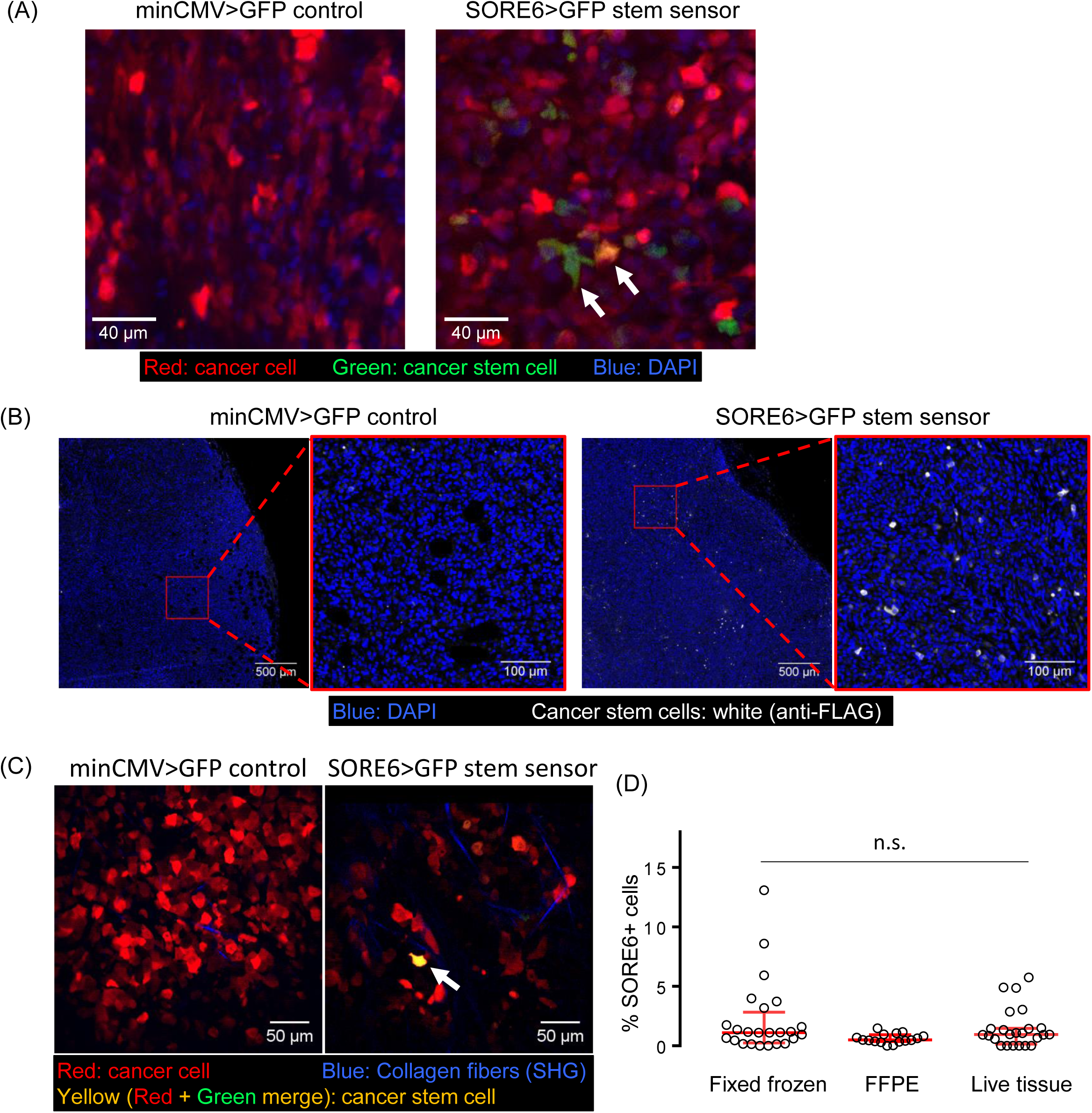
Breast carcinoma stem cells are a minority population in the primary tumor *in vivo*. (A) Representative images of fixed frozen tumor tissues. Left panel: tdTomato-minCMV>GFP vector control tumor tissue, right panel: tdTomato-SORE6>GFP reporter MDA-MB-231 tumor tissue. Arrows point to two examples of GFP+/SORE6+ stem cells, which appear green to yellow due to varying levels of tdTomato (volume marker) intensity in the cell. (B) *In situ* immunofluorescence staining for the FLAG tag (white) on the GFP reporter in FFPE tumor tissues identifies SORE6+ stem cells as a minority population in fixed tissues in SCID host. Two left panels: tdTomato-MDA-MB-231–minCMV>GFP vector control tumor tissue; two right panels: tdTomato-MDA-MB-231-SORE6>GFP reporter tumor tissue. Nuclei were stained with DAPI (blue). (C) Intravital multiphoton microscopy identifies SORE6+ stem cells as a minority population (white arrow) in living mouse. Left panel: tdTomato-MDA-MB-231 –minCMV>GFP vector control tumor tissue, right panel: tdTomato-MDA-MB-231-SORE6>GFP reporter tumor tissue. (D) Quantification of % SORE6+ stem cells in breast tissue using the three different methodologies described in A, B, and C. n=24 fields of 330×330 μm^2^ from 3 mice for Fixed-frozen tissues; n=16 fields of 500×500 μm^2^ from 3 mice for FFPE tissues; n=24 fields of 340×340 μm^2^ from 3 mice for live tissue; Scatter plot showing median ± interquartile range, one-way ANOVA n.s. p-value=0.09.

To visualize SORE6+ CSCs in living tissue in mice, we imaged primary tumors expressing the SORE6 construct using intravital imaging through implantable mammary imaging windows (see materials and methods for details). The tumor expressing control vector (minCMV>GFP) was used to determine the background GFP signal (Fig S3). Live intravital imaging confirmed the presence of GFP+ CSCs in tumors with tumor cells harboring the SORE6>GFP sensor, compared to tumors harboring minCMV>GFP control (Fig 1C). We quantified SORE6+ CSCs in fixed and live tissue using all three different methodologies (fixed-frozen, FFPE, and live intravital imaging) described above and found that *in vivo* SORE6+ CSCs are on average between 0.7-2.5% of the total tumor cells in the primary tumor, constituting a minority population (Fig 1D), as predicted by the cancer stem cell hypothesis (4).

### 2. Breast CSCs exhibit the slow migratory phenotype and contain invadopodia

A major advantage of the live imaging approach is that it allows visualization of dynamic features of the CSC population. To investigate the migratory phenotypes of CSC in the live primary tumor, SORE6+ CSCs were imaged *in vivo* using time-lapse intravital multiphoton microscopy. In MDA-MB-231 cell xenografts, we observed two migratory phenotypes based on locomotion speed of tumor cell; the fast (migrating rapidly toward blood vessels) and slow (perivascular, invadopodium-rich, invasive) phenotypes, as documented previously (18–20). Among motile cells, we found that the high-speed migration phenotype was more characteristic of SORE6-(non-CSCs) cells, whereas SORE6+ CSCs showed the slow-migrating phenotype (Fig 2A, S4A, Movies 1 and 2). We quantified single tumor cell speeds and found that compared to non-CSCs, which moved at high speeds of ∼1 μm/min, CSCs moved approximately five-times slower (∼ 0.2 μm/min) (Fig. 2B).

**Figure 2:**
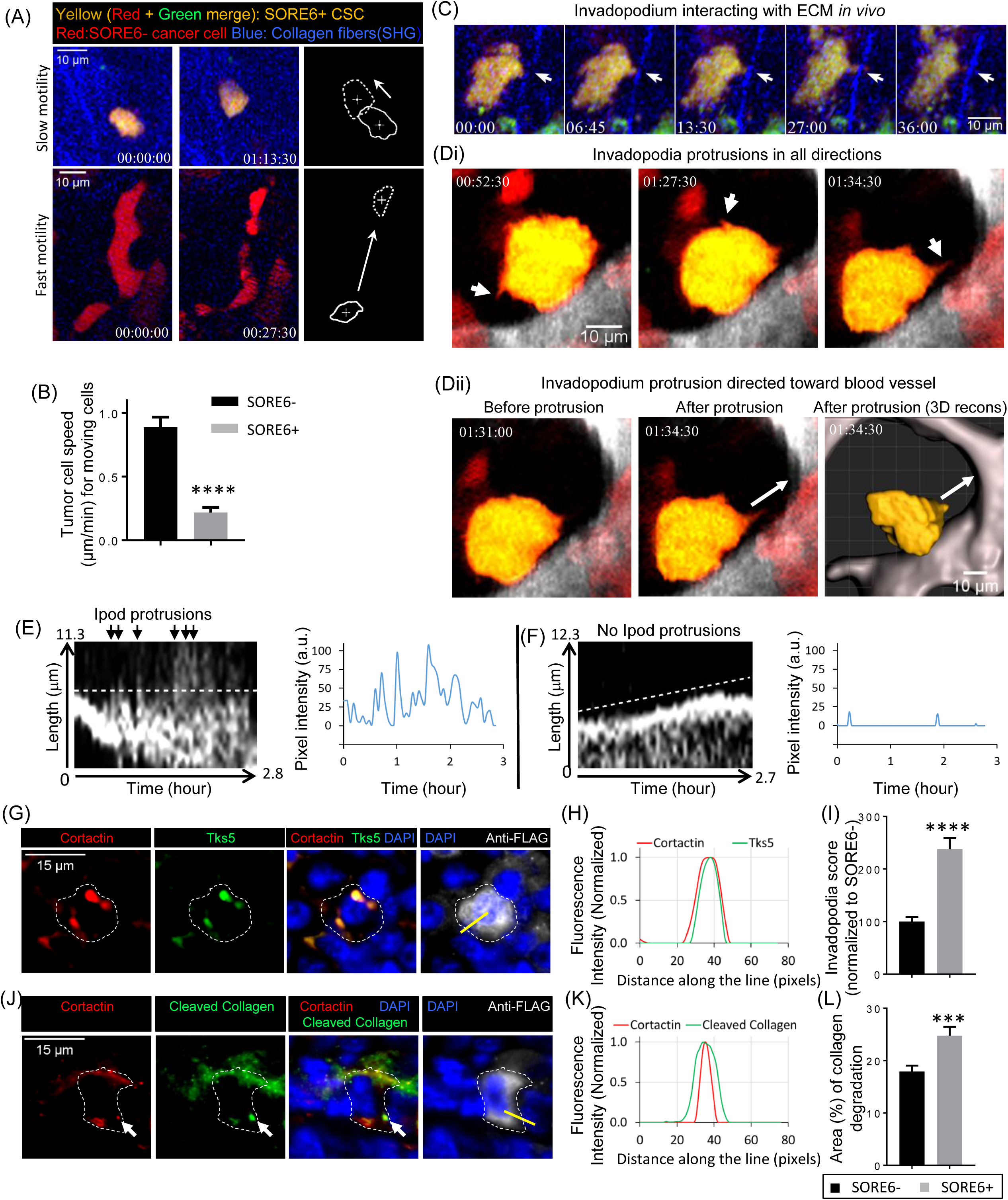
Breast carcinoma stem cells exhibit the slow migratory, invadopodium rich phenotype in vivo. (A) Top panels show first and last frames of intravital movie 1 depicting slow motility phenotype of CSC. Bottom panels show first and last frames of intravital Movie 2 depicting fast motility phenotype of non-CSC. Rightmost panels show cell outlines in first (solid) and last (dotted) frames of the movies with crosses marking cell centroids, and show slow and fast motility phenotypes of CSC and non-CSC, respectively, over the time intervals indicated. White arrows point in the direction of cell movement. Time in hour:min:sec. (B) Quantification of cell speeds (μm/min) for non-stem (SORE6-) and stem cells (SORE6+). n=25 SORE6-, 32 SORE6+ cells in 3 mice; plot showing mean ± SEM, unpaired Mann-Whitney test, **** p-value<0.0001. (C) High-resolution single cell view of SORE6+ stem cell with time-lapse panels, from Movie 3, showing dynamic invadopodial protrusion (white arrows) interacting with ECM fiber seen by second harmonic imaging (blue) in vivo. Time in min:sec. (D) 
i. High-resolution single cell view of a SORE6+ stem cell (yellow) with time-lapse panels, from Movie 4, showing dynamic invadopodial protrusions (white arrows) coming out in all directions, including the one directed toward the blood vessel (rightmost panel).
ii. Two left most panels show CSC before and after invadopodium protrusion directed toward the blood vessel (white arrow). The rightmost panel, from movie 5, shows 3D reconstruction view of CSC with invadopodium protrusion directed toward the blood vessel (white arrow in direction of protrusion). Blood vessels are contrasted grey based on IV injection of Far-red Q dots. (E) Left: kymograph showing the periodicity of invadopodial (ipod) protrusions with oscillations characteristic of active invadopodia (black arrows). Right: Intensity profile through the kymograph (white dotted line), with 17 protrusion peaks during the 2.8-hour long imaging, indicating ∼ 10 min protrusion periodicity characteristic of invadopodial protrusions. (F) Left: kymograph of a SORE6-non-stem cell leading edge showing no periodic protrusions. Right: intensity profile through the kymograph (white dotted line) for non-stem cell showing no invadopodial protrusions. (G) In situ immunofluorescence staining shows that stem cells (anti-FLAG) contain invadopodial protrusions as identified with invadopodial core markers (anti-Cortactin + anti-Tks5). White dotted line marks the boundary of the stem cell and identifies that cortactin+Tks5 positive invadopodia belong to this stem cell. (H) Cortactin and Tks5 fluorescence intensity plots along the yellow line shown in G. (I) Invadopodia score quantification (invadopodia identified as Cortactin/Tks5 co-localized invadopodial cores) in non-stem and SORE6+ stem cells using an automated 36μm ROI scoring method shows that SORE6+ stem cells are enriched in invadopodial protrusions. n=20 SORE6 primary tumor fields at 40X magnification from 4 mice; showing mean ± SEM, Wilcoxon test, **** p-value<0.0001. (J) In situ immunofluorescence staining shows that stem cell (anti-FLAG) invadopodia have degradation activity (anti-Col3/4) associated with them. White arrows indicate the co-localization of freshly cleaved collagen with invadopodia. White dotted line marks the boundary of the stem cell. The position of the cleaved collagen does not always co-localize with invadopodia, This is a well-known issue with cleaved collagen (22). The reason is that cleaved collagen signal represents the current as well as historical buildup of degraded collagen as different invadopodia at different times degrade different areas of ECM and move on. (K) Cortactin and Cleaved collagen fluorescence intensity plots along the yellow line shown in J. (L) Quantification of collagen degradation area in non-stem and stem cell areas using an automated 36μm ROI scoring method. n=16 SORE6 primary tumor fields at 40X magnification from 4 mice; showing mean ± SEM, paired two-tailed Student’s t test, *** p-value<0.0001.

In the *in vitro* setting, invadopodia (invasive protrusions) form only on the ventral surface of the cell, since that is where the cell contacts the ECM. In contrast, in the 3D tumor microenvironment *in vivo*, cells are surrounded by ECM and invadopodia are seen extending in all directions (19, 21–23). To investigate if CSCs make invadopodia, we imaged SORE6+ CSCs at high resolution *in vivo*. We observed that CSCs make oscillatory cellular protrusions directed toward ECM fibers (Fig 2C, Movie 3) and blood vessels (Fig 2D, Movies 4 and 5), similar to the previously described oscillatory invadopodia associated with the slow-migratory phenotype of disseminating tumor cells (19, 24, 25). Kymograph analysis and the line profile through the protrusions showed the highly dynamic oscillatory nature of invasive invadopodial protrusions with an 8-10 min period (Fig 2E), in agreement with previously published values (21, 24). In contrast, non-CSCs, whether close to blood vessels or away from them, did not show oscillatory invadopodial protrusions (Fig 2F, S4B, S4C, Movies 6 and 7).

To determine if the CSC protrusions seen *in vivo* are indeed invadopodia with the canonical cortactin-Tks5 core and ECM degrading activity (19, 21, 26, 27), fixed tissue sections were stained with FLAG antibody to identify CSCs, together with cortactin and Tks5 antibodies. We found that both cortactin and Tks5 co-localized to protrusions on SORE6+ CSCs (Fig 2G, H), indicating these CSC protrusions are invadopodia. We counted the number of invadopodia in stem cells and non-stem cell areas using both an automated and manual method (see materials and methods), and found that compared to non-stem cells, stem cells contain significantly higher numbers of invadopodia (Fig 2I, S4E). To check whether CSC invadopodia have ECM degrading activity, we utilized a cleaved-collagen antibody which detects degraded collagen in vivo (22). Triple IF staining with cortactin, cleaved-collagen and FLAG antibodies showed co-localization of cortactin positive-protrusions in stem cells with degraded collagen *in vivo* (Fig 2J, K). Quantification of degraded collagen showed that CSCs have higher degradation activity than non-stem cells (Figs 2L, S4F).

To further assess the matrix degradation potential of stem cells versus non-stem cells, we performed an *in vitro* invadopodium-dependent degradation assay (21). SORE6+ CSCs or SORE6-non-CSCs were plated separately on fluorescent gelatin overnight and stained for invadopodial markers, cortactin and Tks5. We found cortactin and Tks5 positive invadopodia actively degrading the underlying matrix (Fig S4G). Degradation area quantification showed that CSCs have higher matrix degrading activity compared to the non-CSCs (Fig S4H). Our *in vivo* and *in vitro* results together indicate that CSCs have increased numbers of invadopodia with extracellular matrix degradation activity, a key property needed for the transendothelial migration step of intravasation during the dissemination of tumor cells in the metastatic cascade (26).

### 3. Macrophages regulate stemness in breast cancer cells

Previous studies have found that macrophages can enhance breast tumorigenesis (28), are associated with prometastatic changes during tumor progression including increased metastatic seeding (29, 30), and promote tumor cell intravasation upon direct contact with tumor cells (20, 31, 32). Therefore, we investigated the relative spatial distribution of CSCs and macrophages by intravital imaging of SORE6+ CSCs in primary mammary tumors in mice with CFP expressing macrophages (MacBlue host mice) (Fig 3A). We found in live tumors that 60-70% of the CSCs are in direct physical contact with a macrophage (Fig 3C-live). We confirmed our findings in fixed-frozen tumor tissue, where we identified macrophages with an anti-Iba1 antibody (Fig 3B), and again found that there are more than two times more CSCs in direct physical contact with a macrophage than not (Fig 3C - fixed).

**Figure 3:**
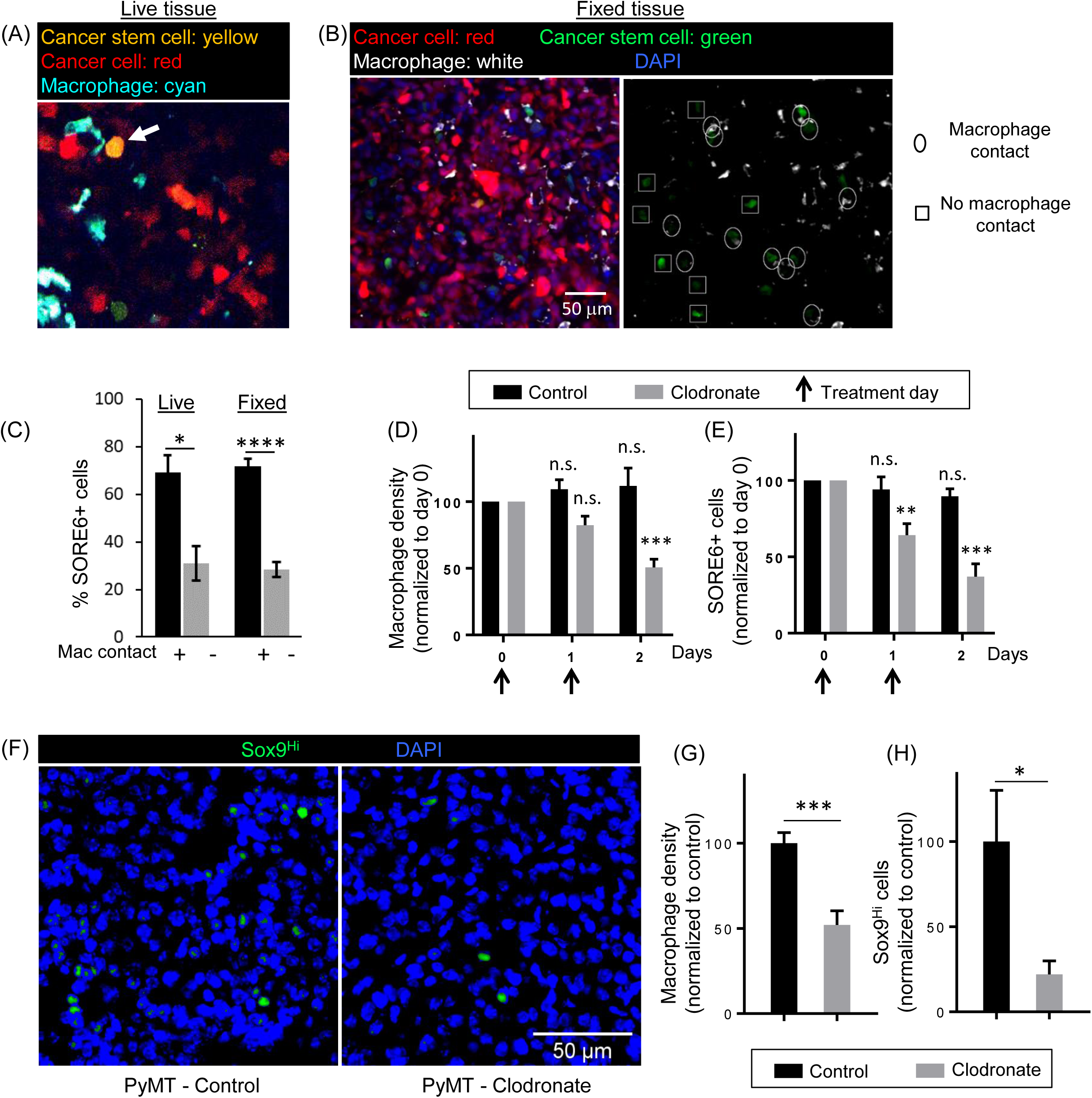
Macrophages regulate stemness in breast cancer cells. (A) Still intravital microscopy image of CSCs (SORE6+) in primary tumor, showing a CSC (yellow) in contact with a macrophage (cyan) in living mouse (B) Representative images of fixed frozen tumor tissues stained with IBA-1 antibody for macrophages in primary tumor. Left panel shows cancer cell (tdTomato in red), cancer stem cell (SORE6+ in green), macrophage (IBA-1 in white) and nuclei (DAPI in blue). Right panel, same as left but lacking tdTomato and DAPI channels, is shown for a better view of cancer stem cells in contact with macrophages (highlighted with an oval), or not in contact with macrophage (highlighted with a square). (C) Quantification of % SORE6+ cells in direct contact with macrophages or not in live tissue and fixed frozen primary tumors. Live: n=17 fields of 340×340 μm^2^ in 4 mice; Fixed: n=24 fields of 330×330 μm^2^ in 3 mice; plot showing mean ± SEM, Wilcoxon test, Live: * p-value=0.0195, Fixed: **** p-value<0.0001. (D) Quantification of changes in macrophage density in live mammary tumor over time after control PBS liposome or clodronate liposome treatment. Vertical arrows indicate the treatment days (day 0 and 1). n=7 fields of 512×512 μm^2^ each for control and clodronate treatments. Data normalized to macrophage density value at day 0 for each treatment and plotted as mean ± SEM, Tukey’s multiple comparisons test; p-values: day0 vs day1(control) 0.44, day0 vs day2(control) 0.67, day0 vs day1(clodronate) 0.09, day0 vs day1(clodronate) 0.0005. (E) Quantification of changes in SORE6+ cells in live mammary tumor over time after clodronate treatment to deplete macrophages. Vertical arrows indicate the treatment days (day 0 and 1). n=7 fields of 512×512 μm^2^ each for control and clodronate treatments. Data normalized to SORE6+ cells value at day 0 for each treatment and plotted as mean ± SEM, Tukey’s multiple comparisons test; p-values: day0 vs day1(control) 0.76, day0 vs day2(control) 0.16, day0 vs day1(clodronate) 0.007, day0 vs day2(clodronate) 0.0007. (F) Images of PyMT primary tumor tissues treated with either PBS liposomes (control) or clodronate liposomes and stained with Sox9 antibody (green) and DAPI (blue). Image shows Sox9^Hi^ cells. (G) Quantification of changes in macrophage density in fixed PyMT mammary tumor after PBS (control) or clodronate liposomes treatment. n= 11 (control) and 6 (clodronate) fields of 2-3 mm^2^ tumor tissue from 5 mice (control) and 4 mice (clodronate); data normalized to macrophage density value for control treatment and plotted as mean ± SEM, Mann-Whitney test, *** p-value=0.0002. (H) Quantification of changes in Sox9^Hi^ cells in fixed PyMT mammary tumor after PBS (control) or clodronate liposomes treatment. n= 17 (control) and 11 (clodronate) fields of 1000×500 μm^2^ tumor tissue from 5 mice (control) and 4 mice (clodronate); data normalized to Sox9^Hi^ cells value for control treatment and plotted as mean ± SEM, Mann-Whitney test, * p-value=0.0221.

To evaluate whether this interaction between macrophages and CSCs was causally associated with stemness *in vivo*, we depleted macrophages in tumor tissues with clodronate treatment (18). *In vivo* intravital imaging with micro-cartography-based localization (33) was performed to image the same tumor fields over 2 days in mice bearing primary mammary tumors expressing the SORE6 sensor with daily clodronate treatment. Clodronate treatment led to significant macrophage depletion in tumor tissue over 2 days (Fig 3D), and this was associated with significant reduction in CSCs *in vivo* during the same time period (Fig 3E), suggesting the macrophages contribute to CSC niche.

Furthermore, we extended the role of macrophages in stemness regulation beyond the MDA-MB-231 human xenograft model to an autochthonous breast cancer model that fully recapitulates the entire cancer progression process, by using the MMTV-PyMT transgenic mouse (“PyMT”) (34). Since the lentiviral SORE6 stemness sensor cannot be used directly in this transgenic model, we sought a surrogate marker of stemness. Met-1 cells, which are derived from a PyMT tumor (35), were transduced with the SORE6 and minCMV control sensors, and FACS analysis showed that the SORE6 sensor identifies a minority stem cell population (∼ 8%) in these cultures (Fig S5A), similar to results obtained with MDA-MB-231 cells (Fig S1F). Although Sox2 mRNA was not detectable in these cells (not shown), expression of Sox9, a Sox2 paralog, was enriched 8-fold in SORE6+ Met-1 cells when compared with SORE6-cells (Fig S5B). As Sox9 has previously been implicated in the maintenance of the mammary stem cell state (36, 37), and was consistently associated with stem cell features in a panel of breast cancer cell lines (38), we proceeded to use Sox9 expression as a surrogate marker of stemness in the PyMT model. PyMT mice bearing primary tumors were treated with either control PBS or clodronate liposomes, and macrophage (Iba-1 IHC, brown staining) coverage area was quantified. We found approximately 50% decrease in macrophage density after clodronate treatment (Fig 3G), similar to results obtained in MDA-MB-231 primary tumor model (Fig 3D). To check the effect of clodronate treatment on Sox9Hi stem cells, PyMT tissues were stained with Sox9 antibody (Fig 3F) and Sox9^Hi^ tissue coverage area was quantified. We found significant reductions (approximately 80%) in Sox9^Hi^ cells after clodronate treatment (Fig 3H), similar to results obtained in MDA-MB-231 primary tumor model (Fig 3E). Thus, in two independent models, we observed a reduction in macrophages to be associated with a concomitant reduction in CSCs.

### 4. Macrophage contact induces stemness in breast cancer cells through Notch1 signaling

The functional link between CSCs and macrophages observed above could result from macrophage-regulated expansion of a pre-existing CSC population, or it could reflect induction de novo of the CSC state in non-CSCs. To distinguish these possibilities, we stained control and clodronate-treated PyMT primary tumor tissues with Ki67 (proliferation marker) and found that macrophage depletion did not affect CSC proliferation in vivo (supplementary Fig S6A), indicating that the primary mechanism of macrophage-mediated tumor cell stemness does not involve proliferative expansion of pre-existing CSCs.

To investigate contributing mechanisms further, we performed intravital time-lapse imaging of tumors to observe the response of individual tumor cells to macrophage interactions. Using the SORE6 biosensor, we found examples of non-stem tumor cells converting to stem cells upon macrophage contact *in vivo* (Fig 4A upper panels, Movies 9 and 10), with the stemness biosensor signal rising significantly above background by ∼ 0.5 hours after first contact with a macrophage (Fig 4A), and on average by 1.5 hours for the average of the tumor cell population (Fig. 4B). In contrast, tumor cells which were not contacted by a macrophage, did not show stemness induction (Fig 4A, lower panels).

**Figure 4:**
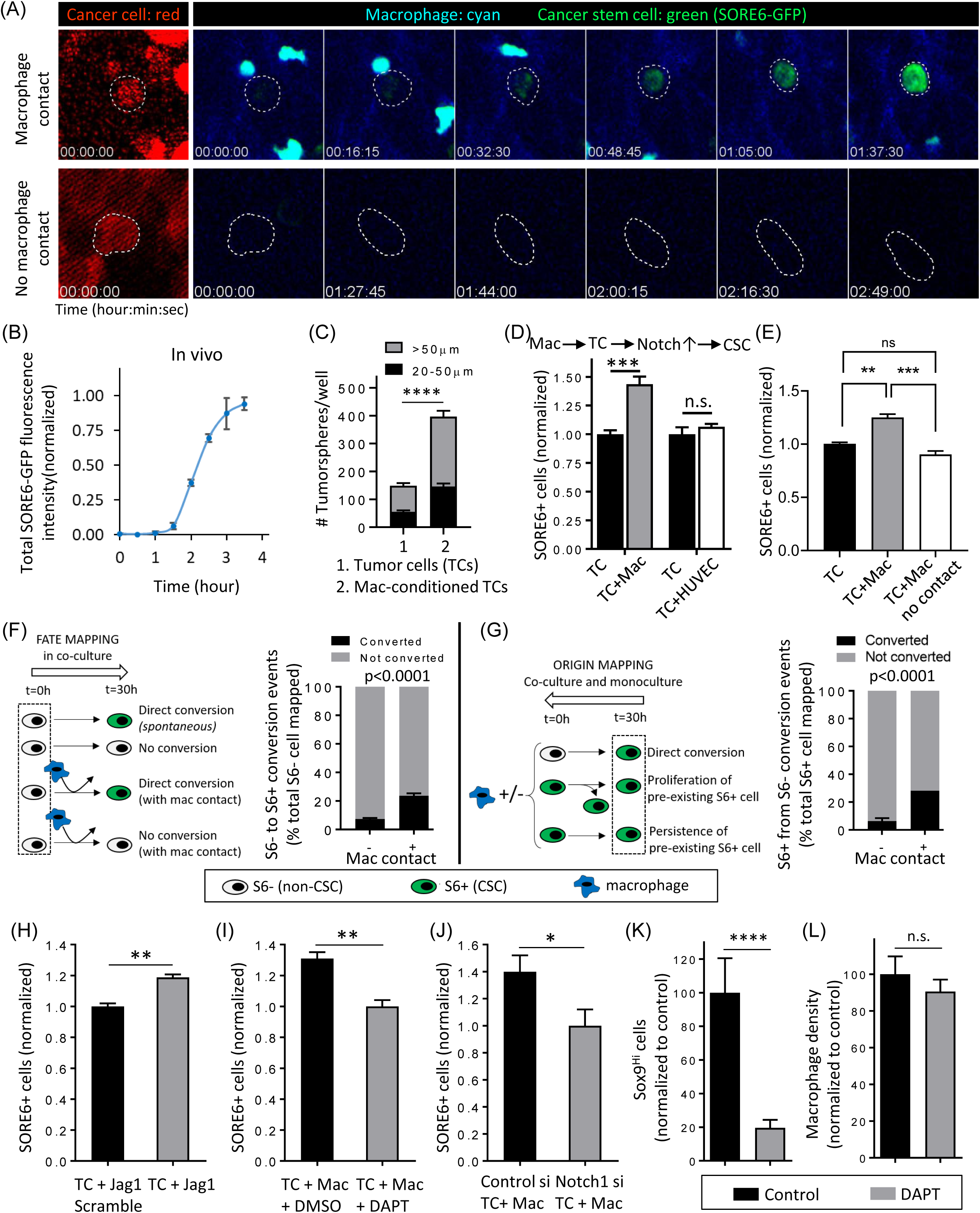
Macrophage contact induces stemness in breast cancer cells through Notch1 signaling. (A) Upper panels are time-lapse images from intravital Movie 9, showing SORE6>GFP induction (green) in a tumor cell after macrophage (cyan) contact. White dotted lines show cancer cell outline before and after stemness induction and is drawn based on the tdTomato volume marker (red channel, left panel). Lower panels show lack of stemness induction (no SORE6-GFP (green) increase) in a tumor cell, which is not contacted by a macrophage. White dotted lines show tumor cell outline during the time course and is drawn based on the tdTomato volume marker (red channel, left panel). (B) Quantification of total SORE6-GFP fluorescence intensity during stemness induction in tumor cells in vivo. Time 0 corresponds to the time when macrophage contacts the tumor cell. n>200 cells analyzed from 10 mice, to identify tumor cells contacting macrophages and showing induced stemness, each data point plotted as mean ± SEM. Data normalized to SORE6-GFP value before macrophage contact to set the 0 background and max SORE6-GFP value as 1 after macrophage contact. (C) Tumorsphere forming activity of MDA-MB-231 cells with or without prior co-culture with BAC cells for 24 hours. Results are mean ± SEM for n=6 (tumor cells), n=12 (BAC-conditioned tumor cells). Unpaired t-tests for each size class, **** p<0.0001. (D) In vitro co-culture assay of tumor cells with or without macrophages or endothelial cells show that contact with macrophages, and not with endothelial cells, induces stemness in tumor cells. Tumor cells were co-cultured with macrophages or endothelial cells overnight before the measurements were made. TC: MDA-MB-231 tumor cells, Mac: BAC1.2F5 macrophages. n=120 fields at 20X magnification for TC+Mac; n= 40 fields at 20X magnification for TC+HUVEC; data plotted as mean ± SEM and normalized to the TC only control; unpaired two-tailed Student’s t test, *** p-value=0.0001, n.s. p-value=0.4226. (E) MDA-MB-231-LM2 cells (TC) were co-cultured with BAC1.2F5 cells (Mac) for 36 hours with or without a 3 μm pore insert to separate the cell populations and assessed for the number of SORE6+ cells by flow cytometry. Data are normalized to the TC only condition. Results are plotted as mean ± SEM (n=3); Tukey’s multiple comparisons test; p-values: ** = 0.0013, ns = 0.0919, *** = 0.0002 (F) Left: Schematic showing the TC+Mac co-culture experimental setup and quantification procedure for the fate mapping of non-CSCs (S6-) in time-lapse movies. Right: Quantification of SORE6 induction (conversion of S6-cell to S6+) or non-induction without or with macrophage contact in TC+Mac co-culture assay. Data plotted as mean+/- SEM, unpaired two-tailed Student’s t test, n=113 S6-cells from 28 fields of 330×330 μm^2^. See details in M+M section “Time Lapse Imaging of in vitro tumor cell-macrophage co-culture.” (G) Left: Schematic showing the TC alone or TC+Mac co-culture experimental setup and quantification procedure for the origin mapping of CSCs (S6+) in time-lapse movies. Right: Quantification of SORE6 non-induction or induction (conversion of S6-cell to S6+) in TC alone (-) or TC+Mac (+) co-culture assay respectively. Data plotted as mean+- SEM, unpaired two-tailed Student’s t test, n=4 wells/condition for a total of 128 cells traced/condition. See details in M+M section “Time Lapse Imaging of in vitro tumor cell-macrophage co-culture.” (H) Quantification of SORE6+ cells in tumor cells (MDA-MB-231 cells expressing tdTomato and with SORE6>GFP) treated with either the Jagged1 Scramble peptide or with functional Jagged1 peptide. Cells were incubated with peptides overnight before the measurements were made. n=80 fields at 20X magnification; data plotted as mean ± SEM and normalized to the TC+Jag1 scramble control; unpaired two-tailed Student’s t test, ** p-value=0.0012. (I) Quantification of SORE6+ cells in tumor cells + macrophages co-cultured and treated with DMSO or with DAPT, γ-secretase inhibitor. n=60 fields at 20X magnification; data plotted as mean ± SEM and normalized to the DAPT condition; unpaired two-tailed Student’s t test, ** p-value=0.0061. (J) Quantification of SORE6+ cells in tumor cells + macrophages co-culture assay. Tumor cells were treated with either control siRNA or Notch1 siRNA for 36 h and co-cultured with macrophages overnight. Measurements were made 48 h post siRNA transfection. n=32 fields of 330×330 μm^2^; data plotted as mean ± SEM and normalized to the Notch1si condition; unpaired two-tailed Student’s t test, * p-value=0.0232. (K) Quantification of changes in Sox9^Hi^ cells in fixed PyMT mammary tumor after PBS (control) or DAPT treatment. n=5 and 7 mice for control and DAPT conditions; two, 2-6 mm^2^ tumor tissue areas were analyzed in each mouse; data normalized to Sox9^Hi^ cells value for control treatment and plotted as mean ± SEM, unpaired Student’s t test, **** p-value<0.0001. (L) Quantification of changes in macrophage density in fixed PyMT mammary tumor after PBS (control) or DAPT treatment. n=5 and 7 mice for control and DAPT conditions; two, 2-6 mm^2^ tumor tissue areas were analyzed in each mouse; data normalized to macrophage density value for control treatment and plotted as mean ± SEM, unpaired Student’s t test, n.s. p-value=0.41.

To confirm and mechanistically dissect these *in vivo* observations we performed *in vitro* co-culture experiments. Tumor cells with the SORE6>GFP sensor and macrophages (unlabeled) were co-cultured and imaged live. Similar to our *in vivo* experiments, we observed induction of the stem phenotype (GFP positivity) in non-stem cells upon macrophage contact (Fig S6B; Movie 11). The kinetics of stemness induction *in vitro* (Fig S6C) were similar to those observed *in vivo* (Fig 4B). Only tumor cells that made direct physical contact with macrophages were induced to become SORE6+ (Fig S6B). Moreover, we found that prior co-culture with macrophages for 24 hours increased the tumorsphere-initiating activity of the tumor cells (Fig 4C), indicating that the induced stem phenotype is both durable and functional.

To determine if the induction of stemness was specific to macrophages, we co-cultured tumor cells with either macrophages or endothelial cells. In co-cultures there was a 40% increase in the SORE6+ CSCs when tumor cells were co-cultured overnight with macrophages compared to tumor cells alone (Fig 4D). This effect was specific to macrophages, as we did not see any increase in SORE6+ CSCs when tumor cells were co-cultured with endothelial HUVEC cells, the other cell type most frequently in contact with disseminating tumor cells at perivascular regions of the tumor (20) (Fig 4D). The macrophage co-culture assay was further repeated in two additional breast tumor models: MDA-MB-231-LM2 and 4T1. We found approximately 20% and 40% increase in SORE6+ CSCs in these two models resulting from direct contact with macrophages (Figs. 4E and S6D), respectively. In addition, we used human M2-like macrophages in the co-culture assay with MDA-MB-231 cells, and found approximately 25% increase in SORE6+ CSCs (Fig S6E). The increase in CSCs in macrophage/tumor cell co-cultures was not seen if macrophages and tumor cells were separated by a 3 μm pore-size filter (Fig 4E), confirming that direct cell contact, not conditioned medium, is required for induction of stemness. Taken together, these results indicate a general role for macrophage contact in inducing stemness de novo.

To rigorously address the quantitative contribution of this direct induction of stemness in non-stem cancer cells upon macrophage contact, we performed cell-fate and cell-origin mapping experiments from video microscopy movies of the co-cultures *in vitro*. Assessing the fate of randomly-selected SORE6-non-CSCs in macrophage/tumor cell co-cultures, we showed that SORE6-cells that underwent direct contact with macrophages in the time-period had a >4x increase in frequency of conversion to SORE6+ CSCs compared with SORE6-cells that did not contact macrophages, with 23% of the non-stem cancer cells that contacted a macrophage undergoing conversion to a stem cell phenotype in the time-frame analyzed (Fig 4F). In a complementary and independent experiment, we mapped the origin of SORE6+ CSCs that were present at the end of the culture period in either tumor cell/macrophage co-cultures or cultures of tumor cells alone. We similarly found that the macrophage co-cultures showed a >4x increase in the direct non-stem to stem conversion frequency (Fig 4G).

Consistent with the finding above that changes in macrophage numbers are not associated with changes in the proliferation of CSCs in PyMT tumors (Fig S6A), we found that macrophage contact *in vitro* did not change the proliferation of the CSCs indicating that macrophages do not induce the expansion of pre-existing CSCs (Fig S6F). Thus, from our *in vitro* and *in vivo* experiments, we conclude that contact with macrophages increases the cancer stem cell population through direct de novo induction of the stem phenotype in non-stem cells, without any impact on the proliferation of pre-existing stem cells.

The Notch signaling pathway was recently shown to be involved in mediating cross-talk between macrophages and the normal mammary stem cell during mammary gland development (39). Furthermore, Notch/Jagged1 signaling can promote cancer stem cell traits in homotypic cultures of breast cancer cells (40). Given studies showing Jagged1 (a Notch ligand) expression in macrophages (41, 42), and Notch1-mediated induction of Mena^INV^ expression which is required for intravasation in association with macrophages at TMEM (32), we investigated if Notch signaling plays a role during the stemness induction by macrophages that we observed in mammary tumors in vivo. Cancer cells containing the SORE6 sensor were incubated with a scrambled peptide or Jagged1 for 24 hours. Cells were fixed, imaged and analyzed for SORE6 (FLAG-expressing) cells. We found a significant increase in SORE6+ CSCs after Notch signaling activation with Jagged1 but not with scrambled peptide (Fig 4H). Conversely, inhibition of Notch signaling with DAPT, a γ-secretase inhibitor which blocks Notch signaling (43), led to a decrease in SORE6+ CSCs in cancer cell-macrophage co-cultures (Fig 4I). To address possible off-target effects of DAPT inhibition (44), we knocked down Notch1 protein in cancer cells using Notch1 siRNA (Fig S6G) and co-cultured them with macrophages. We found that macrophage contact is not able to induce stemness in Notch1 signaling defective cancer cells (Fig 4J). Furthermore, we inhibited Notch signaling *in vivo* by treating PyMT mice with DAPT (Fig S6H) and investigated primary tumors for Sox9^Hi^ cancer stem cells. We found that Notch inhibition led to a five-fold decrease in CSCs in DAPT-treated mice compared to the vehicle control-treated mice (Fig 4K), while not affecting the percentage of macrophages in the tumor (Fig 4L).

These results, together with clodronate results in Figs 3G and 3H, confirm that stemness induction is dependent on physical contact involving Notch signaling between cancer cells and macrophages.

### 5. Breast CSCs preferentially associate with and intravasate at TMEM doorways

The cancer cells capable of forming invadopodia are crucial for the function of intravasation portals called tumor microenvironment of metastasis (TMEM) through which cancer cells enter blood vessels and systemically disseminate (20, 29, 33). TMEM is a triple cell complex composed of a Mena^Hi^ tumor cell, a macrophage and a blood vessel endothelial cell, all in direct physical contact on the blood vessel surface. TMEM doorways are the only known portals for tumor cell intravasation in mammary tumors (20, 45). The density of TMEM doorways is a clinically validated prognostic marker of metastatic recurrence in breast cancer patients (46–48). Since CSC form invadopodia which are required for TMEM function and transendothelial migration; and macrophage contact induces stemness in cancer cells, and the areas around TMEM are a perivascular macrophage-rich compartments (20, 49), we investigated if CSCs are enriched at and around TMEM doorways using intravital imaging. We quantified the location of SORE6+ stem cells with respect to TMEM doorways, or blood vessels away from TMEM. We performed 4-color time-lapse intravital imaging of mammary tumors expressing SORE6>GFP in a Rag2 KO/MacBlue host, where macrophages are labeled in CFP (20). We found CSC enrichment in perivascular regions near TMEM (Fig 5A).

**Figure 5:**
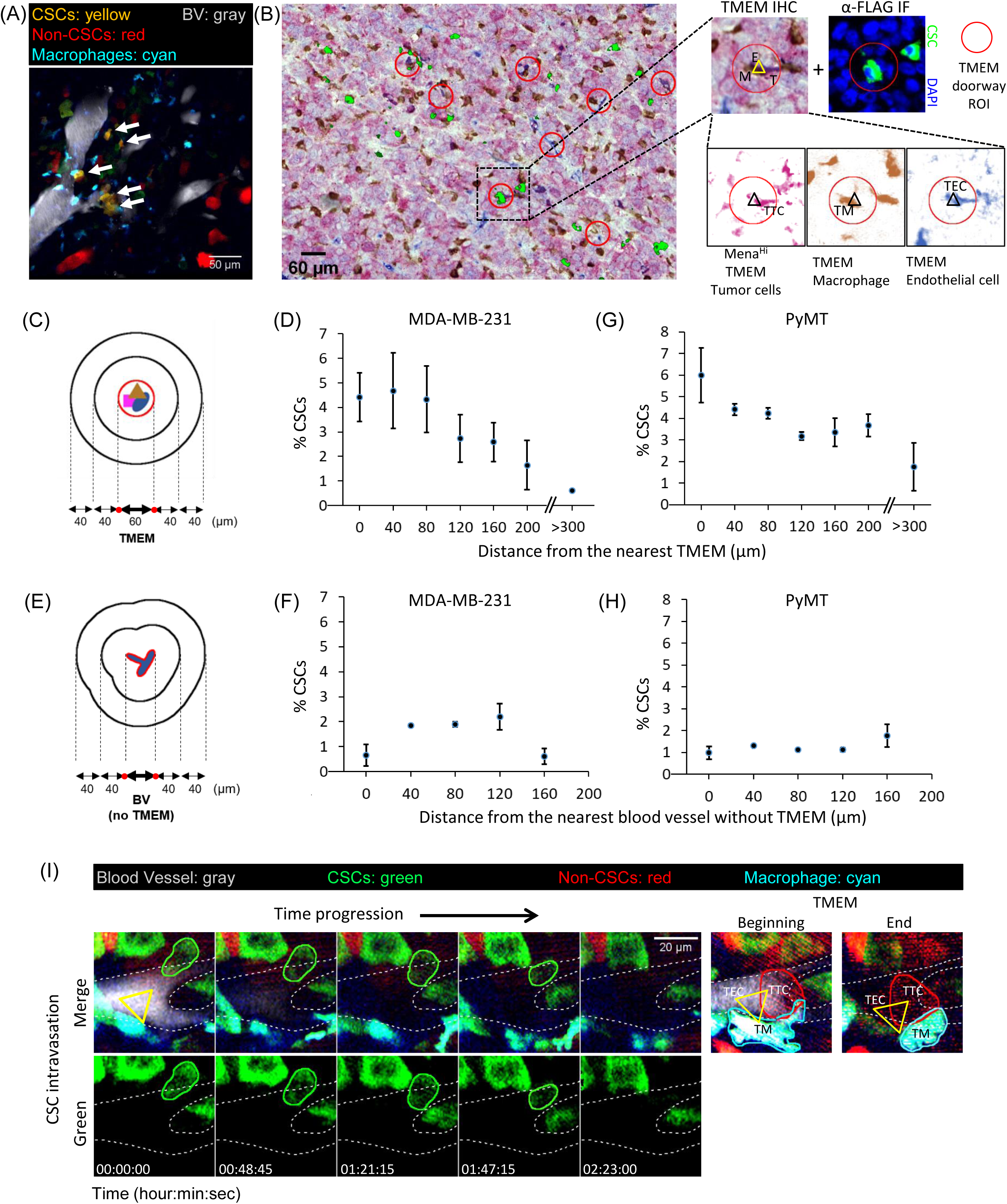
Breast carcinoma stem cells preferentially associate with and intravasate at TMEM doorways. (A) Still image from time-lapse intravital Movie 12 of SORE6>GFP xenograft MDA-MB-231 tumor in Rag2 KO mice, showing CSCs (yellow: tdTomato volume marker + SORE6-GFP) enriched in perivascular regions and in contact with perivascular macrophages (cyan). Blood vessels were labeled by perfusing far-red Qdots (gray) via a tail-vein catheter. (B) A composite image of TMEM staining (pink: Mena^Hi^ tumor cells, brown: macrophages, blue: endothelial cells) and FLAG+ stem cells (shown in green). TMEM doorway ROI are highlighted with red circles. Zoomed images show an example of FLAG+ stem cell (α-FLAG IF panel) being associated with the TMEM doorway (TMEM IHC panel) when panels are aligned. Triple-stained TMEM IHC image is further deconvolved into individual channels and shows that TMEM (marked with triangle) is composed of a TMEM tumor cell (TTC) (Mena^Hi^), TMEM macrophage (TM) (Iba-1), and an TMEM endothelial cell (TEC)(endomucin). (C) The schematic shows how the region containing TMEM within the section is assigned a boundary surrounded by concentric contours radiating out from the TMEM in 40 μm intervals. The TMEM boundary (marked by red dots) and concentric contours are projected onto a linear axis at the bottom of the schematic. For scoring purposes each TMEM doorway is visualized as 60 μm diameter circle and the CSCs were enumerated relative to how close they are to the TMEM circle using a custom-written ImageJ macro. Cells which are inside or touching the TMEM boundary are counted in data point 0 μm in Figs 5D, G, with subsequent cell counts measured within each 40 μm interval area contour moving away from the TMEM. (D) Percentage CSC (anti-Flag stained for SORE6+ cells) distribution from the nearest TMEM in fixed MDA-MB-231 mammary tumor tissue. Note that, there are 7x enrichment of CSCs associated with TMEM than non-CSCs compared to the average CSCs found throughout the tumor at sites farther from TMEM (>300 μm). n=4 fields of 2.2×2.2 mm^2^ from 4 mice; data plotted as mean ± SEM. (E) Schematic shows how the region containing a blood vessel without TMEM within the section is assigned a boundary surrounded by concentric contours radiating out from the blood vessel in 40 μm intervals. The blood vessel boundaries (marked by red dots) and concentric contours are projected onto a linear axis at the bottom of the schematic. For scoring purposes each blood vessel is visualized as vessel and the CSCs were enumerated relative to how close they are to the vessel using a custom-written ImageJ macro. Cells which are touching the vessel are counted in data point 0 μm in Figs 5F, H, with subsequent cell counts measured within each 40 μm interval area contour moving away from the vessel. (F) Percentage CSC (anti-Flag stained for SORE6+ cells) distribution from the nearest blood vessel without TMEM in fixed MDA-MB-231 mammary tumor tissue. Note that, there is no enrichment of CSCs relative to the nearest blood vessel without TMEM. n=4 fields of 2.2×2.2 mm^2^ from 4 mice; data plotted as mean ± SEM. (G) Percentage CSC (anti-Sox9 stained) distribution from the nearest TMEM in fixed PyMT mammary tumor tissue. Note that, there are 3.4x enrichment of CSCs associated with TMEM than non-CSCs compared to the average CSCs found throughout the tumor at sites farther from TMEM (>300 μm). n=8 fields of 2-3 mm^2^ from 4 mice; data plotted as mean ± SEM. (H) Percentage CSC (anti-Sox9 stained) distribution from the nearest blood vessel without TMEM in fixed PyMT mammary tumor tissue. Note that, there is no enrichment of CSCs as a distance from the nearest blood vessel without TMEM. n=11 fields of 2-3 mm^2^ from 5 mice; data plotted as mean ± SEM. (I) Time-lapse panels from intravital movie 13 showing intravasation of SORE6+ CSC (intravasating CSC outlined in green) into blood vessels in the primary tumor. Top panels show merged channels and bottom panels show green channel for a clearer view of intravasating CSC. Blood vessels (outlined by white dotted lines) were visualized by tail-vein injection of far-red Qdots (gray). Right panels showing TMEM (which are taken from the same X-Y field but are in adjacent Z-planes since the TMEM doorway is above the point of intravasation of tumor cells entering the blood vessel as described previously (20)), show that intravasation occurs at TMEM (TTC, TM and TEC are the TMEM tumor cell, macrophage and blood vessel endothelial cell, respectively, forming the three cell contacts (yellow triangle) identifying TMEM). Note the TMEM structure is stable and seen in the beginning of the time course and the end. Yellow triangles in left time progression panels depict the TMEM locations copied over from the right TMEM panels. Note that the tumor cell in the TMEM panels to the right, which is part of the TMEM triangle, is different from the intravasating CSC shown in left time progression panels, and does not intravasate as consistent with previous work (20).

To further investigate stem cell associations with TMEM doorways, we performed triple IHC for TMEM markers (Mena, Iba-1 and CD31) in fixed tumors. TMEM are identified as three cells in direct and stable physical contact by triple staining of TC (Mena^Hi^), macrophage (IBA-1) and blood vessel (CD31). A sequential tissue section was stained with FLAG antibody to identify SORE6+ CSCs. TMEM IHC and FLAG IF images were then aligned and superimposed. As suggested by imaging of live tissue (Fig 5A, Movie 12), we found cancer stem cells in association with TMEM doorways (Fig 5B), with approximately 38% of TMEM doorways being assembled with a cancer stem cell (Fig S6I).

TMEM IHC and FLAG IF images were further analyzed to determine the percentage of CSCs relative to TMEM doorways and blood vessels without TMEM doorways. Approximately 50% of blood vessels contain at least 1 TMEM and 50% are free of TMEM. An enrichment of CSCs was observed in the tumor areas close to TMEM doorways (Figs. 5C and 5D) resulting in about 7 times higher percentage of CSCs compared to > 300 μm away from TMEM doorways (Fig 5D). We also determined the percentage of CSCs relative to blood vessels without TMEM doorways and did not see any enrichment (Figs. 5E and 5F), indicating that CSCs specifically cluster around TMEM doorways. We repeated our analysis in another mammary tumor type (PyMT), with Sox9 marking the CSCs. Again, we found CSC enrichment (3.4x) around TMEM doorways compared to the average seen throughout the tumor at sites not near TMEM (Fig 5G). As seen in the MDA-MB-231 tumors, in PyMT tumors we also did not observe any CSC enrichment around blood vessels not containing any TMEM (Fig 5H). These results suggest that the induction of CSCs occurs in association with the macrophages proximal to TMEM doorways (20, 49), leading to increased CSCs in association with each TMEM doorway. This raises the key question of whether TMEM doorways are not only niches for CSCs but also intravasation portals for seeding of metastatic CSCs.

To determine if CSCs, that are associated with TMEM doorways, intravasate at TMEM, we used intravital imaging as described previously to directly observe tumor cell intravasation in primary tumors (20). We observed intravasation of CSC, identified by expression of the SORE6-GFP biosensor (Fig 5I, Movie 13). High resolution imaging of mice with CFP-macrophages showed that the intravasation occurred at TMEM doorways (Fig 5I right panels show labeled TMEM).

### 6. CSCs are highly enriched in circulation and at their first arrival in the lung, where they seed new micro-metastases

Consistent with the intravasation of CSCs at TMEM doorways, we found CSCs in the circulation (Fig 6A). However, there was a large enrichment of CSCs in the circulating tumor cell (CTC) population, where CSCs were over 60% of CTCs, while CSCs at TMEM doorways waiting to intravasate comprised only ∼4-5% of the tumor cell population at that location (Figs. 6B and 5D).

**Figure 6:**
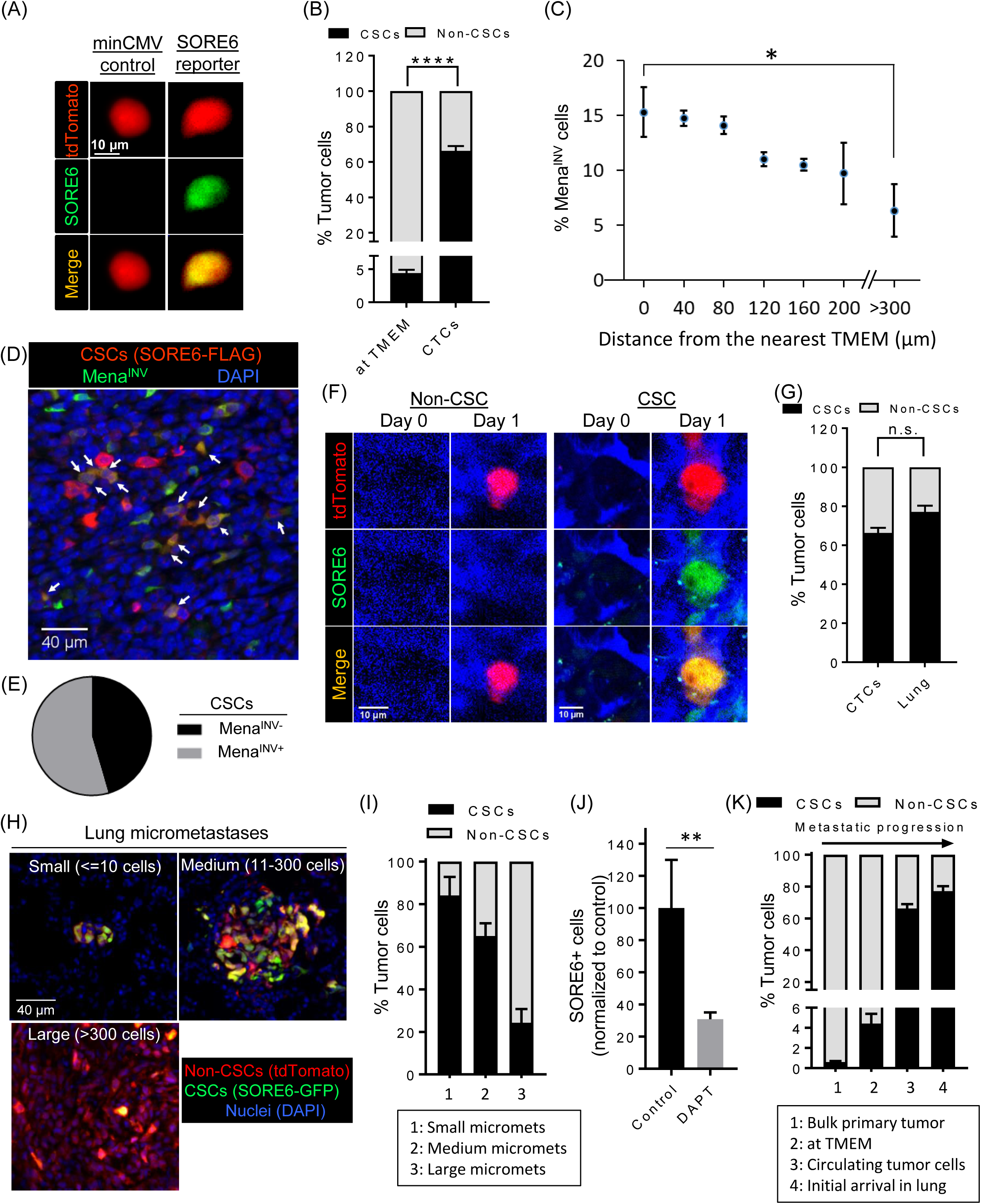
CSCs are highly enriched in circulation and at their first arrival in the lung, where they seed new micro-metastases. (A) CSCs are found in CTCs collected from the mice imaged in figure 5I. Images of stem cell (top panels – tdTomato red, middle panels – SORE6-GFP, green) in the circulation. (B) Quantification of % CSC and non-CSCs in the circulation (n= 3 mice). For comparison, % CSC (SORE6-GFP) and non-CSC values at TMEM doorways were plotted, and shows significant enrichment of CSCs in the circulation compared to their numbers at the TMEM doorway. Data plotted as mean ± SEM, unpaired two-tailed Student’s t test, **** p-value<0.0001 (C) Percentage Mena^INV^ cell distribution from the nearest TMEM in fixed MDA-MB-231 mammary tumor tissue. Note that, there is approximately 2.4x enrichment of Mena^INV^ cells at TMEM doorways compared to Mena^INV^ cells found throughout the tumor at sites farther from TMEM (>300 μm). n=6 fields of 1200×650 μm^2^ from 3 mice having MDA-MB-231 primary tumors stained with antibodies for Mena^INV^ and TMEM doorways; data plotted as mean ± SEM, Mann Whitney test, * p-value=0.038 (D) *In situ* immunofluorescence staining of MDA-MB-231 FFPE primary tumor tissue with anti-FLAG antibody (red, to identify SORE6-FLAG CSC), and Mena^INV^ antibody (green). Double positive (stem + Mena^INV^) cells are seen in yellow (white arrows). (E) Quantification of fraction of double positive Mena^INV^+ stem cells compared to CSCs that are Mena^INV^ negative plotted as a pie chart. Full circle represents all stem cells and shows that ∼55% of stem cells are positive for Mena^INV^ expression. n= 7 fields of 1200×650 μm^2^ from MDA-MB-231 FFPE primary tumor tissue. (F) Representative images of single non-CSC (tdTomato red) and CSC (SORE6+ green, and yellow in merged channel) cells arriving in the lung spontaneously from the primary tumor, imaged with intravital microscopy at the lung site. Note the absence of cancer cells on day 0 and presence of cancer cells on day 1 in the same field of view. The 10 kDa Cascade blue-dextran was used to visualize vascularized areas of the lung (blue). (G) Quantification of newly arriving % CSC and non-CSC cells in the lung shows significant enrichment of CSCs at the secondary site (lung) similar to their numbers in the CTCs (Fig. 6B); n=18 stem cells, 6 non-stem cells in 5 mice, data plotted as mean ± SEM, unpaired two-tailed Student’s t test, n.s. p-value=0.11 (H) Representative images of fixed frozen lung tissue show metastases at different stages of progression. Small micro-metastases (<=10 cells), medium micrometastases (11-300 cells) and large micrometastases (>300 cells) show progressive loss of CSCs as size increases. (I) Quantification of % tumor cells (CSCs and non-CSCs) in lung metastases at different stages of progression as shown in 6H. n=70 metastases analyzed in 3 mice; data plotted as mean ± SEM. (J) Quantification of SORE6+ cells in the lungs of control or DAPT treated SCID mice bearing SORE6>GFP MDA-MB-231 tumors. FFPE lung tissues were stained with FLAG antibody to identify SORE6+ CSCs. n=5 and 6 mice for control and DAPT conditions respectively; two, 1.8×1.8 mm^2^ lung areas were analyzed in each mouse; data normalized to % SORE6+ values for control treatment and plotted as mean ± SEM, Mann-Whitney test, ** p-value=0.002. (K) The % CSC and non-CSC values for the bulk primary tumor (Fig 1D, FFPE), at TMEM (Fig 5D), in circulation (Fig 6B) and initial arrival in the lung (Fig 6G) were plotted, and show progressive enrichment in cancer stem cells up to the point of initial metastatic seeding in the lung.

One possible explanation for this is that CSCs form invadopodia (Fig. 2) and are, therefore, much more efficient at intravasation than non-CSCs from the same primary tumor. Previous work has shown that the assembly of invadopodia and invadopodium-dependent transendothelial migration during intravasation at TMEM depends on Mena^INV^ expression (31) which is induced by macrophage-driven Notch1 signaling in breast tumors *in vivo* (32). To investigate the inter-relationship between Mena^INV^+ cells and CSCs, first we investigated the preferential enrichment of Mena^INV^ in tumor cells associated with TMEM doorways as compared to elsewhere in the primary tumor. As shown in Fig. 6C, there is a 2.4-fold increase in Mena^INV^ expressing tumor cells in the proximity of TMEM doorways compared to the areas away from TMEM indicating enhanced intravasation potential of cancer cells around TMEM. Since CSCs are also enriched at TMEM doorways we wanted to determine if CSCs at the primary tumor also express Mena^INV^. We co-stained primary tumor tissue with FLAG (to identify SORE6+ stem cells) and Mena^INV^ antibodies, and found Mena^INV^ expression in more than 55% of CSCs (Fig. 6D, E). The data support enhanced intravasation activity of these double positive (stem+Mena^INV^) tumor cells at the TMEM doorway, consistent with the enrichment of stem cells seen in circulation (Fig. 6B).

To address the relative efficiency of circulating CSCs and non-CSCs in metastasizing to distant organs such as lung, we performed high-resolution intravital imaging of live lungs using a permanent lung imaging window to visualize the arrival of single circulating tumor cells. Using the permanent lung window and its associated micro-cartography technique to return to the same imaging field over time (33), multiple fields were imaged on two consecutive days. We saw arrival of both CSCs and non-CSCs at the lung (Fig 6F), with the majority (>70%) of cancer cells being CSCs (Fig 6G), similar to their enrichment numbers seen in circulation (Fig. 6B). Taken together, these results indicate that the tumor cell population becomes progressively more enriched for cancer stem cells after passage through TMEM and during the dissemination and seeding of metastasis (Figs. 6B, 6G).

Next, we investigated if the newly arriving tumor cells establish metastases in the lung, and we analyzed relative proportions of CSCs and non-CSCs in the metastases as they grew in size, using fixed-frozen lung sections to identify SORE6+ CSCs relative to all tumor cells. Lung micrometastases composed of a few tumor cells to hundreds of tumor cells were observed (Fig 6H). SORE6+ CSCs were quantified in small (<= 10 tumor cells), medium (11-300 tumor cells) and large (> 300 tumor cells) lung micrometastases. Small lung micrometastases were greatly enriched in stem cells (∼ 84% of total tumor cells) (Fig 6I), suggesting that early lesions are primarily seeded by CSCs. This conclusion is consistent with our observations that the majority (>70%) of tumor cells arriving in the lung are CSCs (Fig 6G), and our previously published demonstration that SORE6+ cells (CSCs) initiate metastasis with ∼8-fold higher efficiency than SORE6-cells (non-CSCs) (13). As the lung metastases size increased, the proportion of stem cells in the metastases decreased, with only 24% SORE6+ CSCs in metastases with more than 300 cells (Fig 6H, I). An independent study, using flow cytometry rather than imaging, showed a similar progressive decrease in the relative CSC representation with time in tumor cells recovered from metastasis-bearing lungs (50). Both datasets are consistent with the expectations of the cancer stem cell hypothesis that CSCs will generate more differentiated non-CSC progeny (51). However, we cannot exclude the possibility that the non-CSCs have a proliferative advantage at the metastatic site, or that a microenvironmental stimulus supporting the CSC phenotype is lost in larger lesions.

Next, we investigated if macrophage-induced Notch signaling plays any role in the metastatic spread of CSCs. We treated SORE6>GFP MDA-MB-231 bearing SCID mice with vehicle control or DAPT, and stained lungs extracted from these mice with FLAG antibody to identify SORE6+ CSCs. We found a three-fold decrease in CSCs in the lung of DAPT-treated mice compared to the control-treated mice (Fig 6J), suggesting a casual role of macrophage-induced Notch signaling in CSC metastatic potential.

Overall, our data show that during the course of dissemination of tumor cells from the primary site, CSCs become progressively enriched in the tumor cell population as they approach the TMEM doorway, intravasate, circulate and arrive at the lung (Fig 6K). Association with and passage through the TMEM doorway potentially generates the greatest enrichment in CSCs (∼ 60-fold), which could be explained by the product of a combination of CSC enrichment at TMEM (7-fold, Fig. 5D), Mena^INV^ enrichment in tumor cells at TMEM (2.4-fold, Fig 6C), and a previously documented 3.5-fold increase in intravasation potential of tumor cells expressing Mena^INV^ (31, 32). These previous studies in addition to our current study could account for the enrichment of CSCs in the CTC population. On arrival in the lung, CSCs represent more than 75% of the disseminated tumor cell population, greatly enriched compared with their representation in the bulk primary tumor of ∼ 1%. Since the CSCs are similarly enriched in the smallest micrometastatic lesions in the lung, the data strongly suggest that CSCs are the main drivers of the early steps of metastatic dissemination, with the generation and expansion of non-CSC progeny driving the later steps of metastatic colonization and growth.

### 7. The density of TMEM doorways correlates with the proportion of CSCs in breast cancer excisions from patients

Since we found that, in the pre-clinical models of breast cancer, areas around TMEM are enriched for CSCs, and that macrophages can induce a stem phenotype, we hypothesized that human breast cancers with high TMEM density would also have a high proportion of CSCs. If high TMEM density is accompanied by a high density of CSCs, this would provide an additional mechanistic insight into the prognostic power of TMEM score for metastatic outcome in patients (46–48) beyond the function of TMEM as a cancer cell intravasation doorway. To investigate the relationship between TMEM density and the proportion of CSCs in breast cancers in patients we used 49 breast cancer excisions from patients who had various breast cancer subtypes (Figure 7A), and a special tissue collection approach described in Figure 7B. Briefly, we collected cancer cells by fine needle aspiration (FNA) from breast cancer excisions as described under Materials and Methods. This approach has several benefits. It allowed us to sample large portions of tumors since the aspiration was performed from at least 3 distinct tumor areas with 5 to 10 needle passes for each tumor area. Furthermore, the FNA sample contained mostly single cells which facilitated the flow cytometry analysis of the expression of CD44+/24-on the cell surface, a commonly used marker combination associated with stemness which is difficult to analyse in fixed tissues (3). Lastly, this approach was not associated with appreciable tissue destruction as assessed by histopathology which allowed us to use the same tumors upon formalin fixation for the analysis of TMEM score. Thus, upon completion of FNA, the very same breast cancer excisions were formalin-fixed and paraffin-embedded and the representative tissue sections were used for TMEM density analysis. We then correlated the density of TMEM doorways with the proportion of cancer cells expressing stem cell markers standardly used for detection of cancer stem cells in breast cancers from patients. We wanted to take into consideration the plasticity that breast cancer stem cells (BCSCs) exhibit as they transit between proliferative, epithelial-like (E) and a quiescent, invasive, mesenchymal-like state. To accomplish that, we used CD44+/ CD24-which marks BCSC in the mesenchymal-like state and aldehyde dehydrogenase 1 (ALDH1) which marks BCSC in the epithelial-like state (52). As an additional marker of stem-phenotype we used CD133, because it is the most frequently used maker of stemness in solid cancers from patients (53, 54) (Fig 7D). We used a variety of methods for detecting stem cell markers to evaluate the results in a technique independent way (Fig. 7 C-E). With all methods we observed an extraordinarily strong positive correlation between TMEM density and the proportion of cancer cells expressing stem cell markers (CD44+/24-, CD133 and ALDH1) (Fig 7C-D).

**Figure 7:**
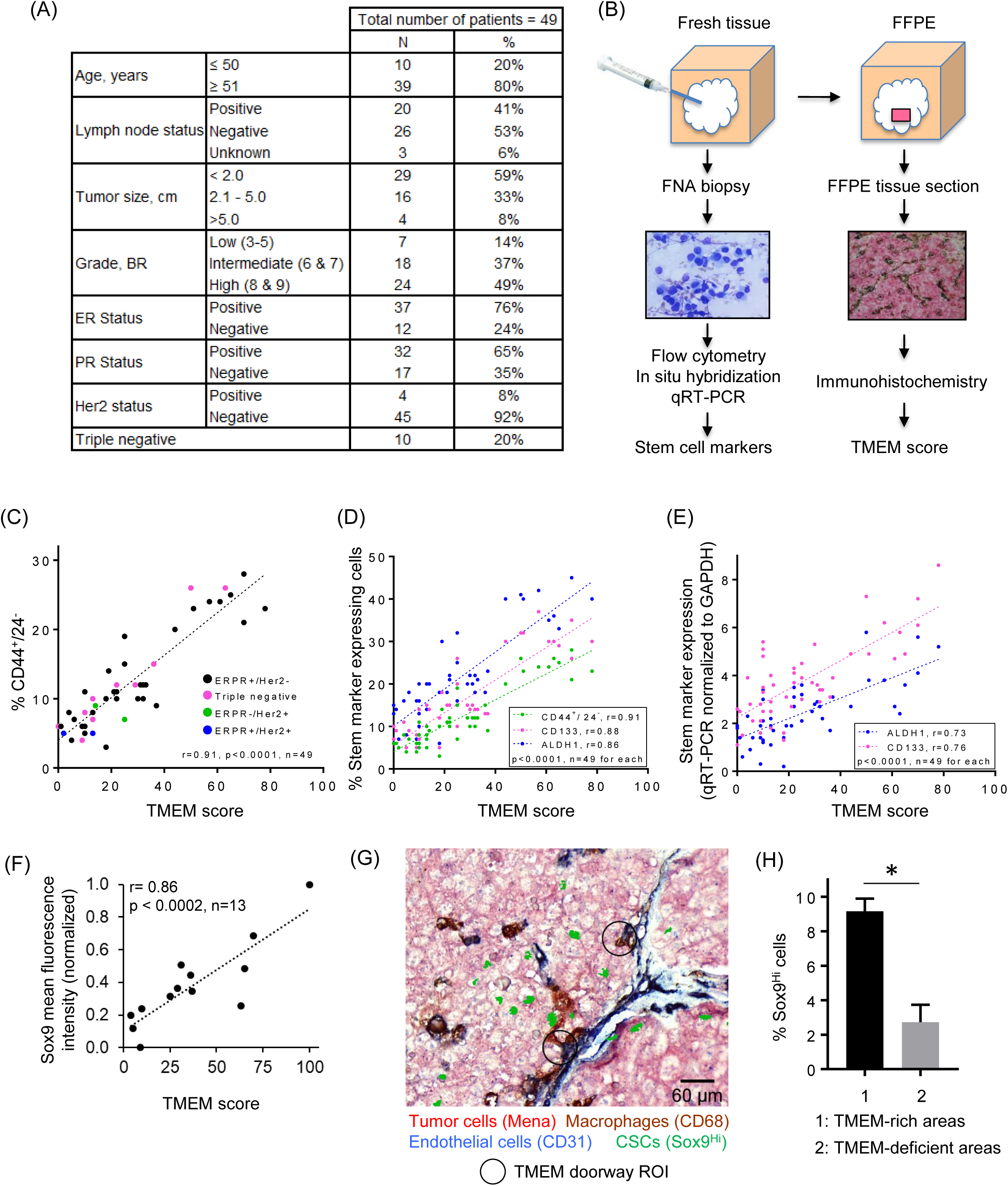
Correlation of TMEM density with the proportion of tumor cells expressing stem cell markers in breast cancer tissues from patients. (A) Clinical and pathological characteristic of the 49 breast cancer cases included in the analysis. The tissue was collected from 49 patients under the Albert Einstein College of Medicine/ Montefiore Medical Center Institutional Review Board protocol approval. (B) Tissue collection procedure for the correlation of TMEM density with the proportion of cancer cells expressing stem cell markers. Fresh tumor was first sampled by fine needle aspiration (FNA) biopsy. Following the FNA procedure the entire lesion was formalin-fixed and paraffin-embedded (FFPE). The cells obtained by FNA were analyzed by flow cytometry for the expression of CD44+/CD24-. They were also analyzed for the expression of ALDH1 and CD133 by in situ hybridization and qRT-PCR. A representative tissue section of the corresponding FFPE tumor was chosen, immunostained for TMEM and the TMEM density was scored. Thus, the TMEM analysis and the analyses for the expression of stem cell markers were blinded and done on the very same tumor. The TMEM density was then correlated with the expression of stem cell markers. (C) TMEM correlation with stem cell marker CD44+/24- in 49 invasive ductal carcinomas. Breast cancer cells collected by fine needle aspiration (FNA) from breast cancer excisions and analyzed by flow cytometry for the percentage of cells expressing high level of CD44 and low level of CD24. Upon formalin fixation and paraffin embedding, breast cancer tissue sections from corresponding tumor samples were stained for TMEM and TMEM score was correlated with the percentage of CD44+/24- cells by Pearson’s correlation. (D) Correlation of TMEM with the percentage of breast cancer cells expressing stem cell markers CD44+/24-, ALDH1 and CD133 in 49 invasive ductal carcinomas from patients. We collected breast cancer cells by fine needle aspiration (FNA) from breast cancer excisions and analyzed the percentage of cells expressing high level of CD44 and low level of CD24 by fluorescence activated cell sorting (FACS), as well as CD133 and ALDH1 by in situ hybridization. Upon formalin fixation and paraffin embedding, breast cancer tissue sections from corresponding tumor samples were stained for TMEM and TMEM score was correlated with the percentage of cancer cells expressing stem cell markers using Pearson’s correlation. (E) Correlation of TMEM with the percentage of breast cancer cells expressing stem cell markers ALDH1 and CD133 in 48 invasive ductal carcinomas from patients. We collected breast cancer cells by fine needle aspiration (FNA) from breast cancer excisions and analyze the percentage of cells expressing CD133 and ALDH1 by qRT-PCR. Upon formalin fixation and paraffin embedding, breast cancer tissue sections from corresponding tumor samples were stained for TMEM and TMEM score was correlated with the percentage of cancer cells expressing stem cell markers using Pearson’s correlation. (F) Sox9 mean fluorescence intensity in FFPE of the above human tissues (n=13) were correlated with the TMEM score for each human tissue. n=13 human tissues, Pearson’s correlation r=0.86, p<0.0002, data normalized to the min and max Sox9 mean fluorescence intensity values. (G) A composite image of TMEM staining (pink: Mena+ tumor cells, brown: CD68+ macrophages, blue: CD31+ endothelial cells) and Sox9^Hi^ stem cells (shown in green), showing enrichment of Sox9^Hi^ cells close to TMEM areas. Two TMEM doorway ROIs are highlighted with black circles. (H) Quantification of % Sox9^Hi^ cells in TMEM-rich and TMEM-deficient areas. n=3 human tissues, data plotted as mean ± SEM, paired two-tailed Student’s t test, p<0.034.

In addition to the percentage of cells expressing ALDH1 and CD133 as determined by in situ hybridization, we also wanted to evaluate the total amount of ALDH1 and CD133. Therefore, we determined the amount of mRNA transcript for ALDH1 and CD133 by qRT-PCR. We then correlated the amount of ALDH1 and CD133 mRNA measured by qRT-PCR (normalized to GAPDH) with TMEM density and again obtained significant correlation (Fig 7E).

Since SORE6 biosensor cannot be expressed in human samples, to identify CSCs in situ, we used Sox9 as a surrogate stemness marker in fixed tissue. We stained FFPE human tissues with Sox9 antibody and correlated Sox9 expression (mean fluorescence intensity) in each tissue with its corresponding TMEM density and found significant correlation between Sox9 expression and TMEM density in human tissues (Fig 7F). To check spatial distribution of stem cells with respect to TMEM doorways in human tumors, we stained sequential tissue sections for TMEM doorways using a humanized version of triple-IHC (panMena, CD68 and CD31 antibodies) and for stem cells using Sox9 antibody. After aligning TMEM and Sox9 sections, we found significant enrichment of Sox9^Hi^ cells close to TMEM-rich areas (Fig 7G). Quantification showed 3-fold enrichment of Sox9^Hi^ cells in TMEM-rich areas compared to that in TMEM-deficient areas in human tissues (Fig 7H).

Overall, these results indicate that TMEM doorways in patients may not only be sites for cancer cell intravasation but also microenvironments enriched for cancer cells with stem properties, in agreement with our animal studies.

## DISCUSSION

A major limitation in studying CSC biology has been a heavy reliance on assays that disrupt tumor microenvironments, thus preventing characterization of CSCs *in situ* in their specialized niches. Here, for the first time, we have used a validated CSC fluorescent reporter system in combination with intravital high resolution multiphoton microscopy to gain important new insights into the induction, dynamic behaviors and fates of CSCs *in vivo* in breast cancer primary tumors, and during the process of seeding of lung metastases. Additionally, we identified specific CSC-enriched niches in the tumor microenvironment and verified that the same niches exist in human breast cancers obtained from patients. These insights may have impact on patient care.

As expected (4), we found that CSCs comprise a minority (∼1%) of cancer cells in the primary tumor. *In vivo* time-lapse imaging revealed that these CSCs demonstrate a slow migratory, invasive invadopodium-rich phenotype, which is the hallmark of disseminating tumor cells (19). In contrast, the non-CSCs had a fast-migrating phenotype. Previous work has shown that the slow migratory invadopodium-rich, invasive phenotype is also present in the transendothelial migration competent cancer cells expressing high levels of Mena^INV^ (19, 31, 32). Given the disseminating phenotype of CSCs and Mena^INV^ expressing cells we evaluated their distributions within the tumor microenvironment in relationship to TMEM doorways which mediate tumor cell intravasation and dissemination, (7, 26, 55) and are prognostic for distant metastatic recurrence (46–48, 55). Interestingly, both CSCs and Mena^INV^ expressing cells are preferentially located close to the TMEM doorways. Furthermore, association with and passage through the TMEM doorways contributed to the 60-fold enrichment of CSCs among CTCs, and 70-fold upon arrival to the lungs, compared with their relative representation in the bulk primary tumor. Thus, our data suggest that TMEM doorways preferentially promote the intravasation of invasive CSCs from the primary tumor. Alternatively, they may reflect survival advantage in anchorage-independent conditions that CSC have (56) over non-CSC in circulation. Our data further indicated that these CSCs initiate metastasis at the distant site, thereby intimately linking stemness and dissemination. This concept is consistent with our observations that early metastatic lesions in the lung have a very high CSC representation (∼80%), a finding that has also been made by others using orthogonal techniques (57).

Recently it was shown by live imaging that small clusters of tumor cells can be seen at intravasation sites, and in the circulation, and that they result from the aggregation of single CSCs, instead of from collective migration and vascular invasion of non-stem tumor cells (58). Further, the aggregation seemed to be mediated by intercellular homophilic interactions of CD44, a classic CSC marker which is strongly associated with TMEM density in breast cancer patients. These findings are consistent with the enhanced metastatic seeding activity of the CSCs reported here and may explain the reports of enhanced metastatic seeding activity of tumor cell clusters (59) if clusters are primarily composed of CSCs (60).

Since self-renewal, differentiation potential and phenotypic plasticity of both normal and cancer stem cells are regulated by input from the local microenvironment (8) we used the stem cell reporter to directly observe the effect of the tumor microenvironment on cancer cell plasticity *in vivo* in real time. We found that CSCs compared to non-CSCs are more frequently present close to or in direct contact (60-70%) with intra-tumoral macrophages. Importantly, we observed a novel contact-dependent de novo induction of the CSC phenotype in non-CSC tumor cells when they directly touched macrophages, both *in vitro* and *in vivo*. The data suggest a model in which macrophages constitute a cell contact-dependent inductive signal that can promote phenotypic plasticity and re-acquisition of stem properties in more differentiated tumor cells.

Interaction between macrophages and post-EMT stem-like cells, characterized by high expression of CD90, has previously been studied in the HMLER model of human breast cancer, and in human breast cancer specimens (28). In that study it was proposed that tumor-associated macrophages constitute a supportive niche by enabling pre-existing CSCs to maintain their residence in the stem cell state, through Ephrin-dependent expression of the cytokines IL6 and IL8 (28). In addition, a juxtacrine signaling interaction between macrophages and tumor cells has implicated the LSECtin – BTN3A3 axis in CSC promotion leading to enhanced tumor growth (61).

In a significant advance, our data show that intra tumoral macrophages may not merely support survival and expansion of pre-existing CSCs leading to increased tumor growth, but can also actively induce stemness in non-stem cancer cells, resulting in a population of CSCs that is linked to systemic dissemination. This induction of stemness operates via a molecular pathway that is distinct from the Ephrin-dependent maintenance of stemness, and the LSECtin – BTN3A3 axis supporting tumor growth, and involves macrophage-tumor cell contact-dependent Notch signaling. Notch is one of the core signaling pathways that has been implicated in regulation of both normal and cancer stem cells in many organ systems (62), and in tumor cell intravasation activity (32). High expression of Notch1 and Jag1 is associated with poor overall survival in breast cancer, suggesting the importance of this signaling axis in human breast cancer (63). Our results connect the de novo induction of cancer cell stemness by contact with macrophages to the dramatic enrichment of CSCs that we see in association with and passage of tumor cells through the TMEM doorway. Since CSCs accumulate around TMEM doorways, a macrophage enriched environment that can contribute to CSC induction (20, 64), dissemination of these newly-induced CSCs is likely leading to more efficient dissemination of CSCs than that for CSCs positioned away from TMEM doorways.

The observation that macrophage-driven induction of stemness in tumor cells is associated with intravasation and dissemination through TMEM doorways is arguably the worst possible scenario from a clinical perspective, likely leading to poor patient survival. This linkage is further supported by earlier findings that macrophages induce invadopodium assembly and maturation in breast tumor cells *in vitro* and *in vivo* via a Notch signaling pathway by inducing Mena^INV^ expression (26, 32, 65). These results indicate that Notch signaling is used in common to induce both stemness and the Mena^INV^-dependent invasive invadopodium-rich tumor cell phenotype. Mena^INV^-expressing tumor cells have greatly enhanced chemotaxis toward both macrophages and blood vessels (7, 64, 66), and Mena^INV^-expression induces invadopodium assembly (65) generating an enhanced transendothelial migration phenotype that is essential for TMEM-mediated dissemination (31). If indeed both the CSC state and Mena^INV^ expression are induced at TMEM this would explain the dramatic enrichment of CSC levels among CTCs. This novel finding is consistent with a previous literature where the presence of CSCs was detected in CTCs in mouse models and breast cancer patients (60, 67).

Previous studies showed that depletion of macrophages, knock out of the VEGF gene in macrophages (20, 49), or inhibition of TMEM by genetic and pharmacologic strategies (29, 45), all lead to depletion of CTCs and inhibition of metastasis, indicating that cancer cell intravasation occurs only at TMEM (20, 29). This is in accordance with the data from breast cancer patients which indicate that TMEM density, independently of blood vessel density, is associated with tumor cell dissemination and metastasis (46–48). The enrichment of CSCs at TMEM doorways in human tumors demonstrated here supports prior work and the conclusion that TMEM doorways may be involved in the dissemination of CSCs in humans.

Given the high percentage of CSCs among CTCs, it is not surprising that the presence of CTCs in breast cancer patients correlates with worse disease-free survival (DFS) and overall survival (OS). For example, a multi-institutional analysis of individual CTC data from more than 3000 patients with non-metastatic breast cancer showed that the presence of CTCs is an independent predictor of poor DFS, distant DFS, breast cancer-specific survival and OS (68). Likewise, the detection of CTCs before the onset of pre-operative chemotherapy is an independent prognostic indicator of worse DFS and OS (69). In fact, meta-analysis of data from individual patients treated with pre-operative chemotherapy collected from 21 studies showed that the hazard ratio of death proportionally increases with the number of CTCs detected before the onset of chemotherapy (70). In addition to the sheer presence of CTCs, the gene expression pattern of CTCs also affects metastatic formation. In accordance to our data showing that SORE6+ cells (CSCs) are the source of early metastatic foci, the expression of gene combinations involving stem programs such as Notch and CD44 confers higher metastatic potential (71, 72). Likewise, the expression of CSC markers in CTC in patients with several epithelial cancer types correlates with occurrence of metastasis and decreased patient survival (73, 74). Interestingly, in patients with metastatic breast cancer the occurrence of CTC expressing not only stem markers, but also markers consistent with partial epithelial to mesenchymal transition (EMT) was shown to be an independent factor for prediction of increased relapse (75) supporting the notion that both stem and EMT programs are involved in metastasis in patients. Our data showing that SORE6+, compared to SORE6-cells express significantly more EMT transcription factor Snail 1 are in accordance with these clinical findings which link patient outcome with activation of both stem and EMT programs in CTCs.

We found a striking correlation between the density of TMEM doorways and the proportion of cancer cells expressing CSC markers in breast cancer samples from patients. These data indicate that TMEM doorways are not only portals for cancer cell dissemination to distant sites but may also represent microenvironmental niches for the induction of the stem cell program in human breast cancer cells. These data further explain the prognostic power of TMEM for metastatic disease (46–48), given the association of stem and EMT program with metastatic dissemination (73, 74, 76). Indeed, TMEM density correlates with the proportion of CSCs expressing markers associated with both proliferative, epithelial-like (ALDH1) and quiescent mesenchymal-like (CD44+/24-) states (52). This transition of CSCs between the epithelial- and mesenchymal-like states closely resembles the EMT program which in itself is associated with the acquisition of stem cell properties (76). Thus, the biology of TMEM is strongly linked with stem and EMT programs, as well as with cancer dissemination. These data further support the use of TMEM score as a prognosticator of distant recurrence of breast cancer (46, 48, 55).

Our findings have potential implications for the prognosis of distant recurrence in breast cancer patients with localized disease, as well as for the prognosis of disease progression and treatment of breast cancer patients with metastases. In terms of prognostication, the association of CSCs with TMEM doorways, resulting in increased CSC-mediated seeding of metastases, provides additional mechanistic insight into the prognostic power of TMEM score. Since TMEM is currently the only clinically validated marker of cancer cell dissemination, it may provide complementary information to current prognostic markers that measure proliferative potential such as OncotypeDX recurrence score (RS). For example, high TMEM score in patients with low OncotypeDX RS could be used to tailor treatment as described in Sparano et al (48). Further, since TMEM score correlates strongly with the percentage of CSCs in tumors as we show here, TMEM score may be used as an indicator of chemoresistance. It is known that CSCs are intrinsically resistant to many therapies (5), and CSC-dependent chemoresistance is believed to be a major cause of metastatic relapse (77). Our observations that tumor-associated macrophages induce the CSC program indicate that chemoresistance may be further increased in situations that promote macrophage influx into tumors. We and others have previously shown that chemotherapy induces macrophage influx (29, 78, 79) and increases the density and the activity of TMEM doorways (29, 80, 81). Consequently, increased macrophage density induces the expression of Mena^INV^ in tumor cells (29) which enhances the transendothelial migration of cancer cells via promoting the assembly and activity of invadopodia (29, 65). Thus, chemotherapy may be impacting two macrophage-associated steps in the dissemination of tumor cells; their stemness and their invasiveness.

Similarly, chemotherapy has previously been associated with enrichment of CSCs in human breast cancer patients (82). To treat metastatic relapse, the inter-related phenomena of chemotherapy-mediated induction of CSC and dissemination need to be addressed. Recent studies have identified drugs that block the recruitment of macrophages to tumors in response to chemotherapy, and block TMEM function (45). Since TMEM doorways are found in metastatic foci in the lymph nodes (83) and in the lungs (33), the same mechanism of dissemination may perpetuate metastases even after the removal of the primary tumor. Thus, it would not be too late to inhibit TMEM function after the removal of the primary tumor because in some patients, cancer cells might have already disseminated from the primary site and formed clinically undetectable micro-metastases, which can be a source of further metastatic cancer cell dissemination via TMEM doorways. Interestingly, the enumeration of CTCs is prognostic even in patients with metastatic breast cancer (84, 85). Our current findings suggest that drugs targeting macrophages might have the important additional benefit of disrupting the induction and dissemination of CSCs (86). Thus, many vexing problems associated with chemoresistance and metastasis may be addressed through this one cellular target.

## GLOSSARY

SORE6>GFP: cancer stem cell reporter construct (Fig S1A)
minCMV>GFP: minCMV control construct
SORE6-GFP: fluorescent protein product of SORE6>GFP construct
SORE6+: same as SORE6-GFP
SORE6-: tumor cells containing the SORE6>GFP construct but not expressing fluorescent protein product (SORE-GFP), due to lack of stem transcription factors activity
CSC: Cancer stem-like cell

## ACKNOWLEDGEMENTS

We thank the Analytical Imaging Facility at Albert Einstein College of Medicine for microscopy help, particularly Dr. Peng Guo for help with Imaris 3D reconstructions; members of the Condeelis, Segall, Cox, Entenberg and Hodgson laboratories for helpful discussions, and Jen Mehalko for cloning expertise. This research was supported in part by CA150344, CA100324, CA216248, F32 CA243350, an IRACDA fellowship, K12 GM102779, the Gruss Lipper Biophotonics Center and its associated Integrated Imaging Program, and SIG #1S10OD019961-01. The research was supported in part by the Intramural Research Program of the NIH grant ZIA BC 005785 to LMW.

## AUTHOR CONTRIBUTIONS

Conceptualization - VPS, BT, YW, JSC, LMW, MHO

Methodology - VPS, BT, YW, GSK, DE, CLD, RJE

Formal Analysis - VPS, BT, YW, GSK, EAX, GK, XY, LMW

Software – VPS, DE

Investigation - VPS, BT, YW, GSK, EAX, LB, AC, CLD, RJE, GK

Writing - VPS, BT, YW, DE, WG, JSC, LMW, MHO

Funding Acquisition - JSC, LMW, MHO, DE

Resources - JGJ, EG, NA, SR, GB, EA, CRS, DE, SG

Supervision - MHO, JSC, LMW, DE

## MATERIALS AND METHODS

### Cell Culturing

The MDA-MB-231 human breast cancer cell line was obtained from ATCC, and the identity of the line was re-confirmed by STR profiling (Laragen Corp.), after expansion and passaging. The human breast cancer stem reporter cell line, td-tomato MDA-MB-231 SORE6>GFP, and the vector control cell line, td-tomato MDA-MB-231 minCMV>GFP, were maintained in 10% FBS in DMEM. The MDA-MB-231-LM2 subline (87) was obtained from Dr. Joan Massague, Sloan Kettering Institute; the Met-1 cell line derived from an MMTV-PyVT mouse mammary tumor (35) was obtained from Dr. Alexander Borowsky, University of California, Davis; the 4T1 metastatic mammary cancer cell line was obtained from Dr. Fred Miller, Karmanos Institute, Detroit. All were maintained in DMEM, 10% FBS. The BAC1.2F5 macrophage cell line was maintained in 10% FBS in α-MEM with 3,000 unit/ml CSF-1. Cryopreserved human primary monocyte-derived macrophages that had been isolated from peripheral blood monocytes and polarized to an M2 phenotype by 4-5 days culture with 10% FBS + 50ng/ml CSF-1 + 10ng/ml IL4 were obtained from StemExpress, Folsom, CA (Cat# PBMAC001.5C) and were thawed and used upon receipt. Human Umbilical Vein Endothelial Cells (HUVECs) (Lonza, Walkersville, MD, USA Lot # 0000396930) were grown in EGM-2 SingleQuot Kit media (Lonza) and used at passage 4-6. All cells were maintained at 37°C in a 5% CO2 incubator, and were shown to be mycoplasma-free (Sigma LookOut Mycoplasma PCR detection kit, Cat# MO0035-1KT).

### Design of cancer stem cell reporter

The original SORE6+ cancer stem cell biosensor consisting of six repeats of a composite SOX2/OCT4 binding element coupled to a minimal CMV promoter driving the expression of a destabilized form of copepod GFP (dsCopGFP) has been described previously (13). A second-generation version of this reporter was generated for these studies. Since the dsCopGFP fluorescence is lost on formalin fixation, the new biosensor has an N-terminal 3x FLAG epitope tag on the dsCopGFP to enable immunohistochemical detection in formalin-fixed paraffin-embedded tissue, and the puromycin drug selection cassette has been replaced with an expression cassette for a C-terminally truncated CD19 marker to enable rapid FACS selection of transduced cells. New constructs were made by Gateway multisite recombinational cloning. The SORE6 enhancer, minimal CMV promoter, tagged destabilized fluorescent protein and SV40-driven truncated CD19 selection marker elements were separately cloned and sequence-verified before Gateway assembly into pDest-412, a lentiviral destination vector based on the pCDF lentiviral backbone to generate the stem cell biosensor (SORE6>GFP). A parallel construct (“minCMV”) lacking the SORE6 enhancer element was generated as a non-specific background control for FACS gating and fluorescent image thresholding. The dsCopGFP protein has a half-life of 1-2 hours, which is similar to the 1.5 hour half-life reported for the OCT4 protein in P19 mouse teratocarcinoma cells (88). Since a small reduction in OCT4 protein levels in sufficient to alter the balance between self-renewal and differentiation (89), the SORE6 reporter kinetics are tuned to parallel closely the dynamics of the stem cell phenotype. The second generation SORE sensor behaves identically to the first generation sensor that was extensively validated by us in multiple breast cancer models (13), and used by others (90–96).

### Tumorsphere assays

MDA-MB-231 cells transduced with the SORE6>GFP stem cell reporter were FACS sorted into SORE6+ and SORE6-populations using cells transduced with minCMV>GFP construct as a gating control. 5000 cells/well were seeded into Costar 24-well Ultralow attachment plates (Corning Cat#3473) in DMEM/10% FBS. After 10 days, tumorspheres were quantitated from images captured using an EVOS FL Auto2 Imaging system and binned by tumorsphere diameter. To address the functionality of SORE6+ cells generated by co-culture with macrophages, 20,000 MDA-MB-231 cells were culture with or without 40,000 BAC cells in 24-well plates for 24h in BAC culture medium. After 24 hours, tumor cells were harvested by brief trypsinization (30 seconds) which selectively removes tumor cells while the BAC cells remain adherent. The harvested tumor cells were seeded at 1500 cells/well in Ultralow attachment plates in DMEM/10% FBS and assessed for tumorsphere formation after 10 days as above.

### Taxol treatment

Unsorted MDA-MB-231 cells containing the SORE6>GFP reporter were seeded into 24-well plates and grown for 24h. Cells were then treated with DMSO vehicle or the indicated concentrations of Paclitaxel (time t=0h) and imaged every 6h using the Incucyte S5 Live Cell imaging system. SORE6-positive cells were quantitated as GFP+ cells and SORE6-negative cells were quantitated as GFP-cells within the same culture, using a parallel culture with the minCMV>GFP construct to set the threshold for positivity.

### qRT-PCR for stem cell transcription factors

RNA was prepared from freshly sorted cells using the Trizol Reagent (Ambion) and cDNA was synthesized using SuperScriptTM III First-Strand Synthesis System (ThermoFisher Scientific) as per manufacturer’s instructions. qRT-PCR was performed with Brilliant II Ultra-Fast SYBR® Green QPCR Master Mix (Agilent) using a Bio-Rad CFX96 Real-Time Detection System. Fold gene expression was calculated using the delta CT method formula: 2^-(DCt). PPIA was used as the reference transcript for normalization. Primer pairs were as follows:

NANOG F: 5’-GTTCTGTTGCTCGGTTTTCT

NANOG R: 5’-TCCCGTCTACCAGTCTCACC

OCT4 F: 5’-AGTCTGGGCAACAAAGTGAGA

OCT4 R: 5’-AGAAACTGAGGAGAAGGATG

SOX9 F: 5’-GTACCCGCACTTGCACAAC

SOX9 R: 5’-TCTCGCTCTCGTTCAGAAGTC

SOX2 F: 5’-TAAATACCGGCCCCGGCGGA

SOX2 R: 5’-TGCCGTTGCTCCAGCCGTTC

SNAI1 F: 5’-GCTGCAGGACTCTAATCCAGA

SNAI1 R: 5’-ATCTCCGGAGGTGGGATG

PPIA F: 5’-GTCAACCCCACGTGTTCTT

PPIA R: 5’-CTGCTGTCTTTGGACCTTGT

### Animal models

All procedures were conducted in accordance with the National Institutes of Health regulations and approved by the Albert Einstein College of Medicine animal use committee. The stem reporter cells tdTomato MDA-MB-231 SORE6>GFP or the reporter control cells tdTomato MDA-MB-231 minCMV>GFP were injected into the mammary fat pad of SCID mice (NCI) as previously described (18). ECFP macrophage/Rag2 knockout mice (Rag2KO Macblue) were generated from the crossing B6.129S6-Rag2^tm1Fwa^ N12(Rag2 - Model RAGN12, Taconic) with Tg(Csf1r*-GAL4/VP16, UAS-ECFP)^1Hume/J^ (Stock No: 026051, the Jackson Laboratory). Tumor cells were orthotopically injected with 50% Matrigel (BD Biosciences, cat # 354234) into the fourth mammary fat pad of Rag2KO MacBlue mice. For drug treatment (Clodronate, DAPT) experiments, animals were randomly assigned to the drug-treated or vehicle-treated groups. All subsequent analyses were performed on randomized samples in a user-blinded fashion.

### In vivo limiting dilution assay for stem cell reporter validation

To validate the newly modified version of the reporter, MDA-MB-231 cells expressing the tdTomato volume marker were transduced with the new SORE6>dsCopGFP construct. In parallel, cells were transduced with the matched minCMV>dsCopGFP as a gating control. Transduced cells were first selected by FACS sorting for the CD19 selectable marker to ensure presence of the reporter in all cells. CD19+ cell populations that were either SORE6+ (CSCs) or SORE6- (non-CSCs) were generated by FACS sorting for GFP. SORE6+ and SORE6-populations were then orthotopically implanted into the #2 and #7 mammary fat pads of 6-week old female SCID/NCr mice in 0.1ml 50% Matrigel. Tumor formation was monitored by regular palpation and caliper measurements, and confirmed at necropsy. Cancer stem cell frequency was calculated from tumor incidence at day 98 post-inoculation, using the ELDA software tool (http://bioinf.wehi.edu.au/software/elda/). 95% confidence intervals for the cancer stem cell frequency are 1 in 5608 to 1 in 1577 for SORE6+ cells, and 1 in 71505 to 1 in 7518 for SORE6-cells. It should be noted that this assay is useful for assessing the relative tumor initiating ability (and hence CSC representation) of two cell populations, but that the absolute CSC frequencies calculated by this approach strongly under-represent the true absolute frequencies. This is because the efficiency of experimental tumor initiation is strongly affected by multiple parameters, such as degree of immunodeficiency, mouse strain background, site of tumor cell implantation and others, as documented in the literature (97).

### Fluorescence-Activated Cell Sorting (FACS) and flow cytometry

For cultured cells, cells were typsinized, pelleted, and resuspended in 2% BSA/PBS as a single cell suspension for FACS sorting. Tissues from tdTomato MDA-MB-231 SORE6>GFP or tdTomato MDA-MB-231 minCMV>GFP primary tumors were washed once with PBS, diced, suspended in PBS, passed through a 70μm cell strainer, and centrifuged down to cell pellets. Pellets were resuspended in PBS, filtered with a 40μm cell strainer, spun down, resuspended in 200μL of 2% BSA/PBS, and stained with APC-lineage antibody cocktail (BD Pharmingen, APC mouse lineage antibody cocktail, Cat# 51-9003632) at 5μL antibody/million cells for 30 minutes at room temperature. After staining, the cells were washed twice with PBS and resuspended in 2% FBS/PBS for FACS sorting. Comparing with isotype control, APC negative cells are the tumor cells for analysis. When sorting the SORE6+ (GFP+) and SORE6-(GFP-) cells from tdTomato MDA-MB-231 SORE6>GFP tissue, tdTomato MDA-MB-231 minCMV>GFP was always used as the GFP gating control. Cell surface marker expression in the SORE6+ and SORE6-compartments of MDA-MB-231 cells cultured in vitro or recovered from primary tumors was assessed by flow cytometry following staining with Brilliant Violet 421^TM^ anti-human CD133 (Biolegend Cat#372807), or APC anti-human CD44 (Biolegend Cat#338805) and CD24-Brilliant Violet 711^TM^ anti-human CD24 (Biolegend Cat#311135), all used at 5μl antibody per million cells. Flow cytometry of stained cells was performed on a BD LSRFortessa SORP1 (BD Biosciences) and data were analyzed using FlowJo software. Cells in the TdTomato positive gate (all tumor cells) were further gated into SORE6+ and SORE6-fractions using the SORE6 stem cell sensor, for assessment of cell surface marker expression in the two compartments.

### Fixed Frozen Tissue Immunofluorescence Staining and Imaging

Primary tumors and the inflated lungs from tdTomato MDA-MB-231 SORE6>GFP or tdTomato MDA-MB-231 minCMV>GFP xenograft mice were collected. The tissues were fixed in 5% neutral buffered formalin and 20% sucrose in 4°C for 2 days before being embedded and frozen in OCT compound. The fixed frozen sections were brought in room temperature for 30 min, treated with −20°C cold acetone for 10 minutes, washed with PBS, outlined with PAP pen, stained with DAPI for 10 minutes, washed with PBS, and mounted on a coverslip for imaging. In the case of immunofluorescent staining, before the DAPI staining was performed, each section was blocked with 1% BSA, 10% FBS in PBS for 1 hour at room temperature, incubated with anti-IBA1 antibody (1:200, Anti-IBA1, Rabbit, Wako) in blocking buffer for 1 hour, washed with PBS three times, incubated with secondary antibody (1:200) for 1 hour, and washed two times as validated previously (45). The dried stained slides were imaged with the DeltaVision microscope. tdTomato-MDA-231 minCMV>GFP tissue slides were used for thresholding the images. Images were analyzed in ImageJ.

### In vivo time-lapse intravital imaging and analysis

A mammary imaging window was implanted into mice with an MDA-MB-231 primary tumor expressing tdTomato + SORE6>GFP or tdTomato + minCMV>GFP, and intravital multiphoton imaging was performed on a custom-built two-laser multiphoton microscope as described earlier (98, 99). Mice were anesthetized with 1-2% isofluorane and kept at physiological temperatures on the microscope stage with the built-in environmental heat enclosure and vitals were monitored with pulse oximeter (PhysioSuite, Kent Scientific). All images were captured using a 25x 1.05 NA water-immersion objective.

For SORE6>GFP stem biosensor imaging, GFP background was set using tumors expressing the minCMV>GFP control construct (Figure S3). All subsequent SORE6>GFP stem biosensor imaging was performed at this laser power and GFP gain setting. Far-red Q dots (Thermo Fisher Scientific, cat# Q21061MP) were used to visualize the blood vasculature. Long time-lapse z-stack imaging was performed by imaging multiple fields (512 x 512 μm^2^ or 341 x 341 μm^2^) with a 5 μm z-step size.

Motility analysis (Fig 2B) in the intravital movies was performed as described before (18). The majority of tumor cells in the field were stationary and not included in the analysis. Only motile tumor cells, consistent with (18), Fig S2, were analyzed for speed calculations. Individual 16-bit TIFF images were assembled into a CZT hyperstack using a custom ImageJ macro. Temporal drift in multiple-channel time-lapse hyperstacks was corrected using HyperStackReg plugin (100). For single cells SORE6- and SORE6+ cell motility analysis, cells were manually tracked using TrackMate plugin (101). Additionally, we measured tissue drift in our intravital movies and found that tissue drift after post-acquisition drift correction is minimal (∼0.02 μm/min, Fig S4D, movie 8) and does not contribute to the cell migration on the time scale of the movies.

### Macrophage depletion using Clodronate

A chronic mammary imaging window was implanted into Rag2KO Macblue mice with an MDA-MB-231 primary tumor expressing tdTomato + SORE6>GFP, and intravital multiphoton imaging of multiple 512 x 512 μm^2^ fields were performed on three consecutive days (day0, day1 and day2) using an in vivo microcartography technique (33) on a custom-built two-laser multiphoton microscope as described above. Mice were injected intraperitoneally with 200 μl of either control PBS liposomes or Clodronate liposomes (Encapsula Nano Sciences, SKU# CLD-8901) on day0 and day1 after intravital imaging. The 3D objects counter plugin in Fiji was used to quantify macrophage voxels in the 3D hyperstacks. Macrophage density was calculated as the voxels covered by macrophages divided by total voxels in the field. SORE6+ cells were manually counted in each 3D hyperstack. In PyMT model, mammary tumors from PBS liposomes or Clodronate liposome treated mice were extracted. FFPE tumor sections were stained with Iba-1 IHC and macrophage density was calculated by the area covered by macrophages (Iba-1 IHC, brown staining) divided by the total area of the field. For Sox9^Hi^ quantification, FFPE tumor sections were stained with Sox9 antibody (EMD Millipore, cat# ab5535) and Sox9 channel was thresholded for the fluorescence intensity range 80-255, such that Sox9^Hi^ tumor cells were ∼ 5% of the total number of tumor cells in the control tissue. Same threshold was applied to the clodronate tissue.

### In vivo cancer stem cell identification in fixed tissue with anti-FLAG IF staining

Primary MDA-MB-231 xenograft tumors from SCID and nude mice, expressing tdTomato + SORE6>GFP or tdTomato + minCMV>GFP, were excised, fixed in formalin, embedded in paraffin (FFPE), and 5 μm sections were cut using a cryostat. For IF staining for the FLAG tag on the dsCopGFP, sections were deparaffinized, followed by antigen retrieval in 1 mM EDTA, pH 8 for 20 min in a conventional steamer and 20 min incubation at room temperature. Tissues were washed 3×5 min and incubated in blocking buffer (1% BSA + 5% goat serum) for 1.5 - 2 hours at the room temperature, followed by rat anti-FLAG at 1:1000 (BioLegend, cat# 637301) incubation overnight at 4 deg C and secondary antibody goat anti-rat Alexa 647 for 1 hour at room temperature. The FLAG channel in the minCMV>GFP tissue was used to set the background in SORE6>GFP tissue to identify single stem cells. For *in vivo* characterization of FLAG antibody (Fig. S2B), SORE6>GFP sensor expressing MDA-MB-231 primary tumor FFPE sections were treated with 1X citrate pH 6.0 buffer (PerkinElmer, cat# AR6001) and stained with chicken anti-GFP (Novus, cat# NB100-1614) and rat anti-FLAG antibodies.

### In vivo invadopodia and degraded collagen IF staining, imaging and quantification

In live tissue (Fig 2C, 2D, 2E), invadopodia are identified morphologically as thin oscillatory protrusions with 8-10 min cycle (19, 24). For invadopodia identification in fixed tissue, primary MDA-MB-231 xenograft tumors were excised from SCID mice expressing tdTomato + SORE6>GFP or tdTomato + minCMV>GFP. Formalin-fixed paraffin-embedded (FFPE) tissues were deparaffinized, followed by antigen retrieval in 1 mM EDTA, pH 8 for 20 min in a conventional steamer. Tissues were incubated in blocking buffer (1% BSA + 5% goat serum) for 2 hours at the room temperature, followed by primary antibodies overnight at 4 deg C and secondary antibodies for 1 hour at room temperature. In fixed tissue (Fig 2G), invadopodia are identified molecularly by Cortactin+Tks5 co-localized dots (19, 27). Cortactin+Tks5 co-localized puncta within the cell boundary marks invadopodia in these cells, consistent with published literature (26, 27, 102). We did not use phalloidin staining, since F-actin is present in other cellular structures (e.g. filopodia, focal adhesions etc.), and is not a specific marker of invadopodia. Degraded collagen was detected with C1,2C (Col 2 3/4C_short_) rabbit antibody (Ibex Pharmaceuticals) (22). Degraded collagen staining co-localizing with cortactin, one of the invadopodia marker, identifies mature invadopodia (i.e. actively degrading shown by white arrow in Fig 2J). Stem cells were identified with rat anti-FLAG antibody as described above.

The slides were imaged on a 3D Histech Pannoramic P250 digital whole slide scanner, using a 40x 0.95 NA air objective lens. Digital whole slide scan tissue images were opened in CaseViewer v2.1. FLAG channel intensity in minCMV>GFP tissue was used to set the background in SORE6>GFP tissue to identify single stem cells, and 5-6 random regions of interest (ROIs) at 40X from non-necrotic areas were saved. Images were opened in ImageJ and a custom ImageJ macro was used for dividing each image into stem cell and non-stem cell areas. Cell areas were defined as the circular 36 μm diameter ROI around each single stem cell (3 x diameter of a single cell, ∼ 12 μm to capture all the extending invadopodia protrusions and their collagen-degrading activity from each cell). Invadopodia were identified as cortactin and Tks5 containing puncta and normalized to the size of stem and non-stem cell areas. Accordingly, collagen degradation was calculated as the Col3/4-positive area (%) in stem and non-stem cell areas as described above. In addition, a higher resolution analysis, to assign the origin of the cortactin and Tks5 containing puncta signal to SORE6+ or SORE6-cells, was done by reanalyzing random fields in Fig 2I and 2L by drawing the boundary of FLAG+ cells manually (as shown in figure panels 2G and 2J) to define stem cell and non-stem cell areas. Reanalyzed results (supplementary figure S4F, S4G) led to the same conclusions as that obtained using the automated 36 μm ROI method (figures 2I and 2L), i.e. compared to non-stem cells, stem cells have significantly more invadopodia and higher degradation activity in vivo.

### Cancer stem cell distance analysis relative to TMEM, and blood vessels without TMEM, in fixed tissue in vivo

Sequential sections from primary MDA-MB-231 xenograft tumors from SCID mice, expressing tdTomato + SORE6>GFP or tdTomato + minCMV>GFP, were stained for TMEM IHC (Mena, Iba-1 and CD31) (29, 47), and anti-FLAG to identify SORE6+ stem cells. TMEMs were identified manually by a pathologist. Stem cells were identified by thresholding the anti-FLAG image based on the threshold value obtained from the anti-FLAG stained minCMV>GFP control tumor section. TMEM IHC and FLAG images were aligned in ImageJ using *Landmark Correspondences* plugin. A custom-written ImageJ macro was used to calculate the distance of each stem cell in the field to its nearest TMEM. For non-stem cell distance with the nearest TMEM in the field, DAPI channel was thresholded followed by watershed algorithm to find non-stem cells. Distance of each non-stem cell to its nearest TMEM was calculated using a custom-written ImageJ macro as above. The distance of CSCs to TMEM was calculated in reference to TMEM, not the entire length of the blood vessel that contains TMEM (TMEM and blood vessel without TMEM schematics shown in Figs 5C and 5E). Only cells which are inside or touching the 60 μm diameter TMEM circle is the region of interaction (ROI) of TMEM with surrounding cells (data point 0 μm in Figs 5D, G). To exclude the possibility of CSCs being near TMEM by chance, CSC distance histograms (Figs 5D, F, G, H) were normalized by distances of all tumor cells in the field (DAPI staining) to nearest TMEM (Figs 5D, G) or blood vessel without TMEM (Figs 5F, H). Similarly, stem cell distance analysis with nearest blood vessel without TMEM was done by thresholding the blood vessel channel and removing blood vessels which contained TMEM. Distance histograms were analyzed and plotted in Excel.

### In vitro tumor cell-macrophage co-culture assay

Tumor cells (td-tomato MDA-MB-231 SORE6>GFP) were plated alone or with macrophages (BAC1.2F5) or HUVEC (Lonza, cat# CC-2517) at a 1:5 ratio (5,000 tumor cells and 25,000 macrophages or HUVEC) in a 35 mm glass-bottom dish (Ibidi, cat# 81156) and allowed to adhere overnight. Next day, cell media was replaced with imaging media (L-15 + 10% FBS) and cells were imaged live on an epi-fluorescence DeltaVision microscope (GE) with a 20X objective and CoolSNAP HQ2 CCD camera. For live imaging, tumor cells were plated overnight. Next day, media was replaced with imaging media, followed by macrophage addition at the beginning of live imaging. To identify stem cells, control td-tomato MDA-MB-231 minCMV>GFP cells plated under identical conditions were used to set the background GFP level. For Notch1 inhibition, 10 μM DAPT (Sigma, cat# D5942) or vehicle control DMSO was added to the co-culture dish. For Notch activation, 50 μM Jagged1 (AnaSpec, cat# AS-61298) and its control, Jagged1 scramble (AnaSpec, cat# AS-64239) was added to the co-culture dish. To confirm the co-culture observations independently in different tumor models using different analysis procedures, 10,000 4T1 cells transduced with SORE6>dsEGFP were co-cultured with 50,000 BAC1 cells in BAC medium (α-MEM, 10% FBS, 10ng/ml CSF1) for 4 days, and then trypsinized for flow cytometry analysis to identify the SORE6+ cells as % total tumor cells using the minCMV>dsEGFP as a gating control. BAC1 cells were gated out using the F4/80 macrophage marker. Similarly, MDA-MB-231-LM2 cells expressing a GFP volume marker were transduced with a SORE6>dsmCherry stem cell sensor. 10,000 tumor cells were cocultured with 20,000 BAC cells in BAC medium for 36 hours and then analyzed by FACS to determine SORE6+ CSCs (mCherry+ cells) as % total tumor cells (identified by GFP volume marker). To assess the requirement for direct contact, BAC cells were seeded into Millicell Hanging Cell Inserts (Millipore Sigma, 3μm pore size PET membrane), with tumor cells in the well below, so that direct cell contact was prevented but secreted factors could be freely exchanged between the two cell types.

### Notch1 signaling inhibition in vitro using siRNA

AllStars Neg. Control siRNA (1027281) was from Qiagen, On-target plus human Notch1 siRNA smartpool was from Dharmacon (Thermo Scientific, cat# L-007771-00-0005). A total of 1.5×10^6^ tumor cells (td-tomato MDA-MB-231 SORE6>GFP) were transfected with 10 μl of 20 μM siRNA stock solution using Nucleofector Kit V from Lonza (cat# VCA-1003) for 48 h, as described previously (32). Notch1 knockdown was confirmed using western blot with the following antibodies: rabbit anti-Notch1 (Cell Signaling, cat# 3608), rabbit anti-Notch2 (Cell Signaling, cat# 5732) and mouse anti-GAPDH (Abcam, cat# ab8245). At 36 h post siRNA transfection, tumor cells were plated with macrophages (1:5 ratio) overnight. Next day, cells were imaged live on DeltaVision microscope as described above.

### Notch signaling inhibition in vivo using DAPT

DAPT (Sigma-Aldrich Cat# D5942) was reconstituted in 100% ethanol to a stock concentration of 20 mg/ml, then further diluted in corn oil to a final concentration of 2 mg/ml. Eight-week-old PyMT mice bearing palpable tumors and separate cohort of SCID mice with MDA-MB-231 xenograft tumors expressing tdTomato + SORE6>GFP or tdTomato + minCMV>GFP were given daily intraperitoneal injections of 10 mg/kg DAPT or vehicle control (1:10 ethanol in corn oil) for 14 days. On day 15, the primary tumors, lungs, and duodenums were collected from the mice and fixed in 10% formalin. Mice were weighed on day 1 and day 15 to determine that no significant weight loss was suffered due to the DAPT treatment. Duodenums were stained using the Periodic acid-Schiff (PAS) staining and saw an increase in goblet cell hyperplasia in the intestinal crypts (Fig S6J), as consistent with successful in vivo Notch signaling inhibition (103–105).

In PyMT model, primary FFPE tumor sections were stained with Iba-1 antibody and macrophage density was calculated by the area covered by macrophages divided by the total area of the field. For Sox9 quantification, FFPE tumor sections were stained with Sox9 antibody and Sox9Hi tissue coverage area was quantified. In MDA-MB-231 model, FFPE lung sections were stained with FLAG antibody to identify SORE6+ CSCs.

### Time Lapse Imaging of in vitro tumor cell-macrophage co-culture

In vitro time lapse movies of co-cultured cells were taken with a DeltaVision microscope at 2 minute intervals for 16 hours, at 20x magnification. To avoid phenol red autofluorescence, the cells were plated into L15 culturing media. Fixed culture plates were also imaged at 20x; these images were used to determine percentage of SORE6+ cells by GFP intensity, thresholded based on the background GFP expression in vector control cell, td-tomato MDA-MB-231 minCMV>GFP. For cell fate-mapping in co-culture experiment shown in Figure 4F, 25,000 tumor cells (MDA-MB-231 SORE6>GFP) were plated in BAC culture media in 35 mm glass-bottom dish (Ibidi, cat# 81156) and allowed to adhere overnight. Next day, media was changed to L-15+10% FBS+CSF-1 media and cells were imaged live every 10 min on an epi-fluorescence DeltaVision microscope (GE) with a 20X objective and CoolSNAP HQ2 CCD camera. After 30 min of imaging, 125,000 BAC cells were added to tumor cells in the dish on the imaging stage and imaging was continued for 30 hours. Time-lapse movies were analyzed by identifying SORE6-cells in the frames before macrophage addition, based on the GFP threshold set by MDA-MB-231 mCMV>GFP fluorescence. SORE6-cells were manually tracked frame-by-frame for macrophage touch or no touch in the phase channel time-lapse and the corresponding GFP fluorescence increase or no change in the GFP channel time-lapse.

For cell origin mapping in co-culture experiments as shown in Fig 4G, MDAMB231 tumor cells were FACS-sorted to give a starting population that contained ∼2% SORE6+ cells so as to simplify the mapping process. 20,000 tumor cells were co-cultured with or without 30,000 BAC cells in 24-well plates in BAC medium, and live imaged every hour starting at 6h after plating, using an Incucyte S3 Live-Cell Analysis System at 10x magnification (Essen BioScience). Cells that were SORE6+ at t=30h were origin-traced back to their starting state at t=6h to determine whether they originated from pre-existing SORE6+ cells or from direct induction of a stem phenotype in SORE6-cells. The fraction of SORE6+ cells at 30h that originated from SORE6-cells at 6h was assessed by origin-mapping 32 SORE6+ cells/well from four wells/condition, for a total of 128 SORE6+ cells/condition.

### Gelatin Degradation Assay and In Vitro Immunofluorescent Staining

Invadopodia matrix degradation assay was performed as described previously (21). SORE6+ and SORE6-cells were FACS sorted and plated on fluorescent Alexa-405-labeled gelatin matrix coated plates overnight. Cells were fixed with 3.7% paraformaldehyde for 15 minutes and permeabilized with 0.1% Triton X-100 for 5 minutes. Cells were blocked in 1% BSA + 5% FBS for 1 hour, incubated with primary antibodies; anti-cortactin (1:400) and anti-Tks5 (1:100 dilution) for 1 hour, followed by incubation with appropriate secondary Alexa antibodies (1:400) for 1 hour. Cells were images on DeltaVision widefield fluorescence microscope. Images were processed in ImageJ.

### Intravasation assay (CTC analysis)

About 1 ml of blood was drawn from the right heart ventricle of anesthetized mice bearing an ∼1.5 cm diameter primary tumor. Blood was immediately fixed in 5% neutral buffered formalin (NBF) at room temperature for 15 min., then washed with 10 ml PBS 3 times by centrifugation at 1000 rpm for 5 min. The pellet was resuspended in PBS / 0.1% Triton-X for 1 min, washed once, and stained with DAPI. The final pellet was resuspended in 300 μl PBS, dropped onto 35 mm glass-bottom μ-Dishes (Ibidi, cat #81156) and left in 4°C overnight before imaging on an epi-fluorescence DeltaVision microscope (GE). Quantification of SORE6+ CSCs was done in ImageJ. CTCs from minCMV>GFP control reporter tumor mice were used for setting the GFP threshold.

### Live lung imaging

SCID mice bearing orthotopic MDA-MB-231-SORE6+ tumors underwent placement of permanent lung imaging windowas previously described (33). Large volume high resolution intravital (LVHR-IVI) imaging (98) was performed through the window 24 hours post-implantation (Day 0) to capture the architecture of lung vasculature. Vasculature was labeled with an intravenous injection of 50μL of a fluorescently tagged dextran (50μg/mL, 10kD Cascade Blue Dextan). 24 hours later (Day 1) LVHR-IVI imaging was repeated to capture disseminated tumor cells that arrived in the lung vasculature. SORE6- and SORE6+ cells were distinguished by the presence or absence of the green GFP signal in addition to the red tdTomato cell volume marker present in all tumor cells.

### Ex vivo lung processing

SCID mice, bearing 1-1.5 cm orthotopic MDA-MB-231-SORE6+ tumors, were euthanized and lungs were inflated with 5% formalin injected into trachea. Lungs were removed and incubated in 5% formalin/20% sucrose solution overnight at 4 deg C. Next day, lungs were embedded in OCT compound and 7 μm thick sections were cut. Slides containing lung sections were equilibrated at room temperature for 10 min, permeabilized in cold acetone for 10 min and washed once with PBS. Slides were then mounted with mounting media containing DAPI (Vector Laboratories, Burlingame, CA). Slides were imaged on P250 scanner and images were acquired in DAPI, FITC (SORE6-GFP) and TRITC (tdTomato) channels.

### Human breast cancer tissue collection

Breast cancer tissue was collected from 49 patients under the Albert Einstein College of Medicine/ Montefiore Medical Center Institutional Review Board protocol approval. Cancer cells were collected by fine needle aspiration (FNA) from unfixed mastectomies and lumpectomies and used for the assessment of stem cell marker expression as described below. FNA primarily collects loose tumor cells, with very few macrophages and no endothelial cells and incurs minimal tissue damage (106). Briefly, five to ten FNA aspiration biopsies per tumor were performed on at least 3 areas of grossly visible tumor using 25-gauge needles. The adequacy of the sample was assessed by the standard Diff-Quick protocol (107). Only samples composed of at least 90% malignant epithelial cells, as determined by standard pathologic characteristics were used for the analysis (108). After cell collection by FNA, the remaining tissue was formalin fixed, embedded in paraffin (FFPE) and sent for routine pathological analysis. FFPE tissue blocks with representative tumor were used for triple immunohistochemical (IHC) analysis of TMEM sites. This approach allowed us to assess the percentage of cancer cells expressing stem cell markers and the density of TMEM doorways from the same samples (Fig 7B).

### Tissue Selection for TMEM Staining and Scoring

At the time of routine microscopic examination of the lesions on which FNA biopsies had been performed, an appropriate area containing invasive cancer suitable for TMEM analysis was identified by low power scanning. The following criteria were used: high density of tumor, adequacy of tumor, lack of necrosis or inflammation, and lack of artifacts such as retraction or folds. TMEM stain is a triple immunostain for predicting metastatic risk in which 3 antibodies are applied sequentially, and developed separately with different chromogens on a Bond Max Autostainer. We used the pan-Mena mouse monoclonal antibody (BD# 610693), CD68 (Dako # M0876) and CD 31 (Dako # M0823). The assessment of TMEM scores was performed using Adobe Photoshop on 10 contiguous 400x digital images of the most representative areas of the tumor. The total TMEM for each image was tabulated, and the scores from all ten images were summed to give a final TMEM density for each patient sample, expressed as the number of TMEM per 10 400x fields (total magnification) (46, 47). Twenty-five randomly chosen cases were each independently scored by 2 pathologists. Since the correlation between the scores read by 2 pathologists was excellent with the correlation coefficient r=0.97, the remaining 24 cases were scored by one pathologist.

### Flow cytometry analysis for CD44+/24-

The cells obtained by FNA were washed with PBS and filtered through 35 μm cell strainer to make it into a single cell suspension (4°C was maintained all the time). The cells were then diluted to 500 cells/μl in PBS to 400 μl and divided into two eppendorf tubes at 200 μl each. In first tube 2 μl each of the isotype controls for CD44 and CD24 were added. Then in second tube each of two antibodies anti-human/Mouse CD44 PE-Cyanine5 and anti-human/Mouse CD24 PE were added. These solution mixtures were kept at room temperature for 20 min in the dark. The samples were then run through GUAVA flow cytometer. The first tube was run to set up the quadrants and antibody isotype controls were used to setup the gain in the machine. Then the samples with antibodies were analysed. Cells present in lower right quadrant showed the % of CD44+/CD24- cell population.

### In situ hybridization for ALDH1 and CD133

In situ hybridization was performed using Cy3 labeled ALDH1 and CD133 probes (Aanera Biotech, Albany, NY), the cell membrane was stained with 5-Hexadecanoylaminofluorescein and the nucleus with and DAPI (Thermo Fisher Scientific, Waltham, MA). The cells were fixed in 3% formaldehyde and permeabilized using 0.1% Triton-100 buffer. 50ng of probe was mixed with 100 μl of hybridization buffer containing 1mg of tRNA, 10mg Dextran Sulphate, 20 μl of deionized formamide, 10ng of N30 oligonucleotide in 2X SSC and hybridized at 37°C in dark in a shaker incubator at 50 rpm. The cells were resuspended in 10% deionized formamide in 2X SSC and incubated at 42°C in dark in a shaker incubator at 50 rpm. Re-suspend cells in wash buffer containing 10ng/μl DAPI for nuclear counterstaining and 200ng of FITC dye an incubate for 5 minutes at 37°C in dark. Reconstitute in 50 μl of Anti-fade Mounting Buffer (Thermo Fisher Scientific, Waltham, MA) and mount in a glass bottom petriplate (MatTek Corporation, Ashland, MA). Imaging was performed for DAPI, FITC and Cy3 channels. Using ImageJ the FITC area labeling the membrane and DAPI stained nucleus area was identified. Cy3 signal was quantified in the donut like cytoplasmic area and was compared to the TMEM values.

### qRT-PCR for ALDH1 and CD133

Total RNA was amplified as previously described (109). Briefly, 1X lysis buffer was added to the cancer cells. Primer mix (50μg/ μl and 100 μg/ μl) was added to the lysis solution. After incubation (80°C for 10 min), RT master mix (5X First strand buffer, DTT, 10 mM dNTP mix, RNase inhibitor, SS III enzyme) was added. This reaction mixture was incubated at 42 °C for 60 min for reverse transcription. cDNA was purified by ZYMO column and 1X wash buffer. Purified cDNA was eluted by nuclease free water. Tailing master mix (10X TdT Buffer, 10 mM dTTP, TdT enzyme) was added to the purified cDNA for TdT tailing. The reaction mixture was incubated for 37 °C for 2 min, 80°C for 10 min and 4 °C for 2 min. Then promoter synthesis master mix (10X Klenows buffer, 50μg/ μl T7 promoter oligo, dNTP mix, Klenow enzyme) was added to the poly A tailed cDNA solution. This reaction mixture was incubated for 22°C for 30 min, hold at 4 °C for T7 promoter synthesis. Finally, in vitro transcription with T7 polymerase was done by T7 high yield RNA systhesis kit (New England Biolabs). Purification of mRNA was done by Qiagen mini column. After eluting the amplified mRNA, was quantified spectrophotometrically using the NanoDrop 2000 spectrophotometer (Thermo Scientific). After first round amplification, 100 ng RNA was taken for second round amplification (same procedure described above). The first Strand cDNA synthesis was carried out by N9, dTVN, Superscript transcriptase II, dNTPs, RNAase, using this 2μg of second round amplified RNA. qPCR was performed with stem cell markers specific primers. The expression of each gene was quantified by using the comparative Ct (ΔCt) method as described in the Assays-on-Demand User’s Manual (Applied Biosystems). The fold values (X) were calculated using the formula: X = 2(-ΔCt), where the data for the sample and sham-treated cell (here, the calibrator being the sham-treated cells) were first normalized against variations of sample quality and quantity. The ΔCt was determined using the formula: ΔC(t) sample = C(t) target gene of sample - C(t) reference gene. reference. The expression of the target genes was normalized to GAPDH.

### Statistical Analysis

GraphPad Prism 7 and Excel were used to generate graphs/plots and for statistical hypothesis testing. Statistical significance was determined by either t-test (normally distributed paired or unpaired dataset), Mann–Whitney test (non-normally distributed unpaired dataset) or Wilcoxon matched-pairs test (non-normally distributed paired dataset). Normality (Gaussian distribution) of each dataset was checked with D’Agostino & Pearson normality test in GraphPad Prism (110). Statistical significance was defined as p-value < 0.05. For the correlation of TMEM density with the proportion of cancer cells expressing stem cell markers we used Pearson’s correlation.

## Supplementary Figures

**Figure S1:**
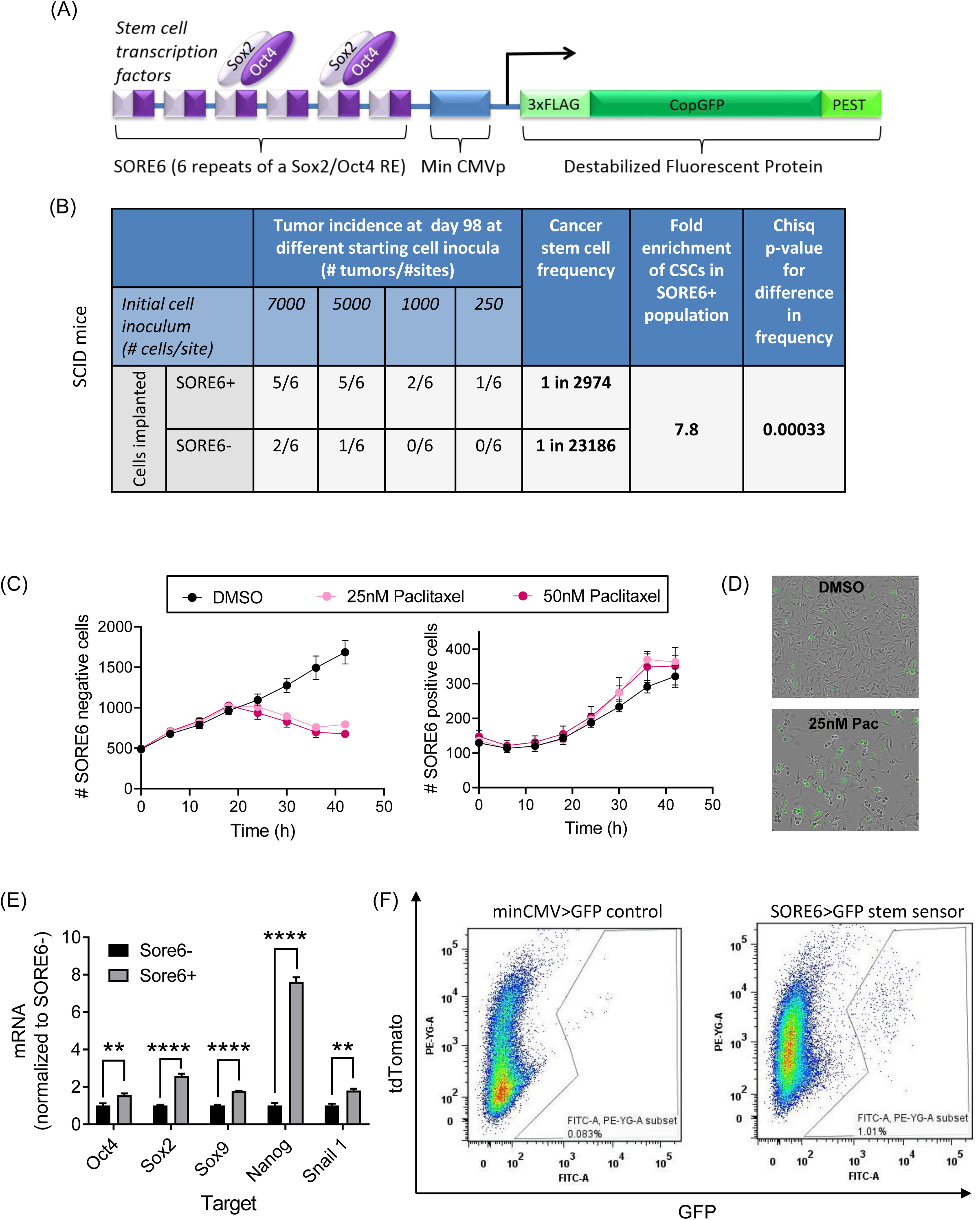
Characterization of SORE6 stemness reporter. (A) SORE6 stem cell reporter schematic showing the SORE6 stem cell-specific enhancer element, consisting of six tandem repeats of a composite response element for Sox2 and Oct4, coupled to a minimal CMV promoter element to provide the TATA box (13). dsCopGFP is a destabilized copepod GFP that has the ornithine decarboxylase PEST sequence added. Three tandem FLAG epitopes were inserted before the dsCopGFP reporter for detection of SORE6+ cells in fixed tissue using FLAG antibody. This stem cell reporter construct is referred to as “SORE6>GFP” throughout. A control construct lacking the SORE6 enhancer element was used for gating and thresholding (minCMV>GFP). (B) In vivo limiting dilution assay showing nearly 8-fold enrichment for tumor-initiating ability of SORE6+ MDA-MB-231 cells compared to SORE6-MDA-MB-231 cells in SCID mice. (C) SORE6-positive cells are resistant to chemotherapeutics. (D) Representative images of the culture at t=42h. (E) qRT-PCR of stem cell and EMT master transcription factor transcripts in MDA-MB-231 cells after FACS sorting into SORE6+ and SORE6-populations. Data are mean ± SEM (n=3); unpaired t-tests. **, p<0.01; ****, p<0.0001 (F) FACS analysis showing that the SORE6 reporter identifies a minority population in MDA-MB-231 tumors in SCID mice of ∼ 1%.

**Figure S2:**
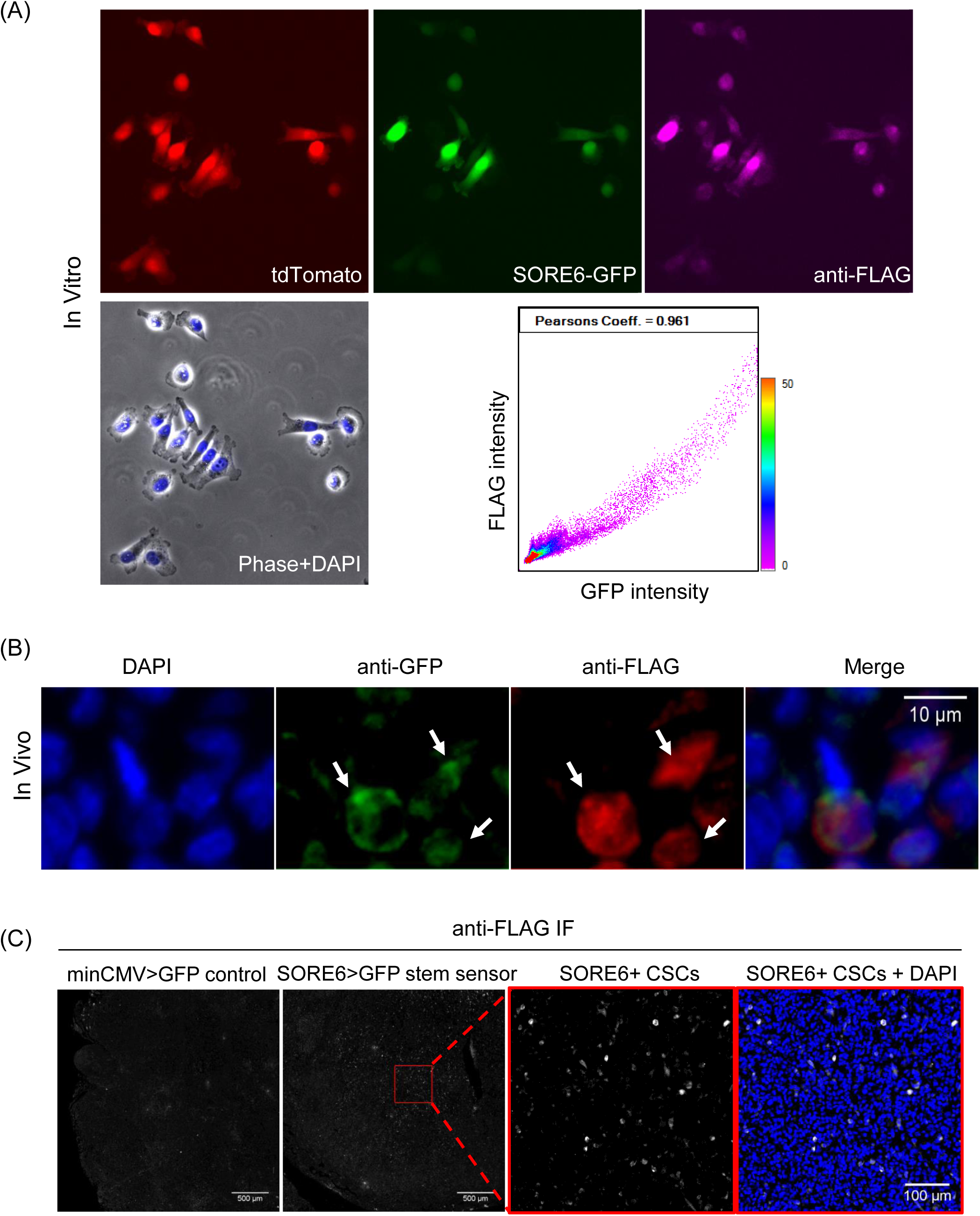
Characterization of FLAG antibody for identifying SORE6+ stem cells. (A) MDA-MB-231 cells expressing tdTomato volume marker and containing the SORE6>GFP reporter construct were plated on glass-bottom dishes, fixed and stained with FLAG antibody to detect the FLAG tag on the dsCopGFP. Antibody staining shows the presence of FLAG in SORE6-GFP positive cells. Scatter plot shows a strong positive correlation between GFP and FLAG signals to identify the SORE6+ tumor cells (Pearson’s coefficient = 0.96). (B) SORE6>GFP sensor expressing MDA-MB-231 primary tumor FFPE sections were stained with anti-GFP and anti-FLAG antibodies and show the presence of FLAG in SORE6-GFP positive cells in vivo (white arrows). (C) *In situ* immunofluorescence anti-FLAG staining identifies minority population of SORE6+ cancer stem cells in fixed tissues in nude mouse host. Left panel: tdTomato-MDA-MB-231 –minCMV>GFP vector control tumor tissue, right three panels: tdTomato-MDA-MB-231-SORE6>GFP tumor tissue. SORE6 expressing CSCs stained with anti-FLAG are shown in white.

**Figure S3:**
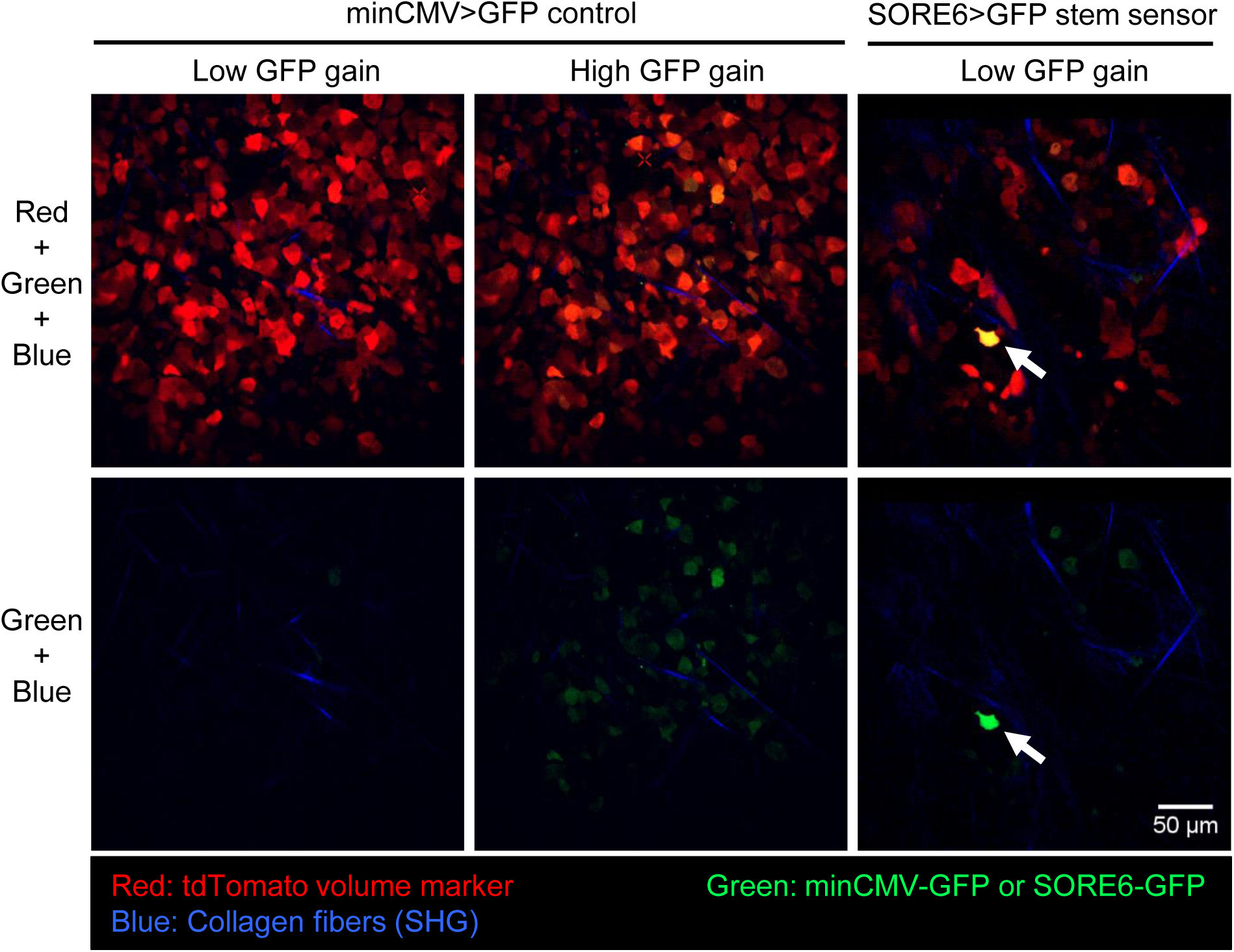
Optimization of imaging conditions for identifying SORE6+ cancer stem cells during intravital imaging in living mouse. Mice with tumor cells expressing minCMV>GFP control plasmid were imaged at low GFP gain settings (left panels) to set the GFP background threshold. High GFP gain image of the same field for minCMV>GFP (middle panels) clearly show low background GFP levels in tumor cells. All intravital imaging in mice was performed at low GFP gain setting and shows minor population of cancer stem cells (right panels, white arrow) in tumors expressing SORE6 reporter.

**Figure S4:**
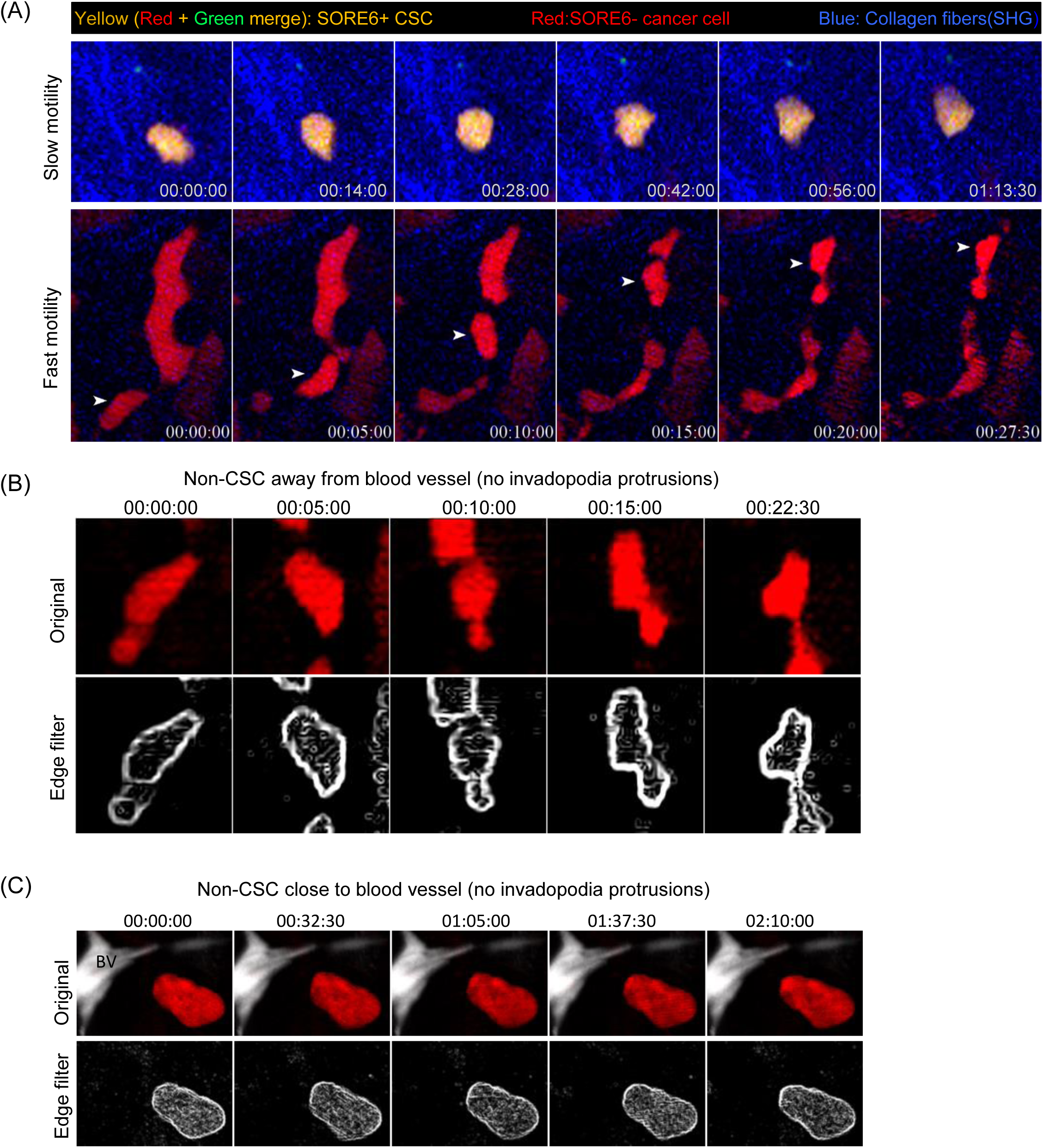
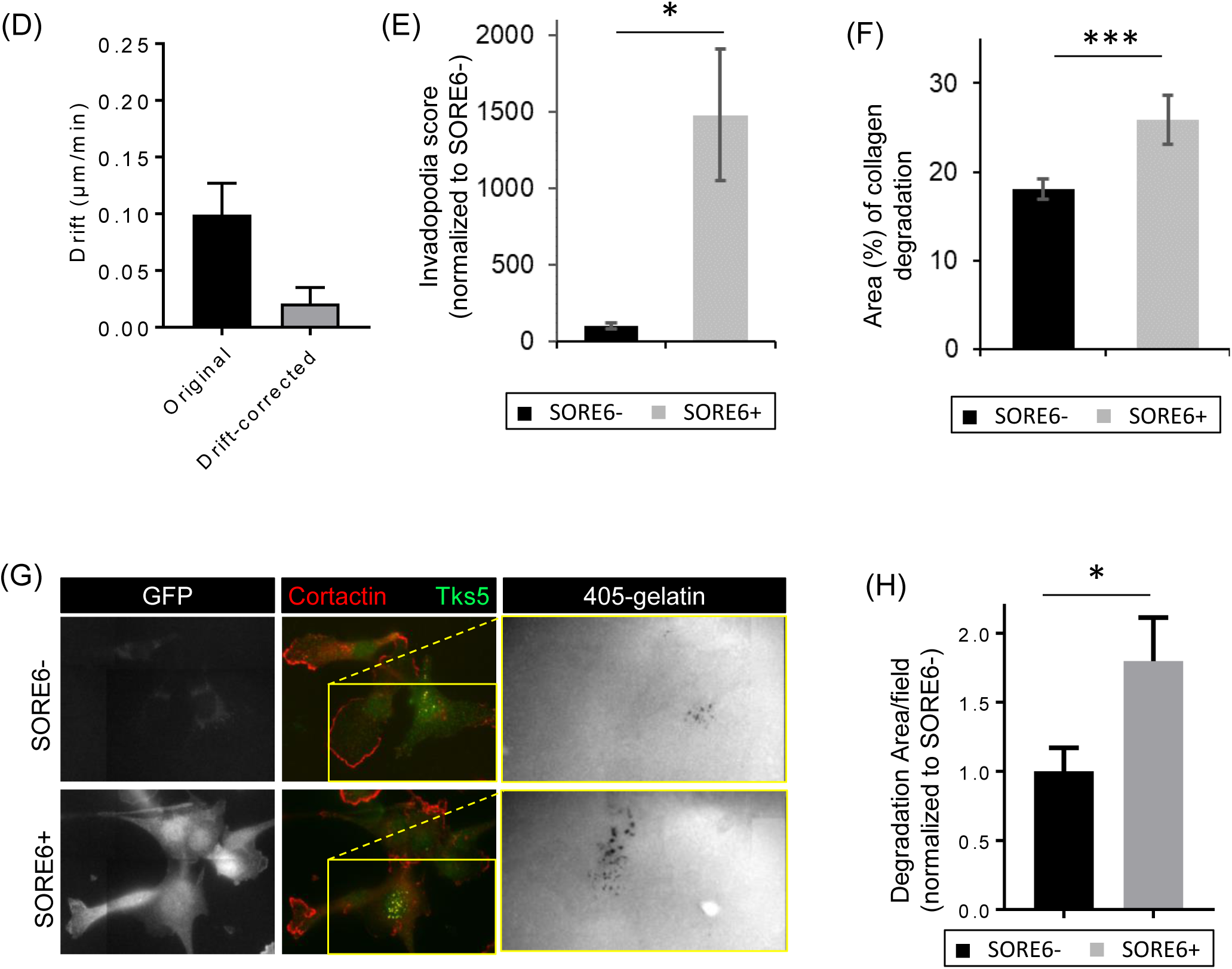
Non-stem cells (SORE6-) do not exhibit spike shaped invadopodium protrusions in vivo. (A) Time-lapse panels from intravital movies 1 and 2 depicting slow and fast motility phenotypes of CSC and non-CSC (related to Fig 2A). (B) Time-lapse panels, from movie 6, for SORE6-fast migrating cells show amoeboid movement without spike shaped invasive protrusions. Bottom panels show the absence of spike shaped protrusions more clearly after applying the edge filter. (C) Time-lapse panels, from movie 7, for SORE6-cell close to blood vessel (BV: white) show amoeboid movement without spike shaped invasive protrusions. Bottom panels show the absence of spike shaped protrusions more clearly after applying the edge filter. (D) Quantification of XY drift in time-lapse intravital movies of live breast tumor tissue before and after correcting for the drift using HyperStackReg plugin (Movie 8, see materials and methods). Data plotted as mean ± SEM, n=5 fields of 340×340 μm^2^ from 3 mice. (E) Invadopodia score quantification (invadopodia identified as Cortactin/Tks5 co-localized invadopodium core) in non-stem and SORE6+ stem cells using a manual 12 μm ROI scoring method shows that SORE6+ stem cells are enriched in invadopodial protrusions. n=5 fields at 40X magnification from 4 tumors; showing mean ± SEM, paired two-tailed Student’s t test, * p-value=0.032. (F) Quantification of collagen degradation area in non-stem and stem cell areas using a manual 12 μm ROI scoring method. n=6 fields at 40X magnification from 4 SORE6 tumors; showing mean ± SEM, paired two-tailed Student’s t test, *** p-value=0.0095. (G) In vitro invadopodia-associated degradation assay was performed with sorted SORE6-and SORE6+ cells by plating them on fluorescent matrix overnight. Invadopodium cores were visualized by staining cells with cortactin and Tks5 antibodies. Yellow rectangles in middle panels indicate the zoomed area shown for the 405-gelatin images. Black areas in the 405-gelatin channel depict degraded matrix. (H) Quantification of matrix degradation by SORE6- and SORE6+ cells show increased degradation potential of SORE6+ stem cells. n=85 fields of 110×110 μm^2^ at 60X magnification for SORE6- and n=61 fields of 110×110 μm^2^ at 60X magnification for SORE6+ samples; plot showing mean ± SEM, unpaired two-tailed Student’s t test, * p-value=0.0187.

**Figure S5:**
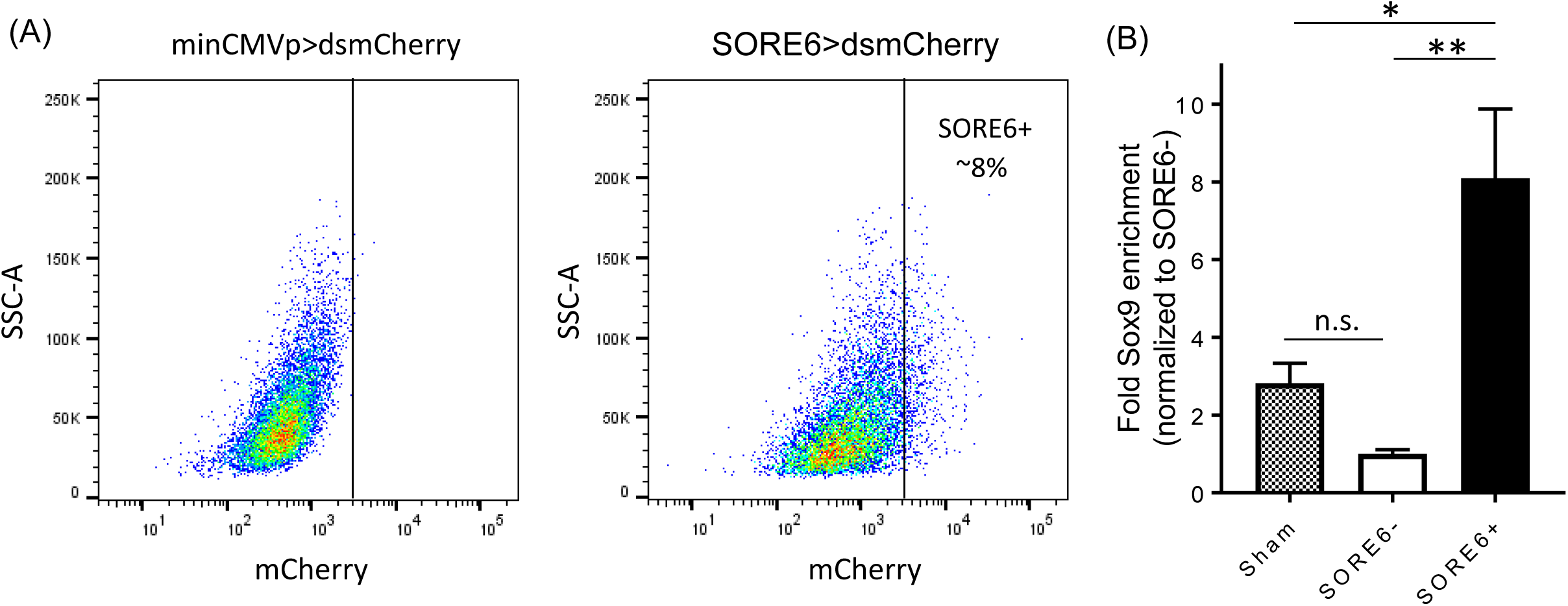
SORE6 reporter marks a minority cell population that is enriched in Sox9 stemness transcription factor in PyMT-derived Met-1 cells. (A) FACS analysis showing that the SORE6 reporter identifies a minority population in cultures of Met-1 cells. SSC-A, side scatter. (B) Quantitative RT-PCR to assess the expression of Sox9, a master stem cell transcription factor, in FACS-sorted SORE6+, SORE6- and sham-sorted Met-1 cells. All values were normalized to the SORE6-mean and results are plotted as mean ± SEM (n=3). Tukey’s multiple comparisons test; p-values: ns = 0.4953, * = 0.0307, ** = 0.0082. Note an 8-fold Sox9 enrichment in SORE6+ cells compared to SORE6-cells.

**Figure S6:**
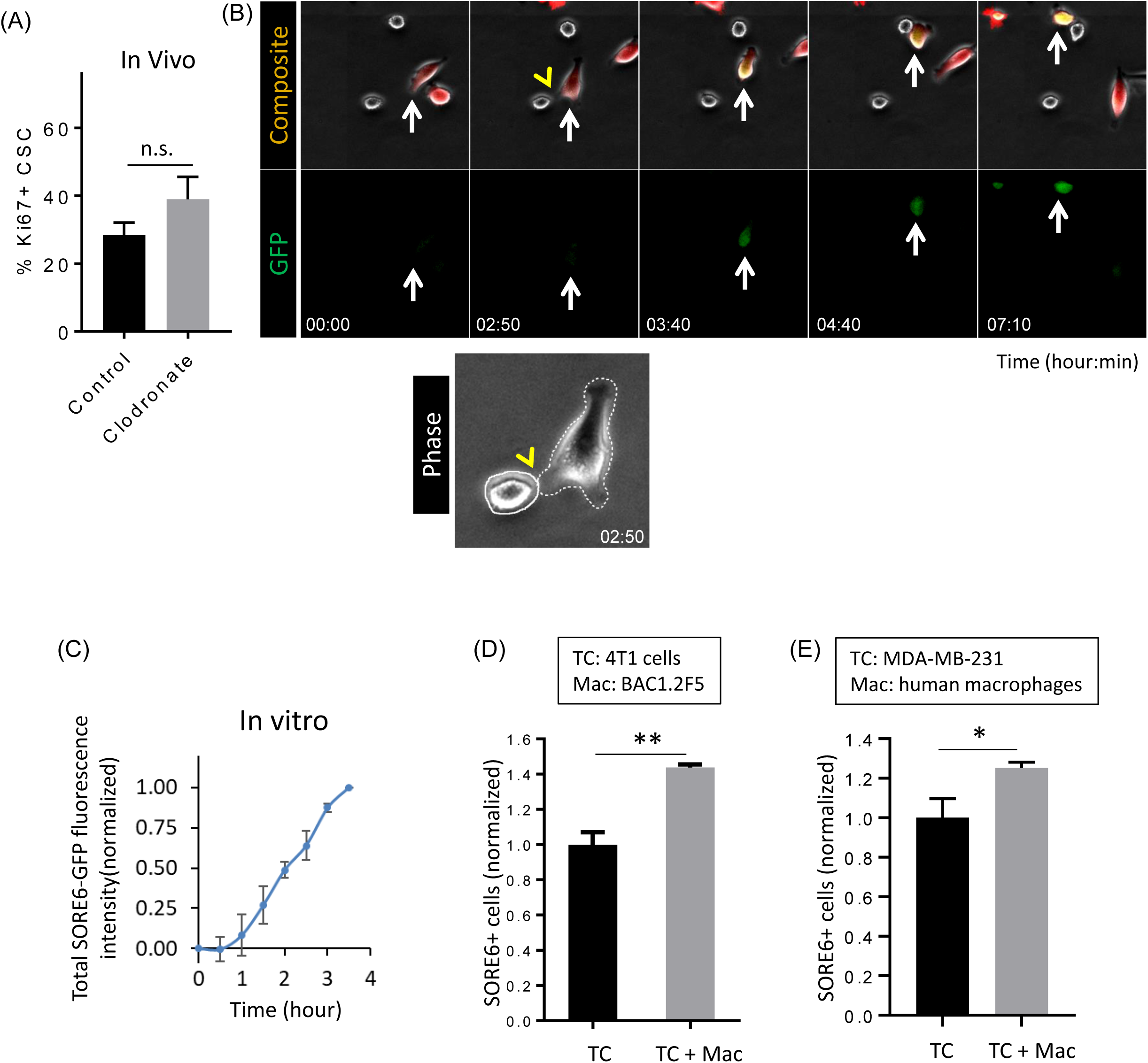
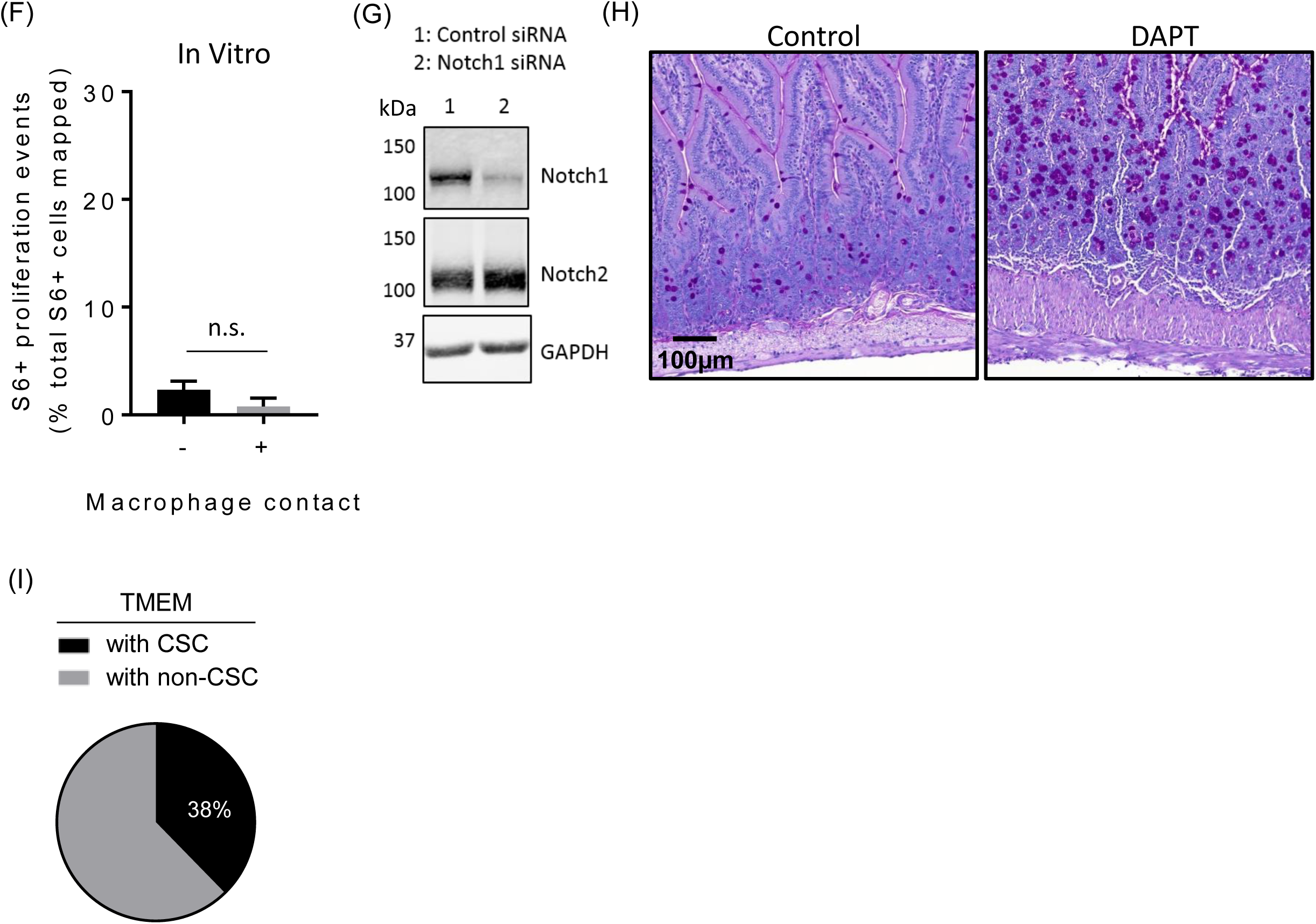
Macrophage contact induces stemness (SORE6 positivity) in multiple breast cancer cell lines. (A) Quantification of % CSCs that are Ki67+ in PyMT primary tumor tissue treated with either PBS liposomes (control) or clodronate liposomes and stained with Sox9 and Ki67 antibodies. n= 11 (control) and 7 (clodronate), 2-3 mm^2^ fields from 5 mice for each case; data plotted as mean ± SEM, unpaired Student’s t test, n.s. p-value=0.148. (B) Macrophage contact induces stemness in tumor cell in *in vitro* co-culture. Representative images from time-lapse movie 11. Time lapse frames were taken every 10 minutes. Composite panel: a non-stem cancer cell (red volume marker td-tomato) transitioning to a CSC (yellow) upon BAC1.2F5 macrophage (gray cell) contact. Yellow arrowhead points to the moment of macrophage contact with the non-stem cancer cell leading to its induction to become a CSC. GFP panel (green channel) shows only SORE6-GFP stem reporter for clarity. White arrows indicate a cancer cell transitioning from non-stem to stem. Zoomed-in phase channel below shows the tumor cell outline in white dotted and macrophage in white solid line. (C) Quantification of total SORE6-GFP fluorescence intensity during stemness induction in tumor cells in vitro in the tumor cell-macrophage co-culture assay. Time 0 corresponds to the time when macrophage contacts the tumor cell. n>100 cells analyzed to identify tumor cells contacting macrophages and showing induced stemness, each data point plotted as mean ± SEM. Data normalized to SORE6-GFP value before macrophage contact to set the 0 background and max SORE6-GFP value as 1 after macrophage contact. (D) In vitro co-culture of 4T1 tumor cells with or without macrophages showing macrophage-dependent induction of stemness in tumor cells. Tumor cells transduced with the SORE6>GFP biosensor were co-cultured with macrophages for 4 days before FACS analysis to determine the fraction of SORE6+ CSCs in the total tumor cell population. TC: 4T1 tumor cells, Mac: BAC1.2F5 macrophages; data plotted as mean ± SEM (n=3), and normalized to the no macrophage condition, unpaired two-tailed Student’s t test, ** p-value=0.0037. (E) In vitro co-culture of MDA-MB-231 tumor cells with or without human macrophages showing macrophage induction of stemness in tumor cells. Tumor cells were co-cultured with macrophages for 2 days before FACS analysis to determine the fraction of SORE6+ CSCs in the total tumor cell population. TC: MDA-MB-231-LM2 tumor cells, Mac: human peripheral blood monocyte-derived macrophages polarized to an M2-phenotype with CSF1 and IL-4; data plotted as mean ± SEM (n=3), and normalized to the no macrophage condition, unpaired two-tailed Student’s t test, * p-value=0.012. (F) Quantification of SORE6+ (S6+) cell proliferation due to division of pre-existing S6+ cells without or with macrophage contact in TC alone (black bar) or TC+Mac (gray bar) co-culture assay respectively, using origin mapping of S6+ cells in time-lapse movies. Data plotted as mean ± SEM, unpaired two-tailed Student’s t test, n.s. p-value=0.207, n=4 wells/condition for a total of 128 cells traced/condition. (G) Western blot showing Notch1 KD in Notch1 siRNA treated SORE6 MDA-MB-231 cells after 48 h. Note that Notch2 levels are unaffected by Notch1 siRNA. (H) Periodic acid-Schiff (PAS) staining of the duodenum from MMTV-PyMT mice after 14 days of treatment with either vehicle control (left panel) or DAPT (right panel). The data confirm that in vivo DAPT-mediated Notch signaling suppression induces goblet cell (dark purple dots) differentiation. (I) Frequency analysis of CSC (SORE6+) and non-CSC (SORE6-) containing TMEMs in SORE6>GFP xenograft MDA-MB-231 primary tumors. n=4 fields of 2.2×2.2 mm^2^ from 4 mice.

## Supplementary Movies

**Movie 1:** Intravital microscopy movie showing slow locomotion of SORE6+ stem cells (yellow). Blue (SHG): collagen fibers Time in hour:min:sec.

**Movie 2:** Intravital microscopy movie showing fast locomotion of SORE6-non-stem cells (red). Time in hour:min:sec.

**Movie 3:** Intravital microscopy movie showing thin invadopodial protrusion (white arrows) from CSC (yellow) interacting with ECM fiber (blue, SHG). Time in min:sec.

**Movie 4:** Left: Intravital microscopy movie of live breast carcinoma stem cell (yellow) showing invadopodial protrusions directed towards blood vessel (gray). Blood channel has been averaged to eliminate fluctuations in intensity due to passing erythrocytes and contrast agent clearance. Right: Processed movie with the edge filter clearly shows dynamic protrusions directed towards the blood vessel. Time in hour:min:sec

**Movie 5:** 3D reconstruction of CSC (yellow) and blood vessel (white) from Movie 4 with rotated views show invadopodium protrusion directed toward the blood vessel.

**Movie 6:** Non-stem cells (SORE6-) away from the blood vessel showing no cellular protrusion. (A, B) Red: tdTomato expressing and SORE6-non-stem cells; white: thresholded movie (C-E) To investigate cellular protrusions in non-stem cells, the overall translation of the cell was subtracted from each frame, so the cells appear stationary. The movie shows absence of fine cellular protrusions in non-stem cells. (E) Movie processed with edge filter.

**Movie 7:** Left: Non-stem cells (SORE6-) close to blood vessel. Right: Movie processed with edge filter show the absences of cellular protrusion in a non-stem cell close to blood vessel.

**Movie 8:** A montage of original (left) and drift-corrected (right) intravital movies of live breast tumor tissue showing stem cells (green), non-stem cells (red) and collagen fibers (blue). Time in hour:min:sec. Tissue drift after post-acquisition drift correction is minimal (∼ 0.02 μm/min, Fig S4D) and does not contribute much to the cell migration on the time scale of the movies.

**Movie 9:** Intravital movie of macrophage (cyan) touching a non-stem cell. After contact, stemness is induced in the in cancer cell *in vivo* as determined by the increase in GFP signal (green) over time. White dotted lines show tumor cell outlines before and after stemness induction and is drawn based on the tdTomato volume marker channel (shown on the right). Collagen fibers are shown in blue (SHG). Time in hour:min:sec.

**Movie 10:** Intravital movie 9 shown with multiple Z-planes (each 5 μm apart), showing the induction of stemness in a non-stem cell. Note that the cell changes its color from red (tdTomato) to yellow (tdTomato+GFP). White dotted lines show tumor cell outlines in different Z-planes before and after stemness induction and is drawn based on the tdTomato volume marker channel (top panels). Collagen fibers are shown in blue (SHG). Time in hour:min:sec.

**Movie 11:** Time-lapsed movie showing macrophage-induced stemness in a non-stem tumor cell in the *in vitro* tumor cell-macrophage co-culture assay. Macrophage (unlabeled in gray in the rightmost panel) can be seen to touch a non-stem tumor cell. After contact, stemness is induced in the cancer cell as determined by the increase in GFP signal (green) over time (middle panel). Time in hour:min

**Movie 12:** Intravital movie of SORE6+ tumor cells (yellow) showing CSCs enriched in perivascular regions and in contact with perivascular macrophages (cyan) in live tissue. Blood vessels are labeled with far-red quantum dots and shown in gray. Fluctuations in blood vessel intensity are due to passing erythrocytes and contrast agent clearance. Blood vessel channel was used to outline vessels with dotted lines in all the frames. Time in hour:min

**Movie 13:** *In vivo* intravital movie of a single SORE6+ stem cell (green) intravasating into the blood vessel in primary tumor. Macrophages are shown in cyan. Blood vessels were labeled with far-red quantum dots (shown in gray), and outlined in white dotted line.

## Notes

### Competing Interest Statement

The authors have declared no competing interest.

## REFERENCES

1. Bouvard C, Barefield C, Zhu S. (2014). Cancer stem cells as a target population for drug discovery. Future Med Chem. 6(14):1567–85. PMID: 25367391.

2. Valastyan S, Weinberg RA. (2011). Tumor metastasis: molecular insights and evolving paradigms. Cell. 147(2):275–92. PMID: 22000009 / PMCID: PMC3261217.

3. Al-Hajj M, Wicha MS, Benito-Hernandez A, Morrison SJ, Clarke MF. (2003). Prospective identification of tumorigenic breast cancer cells. Proc Natl Acad Sci U S A. 100(7):3983–8. PMID: 12629218 / PMCID: PMC153034.

4. Nassar D, Blanpain C. (2016). Cancer Stem Cells: Basic Concepts and Therapeutic Implications. Annual review of pathology. 11:47–76. PMID: 27193450.

5. Zhao J. (2016). Cancer stem cells and chemoresistance: The smartest survives the raid. Pharmacol Ther. 160:145–58. PMID: 26899500 / PMCID: PMC4808328.

6. Quail DF, Joyce JA. (2013). Microenvironmental regulation of tumor progression and metastasis. Nat Med. 19(11):1423–37. PMID: 24202395 / PMCID: PMC3954707.

7. Roussos ET, Condeelis JS, Patsialou A. (2011). Chemotaxis in cancer. Nature reviews Cancer. 11(8):573–87. PMID: 21779009 / PMCID: PMC4030706.

8. Plaks V, Kong N, Werb Z. (2015). The cancer stem cell niche: how essential is the niche in regulating stemness of tumor cells? Cell stem cell. 16(3):225–38. PMID: 25748930 / PMCID: PMC4355577.

9. Batlle E, Clevers H. (2017). Cancer stem cells revisited. Nat Med. 23(10):1124–34. PMID: 28985214.

10. Chaffer CL, Marjanovic ND, Lee T, Bell G, Kleer CG, Reinhardt F, D’Alessio AC, Young RA, Weinberg RA. (2013). Poised chromatin at the ZEB1 promoter enables breast cancer cell plasticity and enhances tumorigenicity. Cell. 154(1):61–74. PMID: 23827675 / PMCID: PMC4015106.

11. Lilja AM, Rodilla V, Huyghe M, Hannezo E, Landragin C, Renaud O, Leroy O, Rulands S, Simons BD, Fre S. (2018). Clonal analysis of Notch1-expressing cells reveals the existence of unipotent stem cells that retain long-term plasticity in the embryonic mammary gland. Nature cell biology. 20(6):677–87. PMID: 29784917.

12. Wuidart A, Sifrim A, Fioramonti M, Matsumura S, Brisebarre A, Brown D, Centonze A, Dannau A, Dubois C, Van Keymeulen A, Voet T, Blanpain C. (2018). Publisher Correction: Early lineage segregation of multipotent embryonic mammary gland progenitors. <u>Nature cell biology</u>. PMID: 30018320.

13. Tang B, Raviv A, Esposito D, Flanders KC, Daniel C, Nghiem BT, Garfield S, Lim L, Mannan P, Robles AI, Smith WI, Jr., Zimmerberg J, Ravin R, Wakefield LM. (2015). A flexible reporter system for direct observation and isolation of cancer stem cells. Stem cell reports. 4(1):155–69. PMID: 25497455 / PMCID: PMC4297872.

14. Bierie B, Pierce SE, Kroeger C, Stover DG, Pattabiraman DR, Thiru P, Liu Donaher J, Reinhardt F, Chaffer CL, Keckesova Z, Weinberg RA. (2017). Integrin-beta4 identifies cancer stem cell-enriched populations of partially mesenchymal carcinoma cells. Proc Natl Acad Sci U S A. 114(12):E2337-E46. PMID: 28270621 / PMCID: PMCPMC5373369.

15. Kalluri R, Weinberg RA. (2009). The basics of epithelial-mesenchymal transition. The Journal of clinical investigation. 119(6):1420–8. PMID: 19487818 / PMCID: PMC2689101.

16. Nieto MA, Huang RY, Jackson RA, Thiery JP. (2016). Emt: 2016. Cell. 166(1):21–45. PMID: 27368099.

17. Thiery JP. (2002). Epithelial-mesenchymal transitions in tumour progression. Nature reviews Cancer. 2(6):442–54. PMID: 12189386.

18. Patsialou A, Bravo-Cordero JJ, Wang Y, Entenberg D, Liu H, Clarke M, Condeelis JS. (2013). Intravital multiphoton imaging reveals multicellular streaming as a crucial component of in vivo cell migration in human breast tumors. Intravital. 2(2):e25294. PMID: 25013744 / PMCID: PMC3908591.

19. Gligorijevic B, Bergman A, Condeelis J. (2014). Multiparametric classification links tumor microenvironments with tumor cell phenotype. PLoS Biol. 12(11):e1001995. PMID: 25386698 / PMCID: PMC4227649

20. Harney AS, Arwert EN, Entenberg D, Wang Y, Guo P, Qian BZ, Oktay MH, Pollard JW, Jones JG, Condeelis JS. (2015). Real-Time Imaging Reveals Local, Transient Vascular Permeability, and Tumor Cell Intravasation Stimulated by TIE2hi Macrophage-Derived VEGFA. Cancer Discovery. 5(9):932–43. PMID: 26269515 / PMCID: PMC4560669.

21. Sharma VP, Eddy R, Entenberg D, Kai M, Gertler FB, Condeelis J. (2013). Tks5 and SHIP2 regulate invadopodium maturation, but not initiation, in breast carcinoma cells. Curr Biol. 23(21):2079–89. PMID: 24206842 / PMCID: PMC3882144.

22. Gligorijevic B, Wyckoff J, Yamaguchi H, Wang Y, Roussos ET, Condeelis J. (2012). N-WASP-mediated invadopodium formation is involved in intravasation and lung metastasis of mammary tumors. Journal of Cell Science. 125(Pt 3):724–34. PMID: 22389406 / PMCID: PMC3367832.

23. Bravo-Cordero JJ, Hodgson L, Condeelis J. (2012). Directed cell invasion and migration during metastasis. Curr Opin Cell Biol. 24(2):277–83. PMID: 22209238 / PMCID: PMC3320684.

24. Magalhaes MA, Larson DR, Mader CC, Bravo-Cordero JJ, Gil-Henn H, Oser M, Chen X, Koleske AJ, Condeelis J. (2011). Cortactin phosphorylation regulates cell invasion through a pH-dependent pathway. J Cell Biol. 195(5):903–20. PMID: 22105349 / PMCID: PMC3257566.

25. Beaty BT, Wang Y, Bravo-Cordero JJ, Sharma VP, Miskolci V, Hodgson L, Condeelis J. (2014). Talin regulates moesin-NHE-1 recruitment to invadopodia and promotes mammary tumor metastasis. J Cell Biol. 205(5):737–51. PMID: 24891603 / PMCID: PMC4050723.

26. Eddy RJ, Weidmann MD, Sharma VP, Condeelis JS. (2017). Tumor Cell Invadopodia: Invasive Protrusions that Orchestrate Metastasis. Trends Cell Biology. 27(8):595–607. PMID: 28412099 / PMCID: PMC5524604.

27. Eckert MA, Lwin TM, Chang AT, Kim J, Danis E, Ohno-Machado L, Yang J. (2011). Twist1-induced invadopodia formation promotes tumor metastasis. Cancer cell. 19(3):372–86. PMID: 21397860 / PMCID: PMC3072410.

28. Lu H, Clauser KR, Tam WL, Frose J, Ye X, Eaton EN, Reinhardt F, Donnenberg VS, Bhargava R, Carr SA, Weinberg RA. (2014). A breast cancer stem cell niche supported by juxtacrine signalling from monocytes and macrophages. Nature cell biology. 16(11):1105–17. PMID: 25266422 / PMCID: PMC4296514.

29. Karagiannis GS, Pastoriza JM, Wang Y, Harney AS, Entenberg D, Pignatelli J, Sharma VP, Xue EA, Cheng E, D’Alfonso TM, Jones JG, Anampa J, Rohan TE, Sparano JA, Condeelis JS, Oktay MH. (2017). Neoadjuvant chemotherapy induces breast cancer metastasis through a TMEM-mediated mechanism. Science Translational Medicine. 9(397). PMID: 28679654 / PMCID: PMC5592784.

30. Sanchez LR, Borriello L, Entenberg D, Condeelis JS, Oktay MH, Karagiannis GS. (2019). The emerging roles of macrophages in cancer metastasis and response to chemotherapy. J Leukoc Biol. 106(2):259–74. PMID: 30720887 / PMCID: PMCPMC6779158.

31. Pignatelli J, Goswami S, Jones JG, Rohan TE, Pieri E, Chen X, Adler E, Cox D, Maleki S, Bresnick A, Gertler FB, Condeelis JS, Oktay MH. (2014). Invasive breast carcinoma cells from patients exhibit MenaINV- and macrophage-dependent transendothelial migration. Science signaling. 7(353):ra112. PMID: 25429076 / PMCID: PMC4266931.

32. Pignatelli J, Bravo-Cordero JJ, Roh-Johnson M, Gandhi SJ, Wang Y, Chen X, Eddy RJ, Xue A, Singer RH, Hodgson L, Oktay MH, Condeelis JS. (2016). Macrophage-dependent tumor cell transendothelial migration is mediated by Notch1/MenaINV-initiated invadopodium formation. Sci Rep. 6:37874. PMID: 27901093 / PMCID: PMC5129016

33. Entenberg D, Voiculescu S, Guo P, Borriello L, Wang Y, Karagiannis GS, Jones J, Baccay F, Oktay M, Condeelis J. (2018). A permanent window for the murine lung enables high-resolution imaging of cancer metastasis. Nat Methods. 15(1):73–80. PMID: 29176592 / PMCID: PMCPMC5755704.

34. Guy CT CR, Muller WJ. (1992). Induction of mammary tumors by expression of polyomavirus middle T oncogene: a transgenic mouse model for metastatic disease. Mol Cell Biol. 12(3). / PMCID: PMCPMC369527

35. Jessen KA, Liu SY, Tepper CG, Karrim J, McGoldrick ET, Rosner A, Munn RJ, Young LJ, Borowsky AD, Cardiff RD, Gregg JP. (2004). Molecular analysis of metastasis in a polyomavirus middle T mouse model: the role of osteopontin. Breast cancer research : BCR. 6(3):R157–69. PMID: 15084239 / PMCID: PMC400667.

36. Guo W, Keckesova Z, Donaher JL, Shibue T, Tischler V, Reinhardt F, Itzkovitz S, Noske A, Zurrer-Hardi U, Bell G, Tam WL, Mani SA, van Oudenaarden A, Weinberg RA. (2012). Slug and Sox9 cooperatively determine the mammary stem cell state. Cell. 148(5):1015–28. PMID: 22385965 / PMCID: PMC3305806.

37. Christin JR, Wang, C., Chung, C.Y., Liu, Y., Dravis, C., Tang, W, Oktay, M.H., Wahl, G. M., Guo W. (2020). Stem cell determinant SOX9 promotes lineage plasticity and progression in basal-like breast cancer. <u>Cell Reports</u>.

38. Richtig G, Aigelsreiter A, Schwarzenbacher D, Ress AL, Adiprasito JB, Stiegelbauer V, Hoefler G, Schauer S, Kiesslich T, Kornprat P, Winder T, Eisner F, Gerger A, Stoeger H, Stauber R, Lackner C, Pichler M. (2017). SOX9 is a proliferation and stem cell factor in hepatocellular carcinoma and possess widespread prognostic significance in different cancer types. PLoS One. 12(11):e0187814. PMID: 29121666 / PMCID: PMC5679634.

39. Chakrabarti R, Celia-Terrassa T, Kumar S, Hang X, Wei Y, Choudhury A, Hwang J, Peng J, Nixon B, Grady JJ, DeCoste C, Gao J, van Es JH, Li MO, Aifantis I, Clevers H, Kang Y. (2018). Notch ligand Dll1 mediates cross-talk between mammary stem cells and the macrophageal niche. Science. 360(6396). PMID: 29773667.

40. Bocci F, Gearhart-Serna L, Boareto M, Ribeiro M, Ben-Jacob E, Devi GR, Levine H, Onuchic JN, Jolly MK. (2019). Toward understanding cancer stem cell heterogeneity in the tumor microenvironment. Proc Natl Acad Sci U S A. 116(1):148–57. PMID: 30587589 / PMCID: PMCPMC6320545.

41. Yamaguchi E, Chiba S, Kumano K, Kunisato A, Takahashi T, Takahashi T, Hirai H. (2002). Expression of Notch ligands, Jagged1, 2 and Delta1 in antigen presenting cells in mice. Immunol Lett. 81(1):59–64. PMID: 11841846.

42. Cabrera RM, Mao SPH, Surve CR, Condeelis JS, Segall JE. (2018). A novel neuregulin - jagged1 paracrine loop in breast cancer transendothelial migration. Breast cancer research : BCR. 20(1):24. PMID: 29636067 / PMCID: PMC5894135.

43. Purow B. (2012). Notch inhibition as a promising new approach to cancer therapy. Advances in experimental medicine and biology. 727:305–19. PMID: 22399357 / PMCID: PMC3361718.

44. Andersson ER, Lendahl U. (2014). Therapeutic modulation of Notch signalling--are we there yet? Nature reviews Drug discovery. 13(5):357–78. PMID: 24781550.

45. Harney AS, Karagiannis GS, Pignatelli J, Smith BD, Kadioglu E, Wise SC, Hood MM, Kaufman MD, Leary CB, Lu WP, Al-Ani G, Chen X, Entenberg D, Oktay MH, Wang Y, Chun L, De Palma M, Jones JG, Flynn DL, Condeelis JS. (2017). The Selective Tie2 Inhibitor Rebastinib Blocks Recruitment and Function of Tie2(Hi) Macrophages in Breast Cancer and Pancreatic Neuroendocrine Tumors. Molecular Cancer Therapeutics. 16(11):2486–501. PMID: 28838996 / PMCID: PMC5669998.

46. Rohan TE, Xue X, Lin HM, D’Alfonso TM, Ginter PS, Oktay MH, Robinson BD, Ginsberg M, Gertler FB, Glass AG, Sparano JA, Condeelis JS, Jones JG. (2014). Tumor microenvironment of metastasis and risk of distant metastasis of breast cancer. Journal of the National Cancer Institute. 106(8). PMID: 24895374 / PMCID: PMC4133559.

47. Robinson BD, Sica GL, Liu YF, Rohan TE, Gertler FB, Condeelis JS, Jones JG. (2009). Tumor microenvironment of metastasis in human breast carcinoma: a potential prognostic marker linked to hematogenous dissemination. Clinical Cancer Research 15(7):2433–41. PMID: 19318480 / PMCID: PMC3156570.

48. Sparano JA, Gray R, Oktay MH, Entenberg D, Rohan T, Xue X, Donovan M, Peterson M, Shuber A, Hamilton DA, D’Alfonso T, Goldstein LJ, Gertler F, Davidson NE, Condeelis J, Jones J. (2017). A metastasis biomarker (MetaSite Breast Score) is associated with distant recurrence in hormone receptor-positive, HER2-negative early-stage breast cancer. Nature PJ Breast Cancer. 3:42. PMID: 29138761 / PMCID: PMC5678158.

49. Arwert EN, Harney AS, Entenberg D, Wang Y, Sahai E, Pollard JW, Condeelis JS. (2018). A Unidirectional Transition from Migratory to Perivascular Macrophage Is Required for Tumor Cell Intravasation. Cell Rep. 23(5):1239–48. PMID: 29719241 / PMCID: PMC5946803.

50. Malanchi I, Santamaria-Martinez A, Susanto E, Peng H, Lehr HA, Delaloye JF, Huelsken J. (2011). Interactions between cancer stem cells and their niche govern metastatic colonization. Nature. 481(7379):85–9. PMID: 22158103.

51. Brooks MD, Burness ML, Wicha MS. (2015). Therapeutic Implications of Cellular Heterogeneity and Plasticity in Breast Cancer. Cell stem cell. 17(3):260–71. PMID: 26340526 / PMCID: PMCPMC4560840.

52. Liu S, Cong Y, Wang D, Sun Y, Deng L, Liu Y, Martin-Trevino R, Shang L, McDermott SP, Landis MD, Hong S, Adams A, D’Angelo R, Ginestier C, Charafe-Jauffret E, Clouthier SG, Birnbaum D, Wong ST, Zhan M, Chang JC, Wicha MS. (2014). Breast cancer stem cells transition between epithelial and mesenchymal states reflective of their normal counterparts. Stem cell reports. 2(1):78–91. PMID: 24511467 / PMCID: PMC3916760.

53. Grosse-Gehling P, Fargeas CA, Dittfeld C, Garbe Y, Alison MR, Corbeil D, Kunz-Schughart LA. (2013). CD133 as a biomarker for putative cancer stem cells in solid tumours: limitations, problems and challenges. J Pathol. 229(3):355–78. PMID: 22899341.

54. Kim WT, Ryu CJ. (2017). Cancer stem cell surface markers on normal stem cells. BMB Rep. 50(6):285–98. PMID: 28270302 / PMCID: PMCPMC5498139.

55. Karagiannis GS, Goswami S, Jones JG, Oktay MH, Condeelis JS. (2016). Signatures of breast cancer metastasis at a glance. Journal of Cell Science. 129(9):1751–8. PMID: 27084578 / PMCID: PMC4893654.

56. Shaw FL, Harrison H, Spence K, Ablett MP, Simoes BM, Farnie G, Clarke RB. (2012). A detailed mammosphere assay protocol for the quantification of breast stem cell activity. J Mammary Gland Biol Neoplasia. 17(2):111–7. PMID: 22665270.

57. Lawson DA, Bhakta NR, Kessenbrock K, Prummel KD, Yu Y, Takai K, Zhou A, Eyob H, Balakrishnan S, Wang CY, Yaswen P, Goga A, Werb Z. (2015). Single-cell analysis reveals a stem-cell program in human metastatic breast cancer cells. Nature. 526(7571):131–5. PMID: 26416748 / PMCID: PMC4648562.

58. Liu X, Taftaf R, Kawaguchi M, Chang YF, Chen W, Entenberg D, Zhang Y, Gerratana L, Huang S, Patel DB, Tsui E, Adorno-Cruz V, Chirieleison SM, Cao Y, Harney AS, Patel S, Patsialou A, Shen Y, Avril S, Gilmore HL, Lathia JD, Abbott DW, Cristofanilli M, Condeelis JS, Liu H. (2018). Homophilic CD44 interactions mediate tumor cell aggregation and polyclonal metastasis in patient-derived breast cancer models. Cancer Discov. PMID: 30361447.

59. Aceto N, Bardia A, Miyamoto DT, Donaldson MC, Wittner BS, Spencer JA, Yu M, Pely A, Engstrom A, Zhu H, Brannigan BW, Kapur R, Stott SL, Shioda T, Ramaswamy S, Ting DT, Lin CP, Toner M, Haber DA, Maheswaran S. (2014). Circulating tumor cell clusters are oligoclonal precursors of breast cancer metastasis. Cell. 158(5):1110–22. PMID: 25171411 / PMCID: PMC4149753.

60. Gkountela S, Castro-Giner F, Szczerba BM, Vetter M, Landin J, Scherrer R, Krol I, Scheidmann MC, Beisel C, Stirnimann CU, Kurzeder C, Heinzelmann-Schwarz V, Rochlitz C, Weber WP, Aceto N. (2019). Circulating Tumor Cell Clustering Shapes DNA Methylation to Enable Metastasis Seeding. Cell. 176(1-2):98–112 e14. PMID: 30633912 / PMCID: PMC6363966.

61. Liu D, Lu Q, Wang X, Wang J, Lu N, Jiang Z, Hao X, Li J, Liu J, Cao P, Peng G, Tao Y, Zhao D, He F, Tang L. (2019). LSECtin on tumor-associated macrophages enhances breast cancer stemness via interaction with its receptor BTN3A3. Cell research. PMID: 30858559.

62. Wang J, Sullenger BA, Rich JN. (2012). Notch signaling in cancer stem cells. Advances in experimental medicine and biology. 727:174–85. PMID: 22399347.

63. Reedijk M, Odorcic S, Chang L, Zhang H, Miller N, McCready DR, Lockwood G, Egan SE. (2005). High-level coexpression of JAG1 and NOTCH1 is observed in human breast cancer and is associated with poor overall survival. Cancer Res. 65(18):8530–7. PMID: 16166334.

64. Leung E, Xue A, Wang Y, Rougerie P, Sharma VP, Eddy R, Cox D, Condeelis J. (2017). Blood vessel endothelium-directed tumor cell streaming in breast tumors requires the HGF/C-Met signaling pathway. Oncogene. 36(19):2680–92. PMID: 27893712 / PMCID: PMCPMC5426963.

65. Weidmann MD, Surve CR, Eddy RJ, Chen X, Gertler FB, Sharma VP, Condeelis JS. (2016). MenaINV dysregulates cortactin phosphorylation to promote invadopodium maturation. Sci Rep. 6:36142. PMID: 27824079 / PMCID: PMC5099927.

66. Roussos ET, Balsamo M, Alford SK, Wyckoff JB, Gligorijevic B, Wang Y, Pozzuto M, Stobezki R, Goswami S, Segall JE, Lauffenburger DA, Bresnick AR, Gertler FB, Condeelis JS. (2011). Mena invasive (MenaINV) promotes multicellular streaming motility and transendothelial migration in a mouse model of breast cancer. Journal of Cell Science. 124(Pt 13):2120–31. PMID: 21670198 / PMCID: PMC3113666.

67. Aktas B, Tewes M, Fehm T, Hauch S, Kimmig R, Kasimir-Bauer S. (2009). Stem cell and epithelial-mesenchymal transition markers are frequently overexpressed in circulating tumor cells of metastatic breast cancer patients. Breast cancer research : BCR. 11(4):R46. PMID: 19589136 / PMCID: PMC2750105.

68. Janni WJ, Rack B, Terstappen LW, Pierga JY, Taran FA, Fehm T, Hall C, de Groot MR, Bidard FC, Friedl TW, Fasching PA, Brucker SY, Pantel K, Lucci A. (2016). Pooled Analysis of the Prognostic Relevance of Circulating Tumor Cells in Primary Breast Cancer. Clin Cancer Res. 22(10):2583–93. PMID: 26733614.

69. Riethdorf S, Muller V, Loibl S, Nekljudova V, Weber K, Huober J, Fehm T, Schrader I, Hilfrich J, Holms F, Tesch H, Schem C, von Minckwitz G, Untch M, Pantel K. (2017). Prognostic Impact of Circulating Tumor Cells for Breast Cancer Patients Treated in the Neoadjuvant “Geparquattro” Trial. Clin Cancer Res. 23(18):5384–93. PMID: 28679772.

70. Bidard FC, Michiels S, Riethdorf S, Mueller V, Esserman LJ, Lucci A, Naume B, Horiguchi J, Gisbert-Criado R, Sleijfer S, Toi M, Garcia-Saenz JA, Hartkopf A, Generali D, Rothe F, Smerage J, Muinelo-Romay L, Stebbing J, Viens P, Magbanua MJM, Hall CS, Engebraaten O, Takata D, Vidal-Martinez J, Onstenk W, Fujisawa N, Diaz-Rubio E, Taran FA, Cappelletti MR, Ignatiadis M, Proudhon C, Wolf DM, Bauldry JB, Borgen E, Nagaoka R, Caranana V, Kraan J, Maestro M, Brucker SY, Weber K, Reyal F, Amara D, Karhade MG, Mathiesen RR, Tokiniwa H, Llombart-Cussac A, Meddis A, Blanche P, d’Hollander K, Cottu P, Park JW, Loibl S, Latouche A, Pierga JY, Pantel K. (2018). Circulating Tumor Cells in Breast Cancer Patients Treated by Neoadjuvant Chemotherapy: A Meta-analysis. Journal of the National Cancer Institute. 110(6):560–7. PMID: 29659933.

71. Baccelli I, Schneeweiss A, Riethdorf S, Stenzinger A, Schillert A, Vogel V, Klein C, Saini M, Bauerle T, Wallwiener M, Holland-Letz T, Hofner T, Sprick M, Scharpff M, Marme F, Sinn HP, Pantel K, Weichert W, Trumpp A. (2013). Identification of a population of blood circulating tumor cells from breast cancer patients that initiates metastasis in a xenograft assay. Nature biotechnology. 31(6):539–44. PMID: 23609047.

72. Zhang L, Ridgway LD, Wetzel MD, Ngo J, Yin W, Kumar D, Goodman JC, Groves MD, Marchetti D. (2013). The identification and characterization of breast cancer CTCs competent for brain metastasis. Sci Transl Med. 5(180):180ra48. PMID: 23576814 / PMCID: PMCPMC3863909.

73. Kreso A, Dick JE. (2014). Evolution of the cancer stem cell model. Cell stem cell. 14(3):275–91. PMID: 24607403.

74. Oskarsson T, Batlle E, Massague J. (2014). Metastatic stem cells: sources, niches, and vital pathways. Cell stem cell. 14(3):306–21. PMID: 24607405 / PMCID: PMC3998185.

75. Papadaki MA, Stoupis G, Theodoropoulos PA, Mavroudis D, Georgoulias V, Agelaki S. (2019). Circulating Tumor Cells with Stemness and Epithelial-to-Mesenchymal Transition Features Are Chemoresistant and Predictive of Poor Outcome in Metastatic Breast Cancer. Molecular Cancer Therapeutics. 18(2):437–47. PMID: 30401696.

76. Lambert AW, Pattabiraman DR, Weinberg RA. (2017). Emerging Biological Principles of Metastasis. Cell. 168(4):670–91. PMID: 28187288 / PMCID: PMCPMC5308465.

77. Shackleton M, Quintana E, Fearon ER, Morrison SJ. (2009). Heterogeneity in cancer: cancer stem cells versus clonal evolution. Cell. 138(5):822–9. PMID: 19737509.

78. Hughes R, Qian BZ, Rowan C, Muthana M, Keklikoglou I, Olson OC, Tazzyman S, Danson S, Addison C, Clemons M, Gonzalez-Angulo AM, Joyce JA, De Palma M, Pollard JW, Lewis CE. (2015). Perivascular M2 Macrophages Stimulate Tumor Relapse after Chemotherapy. Cancer Res. 75(17):3479–91. PMID: 26269531.

79. Chen L, Li J, Wang F, Dai C, Wu F, Liu X, Li T, Glauben R, Zhang Y, Nie G, He Y, Qin Z. (2016). Tie2 Expression on Macrophages Is Required for Blood Vessel Reconstruction and Tumor Relapse after Chemotherapy. Cancer Res. 76(23):6828–38. PMID: 27758887.

80. Chang YS, Jalgaonkar SP, Middleton JD, Hai T. (2017). Stress-inducible gene Atf3 in the noncancer host cells contributes to chemotherapy-exacerbated breast cancer metastasis. Proc Natl Acad Sci U S A. 114(34):E7159–E68. PMID: 28784776 / PMCID: PMCPMC5576783.

81. Karagiannis GS, Condeelis JS, Oktay MH. (2018). Chemotherapy-induced metastasis: mechanisms and translational opportunities. Clin Exp Metastasis. 35(4):269–84. PMID: 29307118 / PMCID: PMCPMC6035114.

82. Creighton CJ, Li X, Landis M, Dixon JM, Neumeister VM, Sjolund A, Rimm DL, Wong H, Rodriguez A, Herschkowitz JI, Fan C, Zhang X, He X, Pavlick A, Gutierrez MC, Renshaw L, Larionov AA, Faratian D, Hilsenbeck SG, Perou CM, Lewis MT, Rosen JM, Chang JC. (2009). Residual breast cancers after conventional therapy display mesenchymal as well as tumor-initiating features. Proc Natl Acad Sci U S A. 106(33):13820–5. PMID: 19666588 / PMCID: PMC2720409.

83. Ginter PS, Karagiannis GS, Entenberg D, Lin Y, Condeelis J, Jones JG, Oktay MH. (2019). Tumor Microenvironment of Metastasis (TMEM) Doorways Are Restricted to the Blood Vessel Endothelium in Both Primary Breast Cancers and Their Lymph Node Metastases. Cancers (Basel). 11(10). PMID: 31597373.

84. Bidard FC, Peeters DJ, Fehm T, Nole F, Gisbert-Criado R, Mavroudis D, Grisanti S, Generali D, Garcia-Saenz JA, Stebbing J, Caldas C, Gazzaniga P, Manso L, Zamarchi R, de Lascoiti AF, De Mattos-Arruda L, Ignatiadis M, Lebofsky R, van Laere SJ, Meier-Stiegen F, Sandri MT, Vidal-Martinez J, Politaki E, Consoli F, Bottini A, Diaz-Rubio E, Krell J, Dawson SJ, Raimondi C, Rutten A, Janni W, Munzone E, Caranana V, Agelaki S, Almici C, Dirix L, Solomayer EF, Zorzino L, Johannes H, Reis-Filho JS, Pantel K, Pierga JY, Michiels S. (2014). Clinical validity of circulating tumour cells in patients with metastatic breast cancer: a pooled analysis of individual patient data. The lancet oncology. 15(4):406–14. PMID: 24636208.

85. Cristofanilli M, Pierga JY, Reuben J, Rademaker A, Davis AA, Peeters DJ, Fehm T, Nole F, Gisbert-Criado R, Mavroudis D, Grisanti S, Giuliano M, Garcia-Saenz JA, Stebbing J, Caldas C, Gazzaniga P, Manso L, Zamarchi R, de Lascoiti AF, De Mattos-Arruda L, Ignatiadis M, Cabel L, van Laere SJ, Meier-Stiegen F, Sandri MT, Vidal-Martinez J, Politaki E, Consoli F, Generali D, Cappelletti MR, Diaz-Rubio E, Krell J, Dawson SJ, Raimondi C, Rutten A, Janni W, Munzone E, Caranana V, Agelaki S, Almici C, Dirix L, Solomayer EF, Zorzino L, Darrigues L, Reis-Filho JS, Gerratana L, Michiels S, Bidard FC, Pantel K. (2019). The clinical use of circulating tumor cells (CTCs) enumeration for staging of metastatic breast cancer (MBC): International expert consensus paper. Critical reviews in oncology/hematology. 134:39–45. PMID: 30771872.

86. Anampa J, Xue X, Oh S, Kornblum N, Sadan S, Oktay M, Condeelis J, Sparano J. Abstract P6-18-22: Phase Ib study of rebastinib plus antitubulin therapy with paclitaxel (P) or eribulin (E) in patients with HER2-negative metastatic breast cancer (MBC). AACR; 2019.

87. Padua D, Zhang XH, Wang Q, Nadal C, Gerald WL, Gomis RR, Massague J. (2008). TGFbeta primes breast tumors for lung metastasis seeding through angiopoietin-like 4. Cell. 133(1):66–77. PMID: 18394990 / PMCID: PMCPMC2390892.

88. Saxe JP, Tomilin A, Scholer HR, Plath K, Huang J. (2009). Post-translational regulation of Oct4 transcriptional activity. PLoS One. 4(2):e4467. PMID: 19221599 / PMCID: PMCPMC2637973.

89. Niwa H, Miyazaki J, Smith AG. (2000). Quantitative expression of Oct-3/4 defines differentiation, dedifferentiation or self-renewal of ES cells. Nat Genet. 24(4):372–6. PMID: 10742100.

90. Keysar SB, Le PN, Miller B, Jackson BC, Eagles JR, Nieto C, Kim J, Tang B, Glogowska MJ, Morton JJ, Padilla-Just N, Gomez K, Warnock E, Reisinger J, Arcaroli JJ, Messersmith WA, Wakefield LM, Gao D, Tan AC, Serracino H, Vasiliou V, Roop DR, Wang XJ, Jimeno A. (2017). Regulation of Head and Neck Squamous Cancer Stem Cells by PI3K and SOX2. Journal of the National Cancer Institute. 109(1). PMID: 27634934 / PMCID: PMCPMC5025278.

91. He C, Danes JM, Hart PC, Zhu Y, Huang Y, de Abreu AL, O’Brien J, Mathison AJ, Tang B, Frasor JM, Wakefield LM, Ganini D, Stauder E, Zielonka J, Gantner BN, Urrutia RA, Gius D, Bonini MG. (2019). SOD2 acetylation on lysine 68 promotes stem cell reprogramming in breast cancer. Proc Natl Acad Sci U S A. 116(47):23534–41. PMID: 31591207 / PMCID: PMCPMC6876149.

92. Esposito M, Mondal N, Greco TM, Wei Y, Spadazzi C, Lin SC, Zheng H, Cheung C, Magnani JL, Lin SH, Cristea IM, Sackstein R, Kang Y. (2019). Bone vascular niche E-selectin induces mesenchymal-epithelial transition and Wnt activation in cancer cells to promote bone metastasis. Nature cell biology. 21(5):627–39. PMID: 30988423 / PMCID: PMCPMC6556210.

93. Gao W, Wu D, Wang Y, Wang Z, Zou C, Dai Y, Ng CF, Teoh JY, Chan FL. (2018). Development of a novel and economical agar-based non-adherent three-dimensional culture method for enrichment of cancer stem-like cells. Stem Cell Res Ther. 9(1):243. PMID: 30257704 / PMCID: PMCPMC6158801.

94. Ramchandani D, Lee SK, Yomtoubian S, Han MS, Tung CH, Mittal V. (2019). Nanoparticle Delivery of miR-708 Mimetic Impairs Breast Cancer Metastasis. Molecular Cancer Therapeutics. 18(3):579–91. PMID: 30679387 / PMCID: PMCPMC6532393.

95. Gorodetska I, Lukiyanchuk V, Peitzsch C, Kozeretska I, Dubrovska A. (2019). BRCA1 and EZH2 cooperate in regulation of prostate cancer stem cell phenotype. Int J Cancer. 145(11):2974–85. PMID: 30968962.

96. De Lope C, Martin-Alonso S, Auzmendi-Iriarte J, Escudero C, Mulet I, Larrasa-Alonso J, Lopez-Antona I, Matheu A, Palmero I. (2019). SIX1 represses senescence and promotes SOX2-mediated cellular plasticity during tumorigenesis. Sci Rep. 9(1):1412. PMID: 30723235 / PMCID: PMCPMC6363751.

97. Gedye C, Sirskyj D, Lobo NC, Meens J, Hyatt E, Robinette M, Fleshner N, Hamilton RJ, Kulkarni G, Zlotta A, Evans A, Finelli A, Jewett MA, Ailles LE. (2016). Cancer stem cells are underestimated by standard experimental methods in clear cell renal cell carcinoma. Sci Rep. 6:25220. PMID: 27121191 / PMCID: PMCPMC4848484.

98. Entenberg D, Pastoriza JM, Oktay MH, Voiculescu S, Wang Y, Sosa MS, Aguirre-Ghiso J, Condeelis J. (2017). Time-lapsed, large-volume, high-resolution intravital imaging for tissue-wide analysis of single cell dynamics. Methods. 128:65–77. PMID: 28911733 / PMCID: PMC5659295.

99. Entenberg D, Wyckoff J, Gligorijevic B, Roussos ET, Verkhusha VV, Pollard JW, Condeelis J. (2011). Setup and use of a two-laser multiphoton microscope for multichannel intravital fluorescence imaging. Nat Protoc. 6(10):1500–20. PMID: 21959234 / PMCID: PMC4028841.

100. Sharma VP. (2018). ImageJ plugin HyperStackReg V5.6. Zenodo.

101. Tinevez JY, Perry N, Schindelin J, Hoopes GM, Reynolds GD, Laplantine E, Bednarek SY, Shorte SL, Eliceiri KW. (2017). TrackMate: An open and extensible platform for single-particle tracking. Methods. 115:80–90. PMID: 27713081.

102. Murphy DA, Courtneidge SA. (2011). The ‘ins’ and ‘outs’ of podosomes and invadopodia: characteristics, formation and function. Nat Rev Mol Cell Biol. 12(7):413–26. PMID: 21697900 / PMCID: PMC3423958.

103. Shah MM, Zerlin M, Li BY, Herzog TJ, Kitajewski JK, Wright JD. (2013). The role of Notch and gamma-secretase inhibition in an ovarian cancer model. Anticancer research. 33(3):801–8. PMID: 23482747 / PMCID: PMCPMC3893696.

104. Milano J, McKay J, Dagenais C, Foster-Brown L, Pognan F, Gadient R, Jacobs RT, Zacco A, Greenberg B, Ciaccio PJ. (2004). Modulation of notch processing by gamma-secretase inhibitors causes intestinal goblet cell metaplasia and induction of genes known to specify gut secretory lineage differentiation. Toxicol Sci. 82(1):341–58. PMID: 15319485.

105. Sjolund J, Johansson M, Manna S, Norin C, Pietras A, Beckman S, Nilsson E, Ljungberg B, Axelson H. (2008). Suppression of renal cell carcinoma growth by inhibition of Notch signaling in vitro and in vivo. The Journal of clinical investigation. 118(1):217–28. PMID: 18079963 / PMCID: PMCPMC2129233.

106. Symmans WF, Ayers M, Clark EA, Stec J, Hess KR, Sneige N, Buchholz TA, Krishnamurthy S, Ibrahim NK, Buzdar AU, Theriault RL, Rosales MF, Thomas ES, Gwyn KM, Green MC, Syed AR, Hortobagyi GN, Pusztai L. (2003). Total RNA yield and microarray gene expression profiles from fine-needle aspiration biopsy and core-needle biopsy samples of breast carcinoma. Cancer. 97(12):2960–71. PMID: 12784330.

107. Gupta PK, Baloch ZW. (2002). Intraoperative and on-site cytopathology consultation: utilization, limitations, and value. Semin Diagn Pathol. 19(4):227–36. PMID: 12469790.

108. As F, Ma Z. Fine needle aspiration biopsy cytology of breast: a diagnostic approach based on pattern recognition. In: Practical Cytopathology: A Diagnostic Approach to Fine Needle Aspiration Biopsy. Practical Cytopathology: A Diagnostic Approach to Fine Needle Aspiration Biopsy: Elsevier; 2017.

109. Kryczek I, Liu S, Roh M, Vatan L, Szeliga W, Wei S, Banerjee M, Mao Y, Kotarski J, Wicha MS, Liu R, Zou W. (2012). Expression of aldehyde dehydrogenase and CD133 defines ovarian cancer stem cells. Int J Cancer. 130(1):29–39. PMID: 21480217 / PMCID: PMCPMC3164893.

110. Stone OJ, Pankow N, Liu B, Sharma VP, Eddy RJ, Wang H, Putz AT, Teets FD, Kuhlman B, Condeelis JS, Hahn KM. (2019). Optogenetic control of cofilin and alphaTAT in living cells using Z-lock. Nat Chem Biol. 15(12):1183–90. PMID: 31740825 / PMCID: PMCPMC6873228.

